# Inhibition of VP2-mediated entry: a potential antiviral strategy to treat or prevent calicivirus disease

**DOI:** 10.64898/2026.05.10.723473

**Authors:** Charlotte B. Lewis, Lee Sherry, Rochelle McGrory, Eibhlin Gould, Millie Freitag, Hagar Sasvari, June Southall, Filip Petrov, Amit Meir, Nithya Govindan, Andrea Spiri, Matteo Bordicchia, Regina Hofmann-Lehmann, Vanessa R. Barrs, Joe Grove, Andrew G. Jamieson, Margaret J. Hosie, David Bhella

## Abstract

The Caliciviridae include many notable human and animal pathogens, including norovirus and sapovirus, which cause outbreaks of acute gastroenteritis. We previously demonstrated that, following receptor engagement, feline calicivirus (FCV) assembles a portal structure at a unique capsid three-fold axis. This comprises twelve copies of the minor capsid protein VP2 and is essential for genome delivery. We designed a short peptide based on our structure data that occludes the VP2 binding sites on the capsid surface, to prevent assembly of the VP2 portal and thereby halt the viral entry mechanism. Incubation with low micromolar concentrations of the peptide considerably reduced the infectivity of two laboratory strains and four clinical isolates of FCV associated with respiratory or virulent-systemic disease. Cryo-electron microscopy structures of FCV virions complexed with the peptide confirmed that the peptide occupies the VP2 binding site on the major capsid protein VP1, preventing portal assembly and subsequent genome delivery. Our data show that targeting VP2 is a viable antiviral approach to preventing calicivirus infection, with potential for the treatment or prevention of norovirus disease.

## Introduction

Caliciviridae are small, non-enveloped, positive-sense RNA viruses. The family includes many pathogens of medical and veterinary importance. The most notorious human-infecting caliciviruses are noroviruses and sapoviruses, which cause outbreaks of acute gastroenteritis, particularly in hospitals, care homes, schools and hospitality settings (Patel *et al*., 2008; Becker-Dreps *et al*., 2020). Norovirus in particular poses a significant economic and healthcare burden, with norovirus-associated gastroenteritis in England alone estimated to cost the National Health Service over £100 million in healthcare costs alongside a societal burden of £297 million (Lopman *et al*., 2004; Sandmann *et al*., 2018). Globally, norovirus has been estimated to cause over $4 billion in healthcare costs per year, with particularly high costs associated with disease in children and older adults (Bartsch *et al*. 2016).

Difficulty propagating norovirus in cell culture has presented a substantial barrier to the development of direct acting antivirals. Much of our understanding of calicivirus biology has therefore been established through the investigation of animal infecting caliciviruses. One well-studied and tractable model virus is feline calicivirus (FCV), which can be readily grown to high titres *in vitro*. FCV causes respiratory disease in cats and was first isolated in New Zealand in the 1950s (Fastier, 1957). Infection is usually associated with oral ulceration, fever, lethargy, and, in rare cases lameness and pneumonia (Radford *et al*., 2007; Berger *et al*., 2015). A clinically distinct form of FCV, termed virulent systemic (VS) FCV, has subsequently emerged (Pedersen *et al*., 2000; Hurley *et al*., 2004; Foley *et al*., 2006; Bordicchia *et al*., 2021); VS FCV infection is often fatal and is associated with high morbidity and mortality, even in vaccinated cats (Hurley, 2005; Bordicchia *et al*., 2021; Park *et al*., 2024). Therefore, FCV is an important veterinary pathogen as well as an established *in vitro* surrogate for the study of human norovirus. FCV is especially well suited to studies of the viral entry pathway since its proteinaceous receptor, feline junctional adhesion molecule A (fJAM-A), has been identified (Makino *et al*., 2006).

Caliciviruses such as FCV package their small (~7 kb) positive-sense RNA genomes in a T=3 icosahedral capsid that is ~40 nm in diameter (Chen *et al*., 2006). The capsid consists of 180 copies of the ~60kDa major capsid protein VP1. These are assembled into 90 large arch shaped dimeric capsomeres, termed AB and CC dimers owing to the position of their constituent VP1 protomers in the icosahedral asymmetric unit (Figure 1A). AB dimers are arranged about the icosahedral five-fold symmetry axes, while CC dimers are located at the icosahedral two-fold axes. VP1 is subdivided into three domains – the N-terminal arm (NTA) at the capsid interior, the shell (S) domain that makes up the contiguous floor of the capsid, and the protruding (P) domain that forms the distinctive spikes. The P-domain is further divided into the P1 and P2 sub-domains. The distal P2 sub-domain bears the receptor binding site and major immunodominant epitopes. In addition to the major capsid protein, virions also incorporate the minor capsid protein VP2, which is critical for infectivity across several caliciviruses (Sosnovtsev *et al*., 2005; Wei *et al*., 2008; Ishiyama *et al*., 2024). We previously showed by cryogenic electron microscopy that FCV VP2 forms a portal assembly at a unique capsid three-fold symmetry axis following receptor engagement (Conley *et al*., 2019). The funnel-shaped portal comprises twelve copies of VP2 in which its long N-terminal α-helix (residues 18-63 - helix a) forms an α-helical barrel that rises from the capsid surface (Figure 1B). VP2 adopts two alternating conformations arranged about the unique icosahedral three-fold symmetry axis. Conformer 1 folds into the lumen of the barrel while conformer 2 anchors the portal to the capsid surface, binding to the neighbouring VP1 P-domains (Figure 1B, 1C). The distal N-terminal amino-acid residues were not resolved in the cryo-EM structure but are largely hydrophobic.

**Figure 1.**
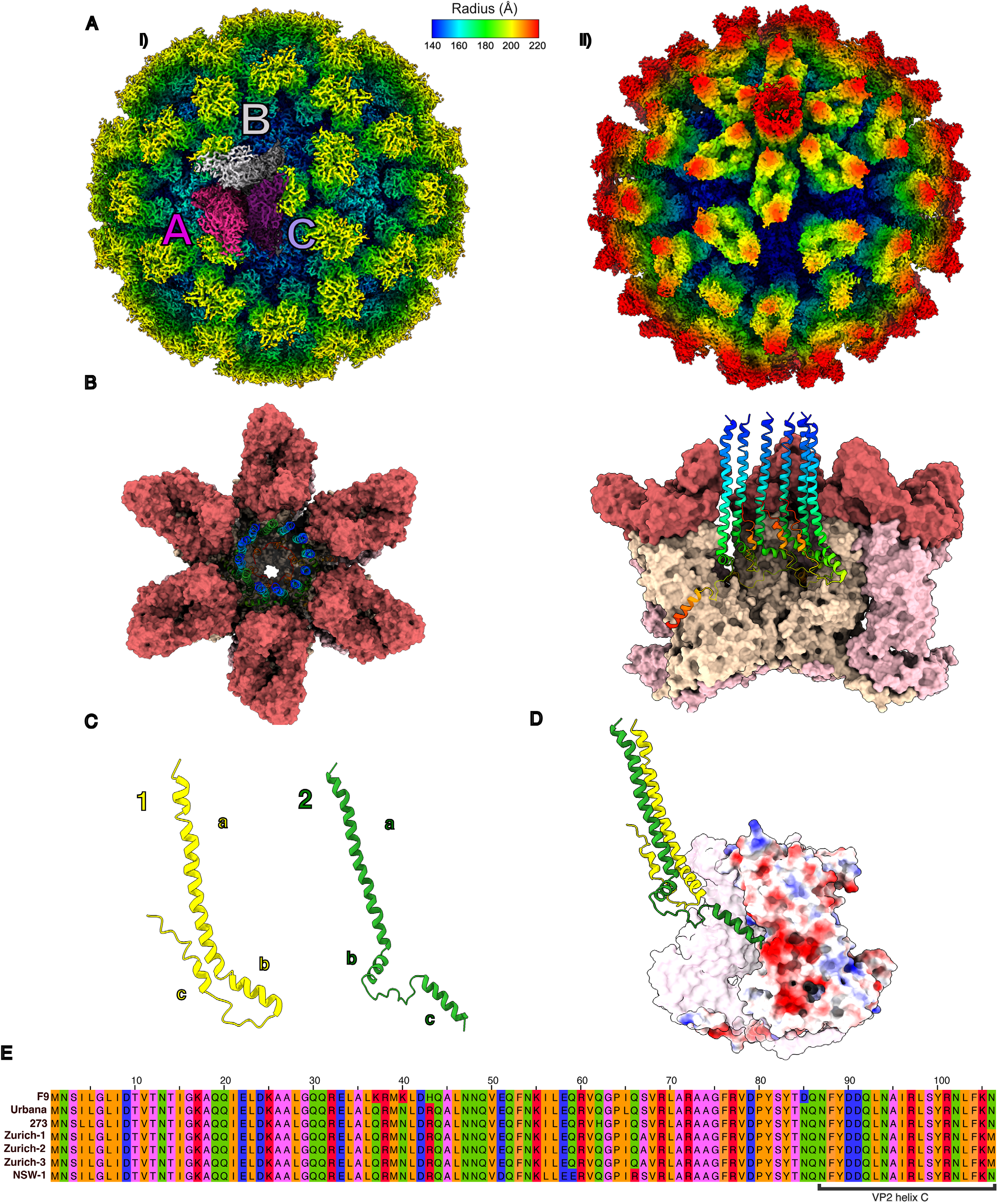
(A) (i) Cryo-EM structure of FCV F9 coloured according to radius from centre. A, B, and C quasi-equivalent positions of VP1 are coloured pink, grey, and purple respectively (Conley *et al*., 2019; EMD-0054). (ii) Cryo-EM structure of FCV F9 bound to fJAM-A, with a portal-like assembly comprising 12 copies of VP2 present at a unique three-fold axis (Conley *et al*., 2019; EMD-0056). (B) Top-down (left) and side view (right) of the VP2 portal, illustrating six of the twelve conformers. VP2 monomers are illustrated as rainbow ribbons, with N-terminus in blue and C-terminus in red. (C) Illustration of the two distinct conformers in the portal structure. (D) Electrostatic diagram illustrating the interaction of conformer 2 of VP2 (green) with a negatively charged cleft of VP1. Negatively charged regions are indicated in red and positively charged regions are indicated in blue. (E) Amino acid sequence alignment of VP2 from the FCV strains used in this study. The 20 C-terminal residues of VP2 are highlighted with a square bracket. Residues are coloured according to biochemical properties. Pink: non-polar (Gly, Ala, Ser, Thr); orange: hydrophobic (Cys, Val, Ile, Leu, Pro, Phe, Tyr, Met, Trp); green: polar (Asn, Gln, His); blue: negatively charged (Asp, Glu); red: positively charged (Lys, Arg). Sequences were aligned and visualised in Jalview 2.11.4.0 (Procter *et al*., 2021).

Following receptor engagement, the capsid undergoes several structural rearrangements to enable portal formation. The dimeric P-domain spikes become more mobile, rotating and tilting towards the capsid icosahedral three-fold axes. At the site of portal assembly, structural rearrangements in the P-domains create two clefts into which conformer 2 of VP2 binds. These six copies of VP2, which are highly conserved at both a structural and sequence level across FCV isolates, anchor the portal to the capsid surface via two α-helices at residues 64-72 (helix b) and 87-106 (helix c) that bind to the P2 and P1 domains respectively (Conley *et al*., 2019) (Figure 1D, 1E). VP2 was shown to be essential for delivery of the genomic RNA to the cytosol (Sun *et al*., 2024). Therefore, we hypothesise that the portal assembly mediates genome delivery by the insertion of the hydrophobic N-terminal distal tips of VP2 into the endosomal membrane, either forming a direct bridge between the capsid interior and the cytosol, or by disrupting the endosomal membrane to enable genome delivery upon capsid disassembly.

Here, we demonstrate that a 24 amino-acid residue peptide based on the C-terminal sequence of VP2 (helix c), targets the VP1-VP2 interface and substantially inhibits FCV infection. This peptide, termed VP2_87-106_, demonstrated antiviral activity against representative VS and acute respiratory clinical isolates of FCV at low micromolar concentrations. Cryo-EM structures of FCV isolates complexed with the peptide revealed that the peptide occupies the VP2 binding cleft in the P1 sub-domain of VP1, suggesting that its mode of action is via disruption of the assembly of the portal structure. As VP2 is highly conserved and essential for infectivity across many caliciviruses, this demonstrates an attractive antiviral strategy with potential to target other members of the calicivirus family, including human norovirus.

## Materials & Methods

### Cells and virus strains

Feline embryo-A (FEA) (Jarrett *et al*., 1973) and Crandell-Reese feline kidney (CRFK) (Crandell *et al*., 1973) cells were maintained in Dulbecco’s minimum essential medium, supplemented with 10% heat-inactivated foetal bovine serum and 1% penicillin-streptomycin (Gibco) at 37°C in an atmosphere of 5% CO_2_.

The vaccine strain F9 (Bittle *et al*., 1976) and reference strain Urbana (Sosnovstev *et al*., 1995) were used, alongside the classical respiratory clinical isolate 273 (Spiri *et al*., 2019), and the virulent systemic clinical isolates NSW-1 (Bordicchia *et al*., 2021), Zurich-1, Zurich-2, and Zurich-3 (Willi *et al*., 2016). FCV 273 was isolated from a 13-year-old privately-owned domestic male cat with diabetes mellitus, which had presented with chronic gingivitis, stomatitis, and lingual ulcerations for approximately one year. NSW-1 originated from an unvaccinated 6-week-old stray kitten presenting with upper respiratory signs and lameness, with disease duration lasting 2-4 days. Zurich-1 was obtained from a 12.5-year-old, regularly vaccinated female cat presenting with tongue lesions and anorexia, and Zurich-2 from a 4.9-year-old unvaccinated male cat presenting with tongue lesions. Both animals subsequently recovered after treatment. Zurich-3 was isolated from an 8.9-year-old vaccinated male cat with severe stomatitis and glossitis as well as tongue ulceration that died four days after presentation to clinic.

### Peptide design

VP2_87-106_ and VP2_87-106 scramble_ were synthesised by Peptide Synthetics and reconstituted from the lyophilised form using distilled water. VP2_87-106_ was designed with an identical sequence to the 20 C-terminal residues of VP2 from FCV strain F9 with an additional C-terminal poly-lysine motif inserted to increase solubility. VP2_87-106 scramble_ was designed by entering the VP2_87-106_sequence (excluding the poly-lysine motif) into the Peptide Nexus website (https://peptidenexus.com/article/sequence-scrambler).

Sequence of VP2_87-106_ : H-NFYDDQLNAIRLSYRNLFKNKKKK-OH

Sequence of VP2_87-106 scramble_: H-RLNNNFRIDKYSLNFQLDAYKKKK-OH

Peptides for alanine scanning mutagenesis were synthesised using a Syro II peptide synthesiser on a 0.02 mmol scale. The peptides were synthesised using Rink Amide functionalised aminomethyl resin (loading 0.53 mmol/g), employing a SPPS Fmoc/tBu protecting group strategy. Resin was swelled at 75 °C for 15 min in DMF. Peptides were elongated in cycles of amino acid coupling followed by Fmoc removal. Coupling reactions were performed using Fmoc-amino acid (5 equiv., 0.5 M in DMF), DIC (5 equiv., 0.5 M in DMF) and Oxyma Pure (5 equiv., 1 M in DMF). Reaction mixtures were stirred at room temperature for 1 h. This procedure was then repeated with fresh reagents at room temperature for a further 1 h. Following coupling, the resin was washed with DMF (6 × 0.6 mL). Fmoc removal was carried out using 40% piperidine in DMF with 5% formic acid (0.5 mL, v/v/v) at room temperature for 3 min followed by 20% piperidine in DMF with 5% formic acid (0.5 mL, v/v/v) at room temperature for 12 min. Following deprotection, the resin was washed with DMF (6 × 0.5 mL). Resin-bound peptides were washed with dichloromethane (5 × 1 mL) prior to peptide cleavage. For further experimental details and peptide sequences, see Supplementary Methods.

### Virus Propagation and Purification

FEA cells were seeded to a density of approximately 0.2×10^6^cells/well in a 24-well plate (Corning) and incubated overnight at 37°C, 5% CO_2_. Viral stocks were generated by infecting cells with FCV strains 273, F9, Urbana, NSW-1, Zurich-1, Zurich-2, and Zurich-3, diluted to a multiplicity of infection (MOI) of 0.1 in serum-free (SF) DMEM (Thermo Scientific). When assessing peptide efficacy, diluted viral stocks were treated beforehand with 25μM – 200nM of peptide and incubated for 1h at 37°C and 5% CO_2_ before being inoculated onto cells. Culture fluids were harvested at 16-20 hours post-in-fection to allow for multiple rounds of viral replication (Abente *et al*., 2010), centrifuged at 5000 × g for 5 minutes to remove cell debris, and stored at −80°C until use.

To prepare high titre virus stocks for cryo-EM analyses, FEA cells were seeded to a density of approximately 5×10^6^ cells/flask in 24 T175 flasks (Nunc) and incubated overnight. Cells were then infected with FCV strains 273 and NSW-1 at an MOI of 0.01 and maintained at 37°C, 5% CO_2_ until complete cytopathic effect (CPE) was observed. Following complete CPE, culture fluids were harvested and centrifuged at 5,000 × g for 10 minutes in a Heraeus Megafuge 16R centrifuge (Thermo Scientific) using a TX-400 swinging bucket rotor (Thermo Scientific) to remove cell debris. Supernatant was then treated with 8% w/v of PEG-8000 (Sigma-Aldrich) and 200 mM NaCl (VWR) and incubated overnight at 4°C to precipitate proteins. Precipitated protein was centrifuged at 5000 × g for 30 minutes at 4°C, the resulting pellet was resuspended in 25ml PBS and centrifuged at 5000 × g for 20 minutes and then the virus was pelleted by centrifugation (Sorvall Discovery 90SE, Thermo Scientific) through a 30% (w/v) sucrose cushion at 151,000 × g for 3.5h at 4°C using a SureSpin 630 swinging bucket rotor (Thermo Scientific). The resulting virus pellet was resuspended in a PBS + 1% NP-40 + 0.5% sodium deoxycholate solution, centrifuged at 10,000 × g for 10 minutes at 4°C to remove remaining insoluble material, and supernatant was then centrifuged through a 15%-60% w/v sucrose gradient (TH-641 swinging bucket rotor, Thermo Scientific) at 28,000 × g for 16h at 4°C. The resulting gradient was collected into 1 mL fractions and analysed by SDS-PAGE to identify virus-containing fractions at ~63kDa (the approximate weight of the major viral capsid protein VP1). Positive fractions were pelleted by centrifugation at 20,000 × rpm for 3h at 4°C (TH-641 swinging bucket rotor). The resulting pellet was then resuspended in 100μl PBS and stored at 4°C until use.

### Plaque-forming assay

CRFK cells were seeded in 12-well plates to an approximate cell density of 0.5×10^6^ cells/well and incubated overnight at 37°C, 5% CO_2_. Previously generated viral stocks were then serially diluted tenfold in SF DMEM to concentrations ranging from 10^−1^ to 10^−6^ before 500μl of inoculum were added to cells. Cells were then incubated for 1h at 37°C. After 1h, wells were overlaid with a solution of 2.4% Avicel (Sigma-Aldrich) mixed 1:1 with SF DMEM, and plates were incubated at 37°C for 28-32 hours. Following incubation, cells were fixed with 4% formaldehyde (Sigma-Aldrich, diluted in PBS) for 30 minutes. Avicel and media were then removed and plates were washed with PBS and stained with Coomassie Blue solution. Infectious viral titres were expressed in plaque-forming units/ml (pfu/ml).

### Nanoluciferase-encoding reporter virus neutralisation assays

CRFK cells were seeded in white plates (Revvity) to a density of approximately 0.04 × 10^6^ cells/well 24 hours before infection. On the day of infection, peptide VP2_87-106_ was diluted to the appropriate concentrations in SF DMEM before being mixed 1:1 with 2000pfu/ml of NanoLuc-Urbana (also diluted in SF DMEM). The mixture was then incubated at 37°C and 5% CO_2_ for 1 hour. 100μl of the mixture was added to the pre-seeded CRFK cells for a final MOI of 100 PFU/well. Cells and virus were then incubated for 16 hours at 37°C and 5% CO_2_. Cells were then treated with the Nano-Glo® Luciferase Assay System (Promega) according to the manufacturer’s instructions, before luminescence was measured with a Revvity Ensight multimode plate reader.

### Circular dichroism spectroscopy

Circular dichroism experiments were performed using a Jasco J-180 spectrophotometer at the Neil Bulleid Integrated Protein Analysis facility (University of Glasgow). Peptides were diluted to 0.5mg/ml in ultra-pure water and scans were acquired in the far-UV region (195-250nm) using a pathlength of 0.2mm. Buffer reference CD spectra were subtracted and data were corrected for concentration and path length. Signals were smoothed using means-movement with a convolution width of 21.

### Cryo-EM sample preparation

Quantifoil 2/2 Cu300 mesh grids were cleaned with acetone and glow-discharged at 35mA for 45 seconds using an Emitech K100X Glow Discharge System. Three microlitres of each purified virus sample were then added to grids held in an FEI Mark IV Vitrobot (at 100% humidity and 4°C) and blotted for 3 seconds with a blot force of 3 to remove excess liquid. Grids were then plunge-frozen into liquid-nitrogen-cooled liquid ethane and stored in liquid nitrogen prior to imaging.

### Cryo-EM imaging

Grids were imaged in a JEOL Cryo-ARM 300 equipped with a Direct Electron Apollo detector at the Scottish Centre for Macromolecular Imaging. Data were collected at a nominal magnification of 60kx, corresponding to a pixel size of 1.04 Å/pixel with the energy filter slit width set to 20 eV and a dose of ~50 e/Å^2^. Micrograph movies were captured in super-resolution mode as 2s exposures and saved at a frame rate of 20 frames/s. Automated data collection was performed using SerialEM (David N. Mastronarde, 2003).

### Cryo-EM data processing

RELION 5.0.0 was used to process micrograph movies (Scheres 2012). Motion correction was performed using RELION’s own implemention of MotionCor2, and CTF estimated using CTFFIND-4.1 (Rohou and Griogrieff, 2015). Particles were selected from micrographs using a Topaz auto-picking model trained on previous FCV datasets. Particles were then used as input for multiple rounds of reference-free 2D classification; 100 class averages were generated during early rounds, which were then filtered down to 20 class averages. Classes with sharp, high-resolution features were selected for subsequent processing. One round of 3D classification (2 class averages) was performed using a previously generated FCV structure as a reference map, with the class that achieved the highest resolution selected for 3D refinement with I2 icosahedral symmetry imposed. The resulting map was then refined through repeated rounds of CTF refinement, 3D refinement, and post-processing until the structures converged. To resolve the structure of CC-dimers, where they deviated from strict icosahedral symmetry, symmetry expansion followed by focussed classification with a cylindrical mask was used to resolve the tilted P-dimer, as previously described (Conley, 2019).

### Focused classification

Particles from the global map were re-extracted and Fourier cropped by a factor of 6. Extracted particles were then subjected to symmetry expansion with icosahedral (I2) symmetry imposed, using relion_symmetry_expand. A cylindrical mask covering the P-dimer was used to exclude all areas of the particle except the region of interest and was resampled onto the virus reference map in ChimeraX (Goddard *et al*. 2018). RELION was used to apply a soft edge to the mask, and 3D classification of the symmetry-expanded particles into 10 class averages was then performed with no symmetry imposed (C1) and a Tau value of 20.

### Atomic model building

Upon convergence, the post-processed, B-factor sharpened map of each virion was submitted to ModelAngelo for automated model building, along with files containing the sequence of NSW-1 or 273 as appropriate alongside the sequence of VP2_87-106_. This built sequence chains into the map density, with the final model output as a .cif file. Two chains for each AB or CC subunit, as well as additional peptide chains, were selected for further refinement and all other chains of the model were deleted, resulting in a model of eight chains for 273+VP2_87-106_ and six chains for NSW-1+VP2_87-106_. ISOLDE (Croll, 2018) was used to relax the model into the cryo-EM density map using an AMBER forcefield simulation at a temperature of 20K. The positions and orientations of each residue in the model were refined to ensure correct rotamer orientation and phi-psi torsion angles, as well as to resolve steric clashes between atoms, using the cryo-EM density map as reference. The temperature was then lowered to 0K and a further round of per-residue refinement was conducted before one round of Phenix real-space refinement (Adams *et al*., 2010) was performed to generate the final map.

## Results

### Peptide VP2_87-106_ blocks infection of multiple FCV strains in a sequence-specific manner

To explore the potential for calicivirus entry inhibition by targeting the VP2-VP1 interface, we devised a peptide based on the C-terminal 20 residues of FCV strain F9 VP2, termed VP2_87-106_. To improve solubility, we added 4 lysine residues at the C-terminus. We have previously showed that in the native structure of the dodecameric VP2 portal, residues 87-106 of VP2 form an α-helix that binds into a negatively charged cleft formed by the P1-domains of both AB- and CC-dimers arranged about the portal-vertex, anchoring the VP2 portal to the capsid surface.

To test the efficacy of VP2_87-106_ against FCV, clinical isolates NSW-1, Zurich 1-3, 273 and reference strains F9 and Urbana were diluted to an MOI of 0.1 in SF DMEM and VP2_87-106_ was added to a final concentration of 200nM-25μM. The virus-peptide mixtures were either pre-incubated for 1 hour at 37°C before being inoculated onto FEA cells, or inoculated onto cells immediately after mixing. The infections were then incubated for 1 hour, after which media were removed and replaced with fresh SF DMEM. Culture fluids were harvested after 16-20 hours and titrated by plaque assay using CRFK cells (Figure 2A).

**Figure 2.**
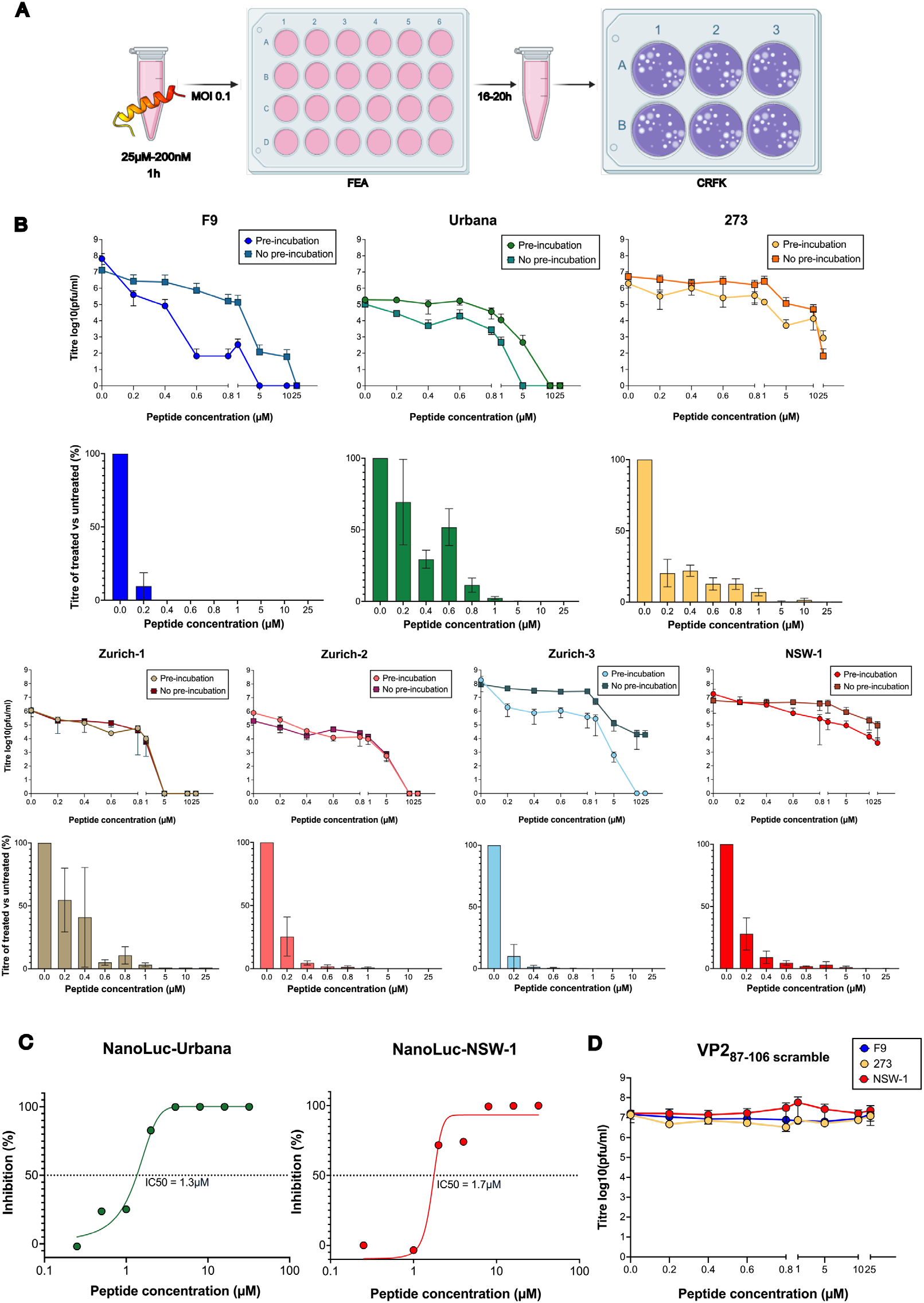
(A). Schematic illustrating protocol for FCV stock generation and plaque assays to assess peptide efficacy. (B).Titre of FCV strains pre-incubated with VP2_87-106_ (circles) or added to cells together with VP2_87-106_ (squares). Experiments were performed independently in triplicate. Error bars: mean + SEM. (C) Concentration-response curve of NanoLuc-Urbana and NanoLuc-NSW-1 treated with 250nM-32μM of peptide, with IC50 indicated by a dotted line. Experiments were performed independently in triplicate. (D) Titres of F9 (blue circle), 273 (orange circle), and NSW-1 (red circle) upon treatment with concentrations of peptide VP2_87-106 scramble_ ranging from 200nM-25μM. Experiments were performed independently in triplicate. Error bars: mean + SEM.

We found that VP2_87-106_ reduced FCV infectivity in a dose dependent manner for all isolates tested.In all cases, pre-treatment with 25μM of VP2_87-106_ resulted in a decrease of infectivity of 3x log^10^ or greater. A decrease in infectivity of of 100-1000 fold was observed consistently across most isolates upon pre-treatment with 5μM of peptide, and titres across all viruses were reduced by over 50% upon pre-treatment with 800nM of peptide. Concurrent addition of both virus and peptide to cells similarly inhibited infection with all isolates tested (Figure 2B).

Overall, pre-incubating virus with VP2_87-106_ resulted in a superior antiviral activity against four of the seven isolates tested (F9, Urbana, Zurich-3, and NSW-1). This suggests that VP2_87-106_ might access the binding cleft in VP1 prior to receptor engagement, potentially because this site is transiently exposed by the innate capsid plasticity of FCV P-domains (Lewis *et al*., 2025).

To determine the half-maximal inhibitory concentration of VP2_87-106_, a nanoluciferase-encoding FCV reporter system was used. Briefly, NanoLuc® luciferase was inserted into the leader of the capsid (LC) protein in FCV Urbana, as this region was shown previously to tolerate insertions (Abente *et al*., 2010). This allows for sensitive luciferase-based quantification of infectious virions. NanoLuc-Urbana was pre-incubated for 1 hour with peptide (concentrations ranging from 250nM-32μM). The virus-peptide mixtures were then inoculated onto CRFK cells and incubated for 16 hours before cells were lysed for infection readout. To measure the IC50 of VP2_87-106_ against a VS-FCV strain, a second reporter virus was created in which the Urbana VP1 sequence was replaced with that of NSW-1. The IC50 values of VP2_87-106_ measured against NanoLuc-Urbana and NanoLuc-NSW-1 were 1.3μM and 1.7μM respectively (Figure 2C).

To investigate the sequence specificity of peptide VP2_87-106_, a scrambled peptide VP2_87-106 scramble_ was devised by randomly shuffling the original VP2_87-106_ peptide sequence, with the exception of the four N-terminal lysines, which were retained to preserve solubility. VP2_87-106 scramble_ did not cause any reduction in infectivity when tested against several FCV strains, suggesting that the antiviral activity of VP2_87-106_ is specific to its sequence (Figure 2D). The secondary structures of peptides VP2_87-106_ and VP2_87-106 scramble_ were assessed using far-UV circular dichroism spectroscopy (CD) in H_2_O showing that VP2_87-106_ retains an α-helical secondary structure in solution, whereas VP2_87-106 scramble_ exhibited a random coil conformation (Supplemental Figure 1).

Overall, these results highlighted a dose-dependent susceptibility of multiple field and clinical strains of FCV to VP2_87-106_. It should be noted that the strains tested appeared to fall into two groups; F9, Urbana, and Zurich-1 appeared more susceptible to VP2_87-106_, being fully neutralised at concentrations of 5μM upon pre-incubation with peptide. Comparatively, 273, Zurich-3, and NSW-1 appeared to be less susceptible, requiring pre-treatment with ≥10μM of peptide for full neutralisation in plaque reduction assays. Differences in the neutralisation susceptibility of vaccine strains such as F9 compared to clinical isolates has been previously noted by Mcdonagh *et al*. (2015) in the context of mefloquine treatment, although in that case mefloquine exhibited reduced potency against F9 compared to field strains. Further differences in susceptibility of field strains to inactivation by biocides was observed by Di Martino *et al*. (2010), highlighting the importance of testing antiviral treatments across a range of clinically relevant viruses.

### VP2_87-106_ interacts with the FCV capsid at the VP1-VP2 interface

To confirm that VP2_87-106_ inhibits FCV entry by blocking the VP2 binding site in the P1 sub-domain of VP1, we used cryo-EM to calculate the structures of VP2_87-106_ in complex with the respiratory clinical isolate FCV 273 and the virulent systemic clinical isolate NSW-1 in the presence or absence of a soluble ectodomain fragment of the receptor fJAM-A. In brief, virions were purified and incubated with a molar excess of fJAM-A and VP2_87-106_ or with VP2_87-106_ alone. The virus-receptor-peptide complexes were prepared for cryo-EM imaging by plunge-freezing in liquid ethane and imaged on a JEOL CryoARM 300 equipped with a Direct Electron Apollo detector at the Scottish Centre for Macromolecular Imaging.

A cryo-EM map of FCV 273 in complex with VP2_87-106_ and soluble fJAM-A was calculated, with the imposition of icosahedral symmetry to a resolution of 2.8Å. Cryo-EM maps of 273 and NSW-1 in complex with VP2_87-106_ alone were calculated to a resolution of 2.7Å and 2.5Å respectively. In all structures, clear density corresponding to VP2_87-106_ was seen to occupy the VP2 binding cleft on the P1 domain of VP1 at both AB and CC dimer positions across the entire icosahedral virion (Figure 3). To clearly resolve the CC-dimer - VP2_87-106_ complex it was necessary to apply symmetry expansion and focussed classification, as the CC-dimers were found to tilt away from the icosahedral two-fold symmetry axes, leading to incoherent averaging of the P-domain density at this site. This deviation from icosahedral symmetry closely resembles that which we have previously described in structures of FCV bound to fJAM-A (Bhella and Goodfellow, 2011; Conley *et al*., 2019).

**Figure 3.**
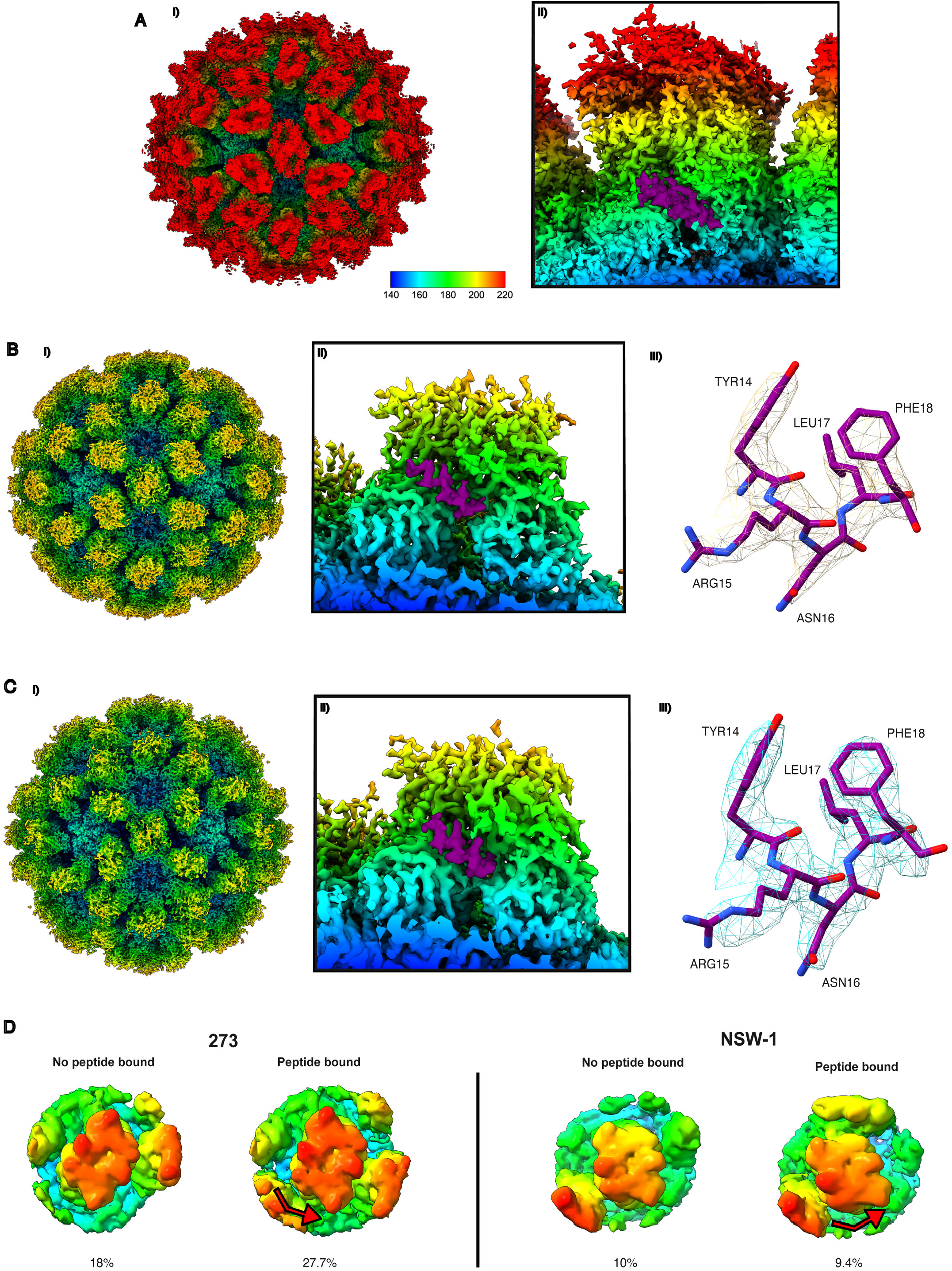
A) i) Cryo-EM structure of FCV 273 incubated with peptide VP2_87-106_ and fJAM-A. Map is radially coloured according to distance from centre. ii) Section through the structure described in A) i) displaying an A/B P-dimer with VP2_87-106_ present at the P1 domain (twisted density, coloured purple). B) i) Cryo-EM structure of FCV 273 incubated with peptide VP2_87-106_ only. Note the absence of density corresponding to fJAM-A (shaded red in Figure 3A). ii) Section through the structure described in B) i) displaying an A/B P-dimer with VP2_87-106_ present at the P1 domain (twisted density, coloured purple). C) i) Cryo-EM structure of FCV NSW-1 incubated with peptide VP2_87-106_ only. ii) Section through the structure described in C) i) displaying an A/B P-dimer with VP2_87-106_ present at the P1 domain (twisted density, coloured purple). iii) Representative density of VP2_87-106_ in the structure described, with residue positions noted. D) Representative classes from focused classification of AB P-dimers of 273 and NSW-1 in the absence (left) and presence (right) of peptide, with red arrows indicating direction of movement. The proportion of total particles that each class represents is indicated in percentages.

The finding that VP2_87-106_ occupies the VP1 P1 binding cleft in the absence of fJAM-A was surprising. We had anticipated that receptor engagement would be necessary to induce the substantial conformational changes required for portal assembly. To accommodate VP2 helix c, the two P1 domains of a VP1 dimer move together to form the binding cleft. This rearrangement occurs when P dimers move to a raised state following binding of fJAM-A. Our data suggest that binding of VP2_87-106_ prompts the same VP1 tilting and/or rotation towards the three-fold axis, as well as transition to the raised conformation that we previously observed to be a consequence of fJAM-A engagement, leading to portal assembly (Figure 3BII/3CII). (Conley *et al*., 2019, extended data). In prior cryo-EM maps of FCV decorated with soluble ectodomain fragments of fJAM-A, P-domains exhibited considerable flexibility and were not well-resolved at sites that were not proximal to the portal. At the portal axis, however, adjacent P-dimers were well resolved. We attributed this finding to the stabilising effect of VP2 which tightly binds the P-dimers to the dodecameric portal complex, constraining their movement (Bhella and Goodfellow, 2011; Conley *et al*., 2019). Similarly, the raised/tilted conformations of AB and CC P-dimers adopted upon VP2_87-106_ engagement appear to be considerably more constrained than those observed upon fJAM-A engagement alone. We also observed an opening at the three-fold pore in the capsid shell at the site of VP2_87-106_ engagement consistent with that observed during portal assembly (Conley *et al*., 2019), suggesting that the binding of VP2_87-106_ is sufficient to prompt equivalent conformational rearrangements in both the protruding and shell domains.

The observation that VP2_87-106_ binds FCV virions in the absence of soluble receptor is consistent with the increased inhibitory activity of VP2_87-106_ upon pre-incubation with some of the FCV isolates tested. The inherent flexibility of FCV P-dimers might transiently expose the VP1-VP2 binding cleft, allowing VP2_87-106_ to saturate this site before the virus encounters the host cell surface and engages the cellular receptor fJAM-A.

To identify the proportion of AB and CC P-domains occupied by peptide, symmetry expansion followed by focused 3D classification with a cylindrical mask was applied. Briefly, each particle was assigned 60 orientations corresponding to those of the 60 asymmetric units present in icosahedral objects; a cylindrical mask was then placed over the P-dimers, such that further refinement would only be focused on this region. 3D classification of P-dimers into 10 classes was then performed without symmetry imposition, allowing for identification of structural heterogeneity. When isolate 273 was incubated with VP2_87-106_, two of ten 3D classes representing AB-dimers consisted of poorly-re-solved densities of insufficient quality for further analysis (‘junk’ classes); however, these represented a small overall proportion (6.3%) of total particles. Unambiguous peptide density was present in six of the seven remaining classes, which represented the majority of particles (73.3%) (Supplemental Figure 3). For CC-dimers, two of the ten 3D classes were insufficiently resolved, representing 3.5% of total particles. Unambiguous peptide density was present in seven of the eight remaining classes, representing 90.5% of CC-dimers. This discrepancy in peptide occupancy between AB and CC-dimers could be due to differences in flexibility between the two P-dimer types. CC-dimers, similarly to those observed in previous FCV structures (Bhella *et al*., 2008) appeared to be more poorly resolved and therefore might exhibit a greater degree of flexibility, facilitating access to the binding cleft. For those classes in which peptide was bound, peptide density was observed on both sides of the P-dimers in all cases.

Similar analysis of isolate NSW-1 incubated with VP2_87-106_ showed peptide density present in eight of ten classes corresponding to AB-dimers, and nine of ten classes corresponding to CC-dimers, representing 83.6% and 79.5% occupancy respectively (Supplemental Figure 3). All AB-classes and four of ten CC-classes had peptide unambiguously bound to one side of the P-dimers only; the remaining peptide-bound CC-classes had peptide bound to both sides. It should be noted that in one AB-class and one CC-class (representing 7.6% and 14.4% of particles respectively), a weak sausage-shaped density potentially representing VP2_87-106_ could be observed on both sides; however, this was only visible at very low thresholds. As discussed above, binding of peptide induced conformational shifts of P-dimers towards the three-fold axis, similarly to that observed upon assembly of the VP2 portal structure (Figure 3D; Conley *et al*., 2019).

In both cases, VP2_87-106_ formed two hydrogen bonds with A and C subunits, and 1-2 hydrogen bonds with B and D subunits (Figure 4A, 4B). In NSW-1, an additional hydrogen bond is formed between AB-bound VP2_87-106_ and subunit D (ASN8 −> GLU555; not shown). In both models, the N-terminal poly-lysine motif was not resolved, suggesting it does not contribute to peptide interactions with VP1 and instead exists in an unbound or mobile conformation. To evaluate essential residues required for VP2_87-106_ activity, alanine scan mutagenesis was performed at positions 1-20 across the native peptide sequence. Residues at which an alanine was already present were not modified. Nano-Luc-Urbana was incubated for one hour with two-fold serial dilutions of each edited peptide at concentrations ranging from 250nM to 32uM, before being inoculated onto CRFK cells and incubated for 16-18 hours. Cells were then lysed for infection readout and luminescence quantified.

**Figure 4.**
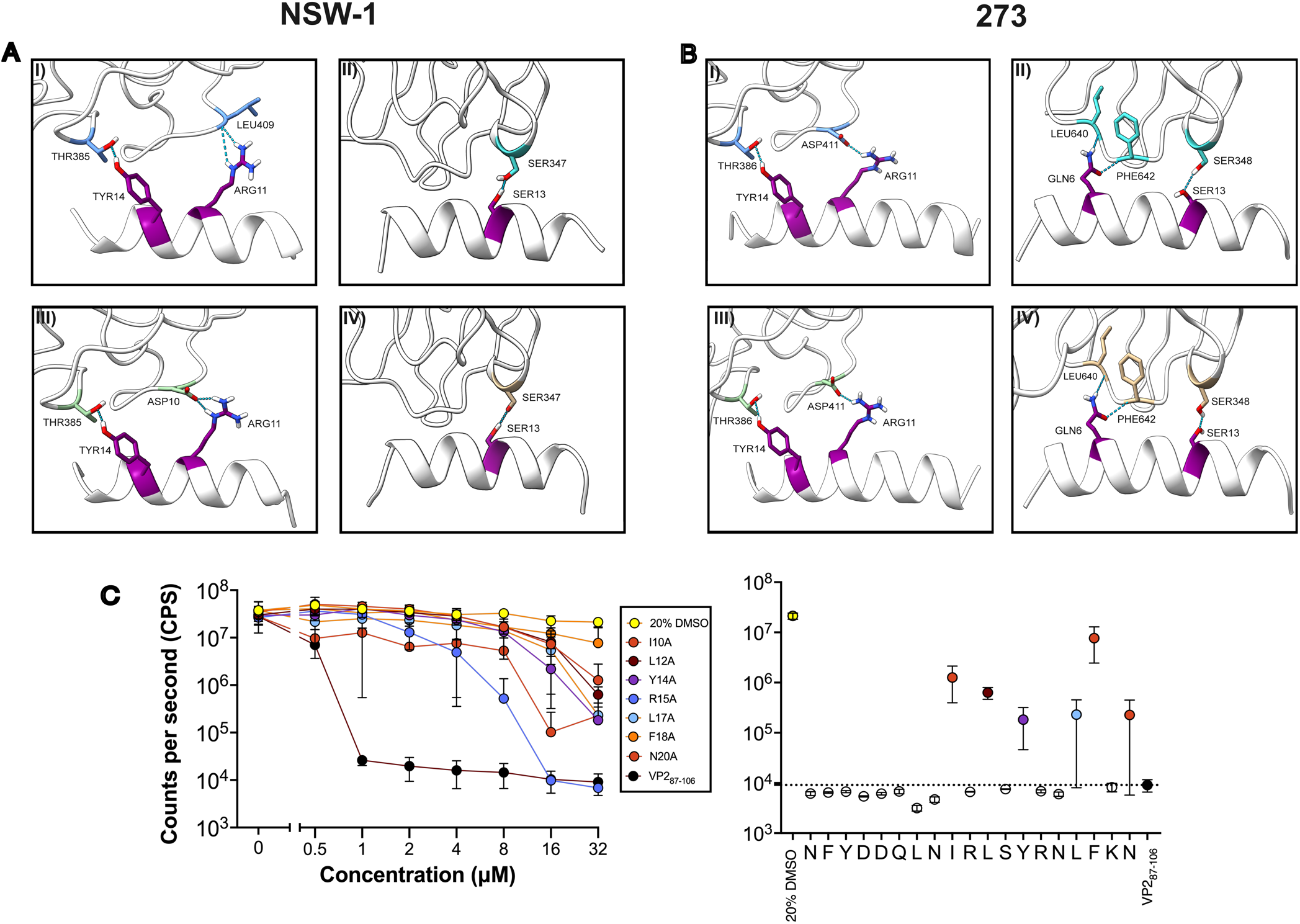
Ribbon diagrams indicating VP1-VP2_87-106_ interactions in NSW-1 (A) and 273 (B) with representative examples of interactions between peptide and A (I), B (II), C (III) and D (IV) VP1 subunits shown. Interacting residues are shown as sticks alongside residue identities and positions. Hydrogen bonds between residues, identified using the ‘H-bonds’ tool in ChimeraX, are indicated by a dashed blue line. C) (left) Concentration-response curve of NanoLuc-Urbana treated with 250nM-32μM of peptide analogs as well as native peptide (VP2_87-106_) and 20% DMSO controls indicated in the legend. For clarity, only those mutants that demonstrated reduced inhibitory activity against NanoLuc-Urbana are shown. (Right) titre of NanoLuc-Urbana upon treatment with 32μM of the respective peptide. The residue which has been substituted for an alanine is highlighted along the X-axis. The dashed line indicates titre of NanoLuc-Urbana upon treatment with 32μM of VP2_87-106_. Error bars = mean + SEM.

From this analysis, seven mutants were identified that exhibited reduced activity against Nano-Luc-Urbana (I10A, L12A, Y14A, R15A, L17A, F18A, N20A), suggesting that residues at these positions might affect antiviral potency by contributing to the VP1-VP2_87-106_ interface (Figure 4C). It was notable that Y14 consistently interfaced directly with VP1 in both our models and that of Conley *et al*. (2019), whereas other mutants such as I10A and L12A were directly adjacent to interfacing residues (such as R11 in our model), suggesting that mutagenesis might impact peptide secondary structure in a manner that disrupts binding of adjacent residues. Taken together, these data suggest the most crucial residues for VP1-VP2_87-106_ interactions, regardless of FCV isolate, are located in the C-terminal half of the peptide, and that truncation of the first eight N-terminal residues may not impact antiviral activity.

## Discussion

Human norovirus is the leading cause of non-bacterial gastroenteritis worldwide, causing ~700 million cases and >200,000 deaths annually, including ~70,000 children under the age of 5. Infant mortality is highest in low- and middle-income countries, while mortality in elderly populations is more widespread. The societal and economic costs are substantial, being estimated at ~$97B per year (Bartsch *et al*., 2020; Standaert *et al*., 2008). Therefore, there is an urgent need for antivirals targeting calicivirus disease. FCV remains a significant veterinary concern due to the lack of clinically approved antiviral treatment options, particularly in the face of VS-FCV outbreaks.

This study aimed to characterise the antiviral activity of a novel peptide VP2_87-106_. We designed this peptide to target the FCV entry pathway by inhibiting assembly of the VP2 portal structure that is necessary for delivery of the calicivirus genome into the host cell. VP2_87-106_ exhibited potent *in vitro* antiviral activity against seven clinical isolates of FCV in the low micromolar range, indicating the potential utility of VP2-targeting antivirals as a treatment against calicivirus infection. Cryo-EM structures of VP2_87-106_ in complex with the respiratory isolate 273 and virulent systemic isolate NSW-1 indicate that the peptide occludes the VP2 helix c interaction site formed by conformational rearrangements of VP1 P-dimers, in the P1-subdomain. The mode of action of this peptide is therefore most likely that which we intended, preventing or destabilising the assembly of the VP2 portal complex and thereby preventing delivery of the viral genome to the host cytosol.

The VP2 protein is highly conserved and has been shown to be essential in many caliciviruses (Sosnovtsev *et al*., 2005; Wei *et al*., 2008; Ishiyama *et al*., 2024). The structure and function of VP2 is well understood for FCV due to its tractability in the laboratory and prior knowledge of the protein receptor, which has enabled detailed investigations of the entry pathway by ourselves and others. VP2 is also critical for Tulane virus entry (Wei *et al*., 2008) and appears to play a significant role in norovirus capsid stability, where it might associate with the interior of the shell domain of VP1 (Vonpungsawad *et al*., 2013; Lin *et al*., 2014). However, the role of VP2 is less understood in the context of human norovirus, owing to its poor cultivability *in vitro*. Nonetheless, it is likely that VP2 assembles a portal structure in human norovirus and other caliciviruses of interest. AlphaFold 3 predictions of VP2 across a variety of caliciviruses predict the formation of a large α-helical barrel, similar to that observed in FCV (Abramson *et al*., 2024; Supplemental Figure 4). Unfortunately, limitations on current AlphaFold 3 performance prevent inference of the structure bound to the capsid surface.

Our strategy of targeting the VP2-VP1 interface with a peptide derived from the sequence of native VP2 is attractive owing to the high degree of conservation of VP2 and its interaction sites on VP1 (Supplemental Figure 5). We hypothesise that there is less potential for escape mutations to arise as any mutations in VP1 that prevent peptide binding would require compensatory mutations in VP2.

In summary, our data show that targeting assembly of the VP2 portal by occluding assembly sites on the capsid surface is a viable antiviral strategy for FCV. VP2 is found in all caliciviruses and has been shown to be essential for infectivity in many of them. Structure predictions suggest that VP2 from other caliciviruses, including human norovirus and the widely studied laboratory model murine norovirus, likely forms portal assemblies to mediate genome delivery. Therefore, designing antiviral peptides, peptidomimetics or small molecules that target or occlude VP2-VP1 interaction sites is a promising strategy with potential to be extended to Caliciviridae that cause major health and economic impacts in human populations.

## Author Contributions

Conceptualization CBL, DB, JG, LS; Methodology AGJ, CBL, DB, JG, LS, RM; Validation AM, CBL, FP; Formal analysis CBL; Investigation CBL, DB, EG, JS, MF, RM; Data curation CBL, DB; Resources AS, HS, MB, NG, RH, VB; Writing—original draft preparation, CBL, DB; Writing—review and editing, AM, DB, CBL, LS, MJH; Visualization, CBL, DB.; Supervision, DB, JG, LS, MJH; Project administration, DB; Funding acquisition, DB, MJH.

All authors have read and agreed to the published version of the manuscript.

## Funding acknowledgement

This work was supported by the Biotechnology and Biological Sciences Research Council (BB/T002239/1), to DB and MJH, Medical Research Council (MC_UU_00034/1) to DB. CL and FP are the recipients of MRC PhD studentships (MC_ST_00034). HS is supported by the PetPlan Charitable Trust (S24-1326-1365). EG was supported by a Royal Society of Chemistry-funded summer internship. We acknowledge the Scottish Centre for Macromolecular Imaging (SCMI) for access to cryo-EM instrumentation, funded by the MRC (MC_PC_17135, MC_UU_00034/7, MR/X011879/1) and SFC (H17007).

## Data Availability Statement

Raw data micrograph movies are deposited in EMPIAR (https://www.ebi.ac.uk/empiar/). Cryo-EM maps are deposited in the EM-databank (https://www.ebi.ac.uk/emdb/). Atomic models are deposited in the protein-databank (https://www.ebi.ac.uk/pdbe/). Accession numbers: FCV 273 + VP2_87-106_-EMPIAR-13515, EMD-57695, PDB 30FJ. FCV 273 + fJAM + VP2_87-106_-EMPI-AR-13522, EMD-57687. FCV NSW-1 + VP2_87-106_: EMPIAR-13523, EMD-57678, PDB 30EX

## Acknowledgments

We thank James Streetley for support with cryo-EM data collection.

## References

Abente, E. J., Sosnovtsev, S. V., Bok, K. & Green, K. Y. Visualization of feline calicivirus replication in real-time with recombinant viruses engineered to express fluorescent reporter proteins. Virology 400, 18–31 (2010).

Abramson, J. et al. Accurate structure prediction of biomolecular interactions with AlphaFold 3. Nature 636, E4–E4 (2024).

Adams, P. D. et al. PHENIX : a comprehensive Python-based system for macromolecular structure solution. Acta Crystallogr D Biol Crystallogr 66, 213–221 (2010).

Bartsch, S. M., Lopman, B. A., Ozawa, S., Hall, A. J. & Lee, B. Y. Global Economic Burden of Norovirus Gastroenteritis. PLoS One 11, e0151219 (2016).

Bartsch, S. M., O’Shea, K. J. & Lee, B. Y. The Clinical and Economic Burden of Norovirus Gastroenteritis in the United States. J Infect Dis 222, 1910–1919 (2020).

Becker-Dreps, S., González, F. & Bucardo, F. Sapovirus: an emerging cause of childhood diarrhea. Curr Opin Infect Dis 33, 388–397 (2020).

Berger, A. et al. Feline calicivirus and other respiratory pathogens in cats with Feline calicivirus-related symptoms and in clinically healthy cats in Switzerland. BMC Vet Res 11, 282 (2015).

Bhella, D., Gatherer, D., Chaudhry, Y., Pink, R. & Goodfellow, I. G. Structural insights into calicivirus attachment and uncoating. J Virol 82, 8051–8058 (2008).

Bhella, D. & Goodfellow, I. G. The cryo-electron microscopy structure of feline calicivirus bound to junctional adhesion molecule A at 9-angstrom resolution reveals receptor-induced flexibility and two distinct conformational changes in the capsid protein VP1. J Virol 85, 11381–11390 (2011).

Bittle, J. L. & Rubic, W. J. Immunization against feline calicivirus infection. Am J Vet Res 37, 275–278 (1976).

Bordicchia, M. et al. Feline Calicivirus Virulent Systemic Disease: Clinical Epidemiology, Analysis of Viral Isolates and In Vitro Efficacy of Novel Antivirals in Australian Outbreaks. Viruses 13, 2040 (2021).

Chen, R., Neill, J. D., Estes, M. K. & Prasad, B. V. V. X-ray structure of a native calicivirus: Structural insights into antigenic diversity and host specificity. Proc. Natl. Acad. Sci. U.S.A. 103, 8048–8053 (2006).

Conley, M. J. et al. Calicivirus VP2 forms a portal-like assembly following receptor engagement. Nature 565, 377–381 (2019).

Crandell, R. A., Fabricant, C. G. & Nelson-Rees, W. A. Development, characterization, and viral susceptibility of a feline (Felis catus) renal cell line (CRFK). In Vitro 9, 176–185 (1973).

Croll, T. I. ISOLDE: a physically realistic environment for model building into low-resolution electron-density maps. Acta Crystallogr D Struct Biol 74, 519–530 (2018).

Di Martino, B., Ceci, C., Di Profio, F. & Marsilio, F. In vitro inactivation of feline calicivirus (FCV) by chemical disinfectants: resistance variation among field strains. Arch Virol 155, 2047–2051 (2010).

Fastier, L. B. A new feline virus isolated in tissue culture. Am J Vet Res 18, 382–389 (1957).

Foley, J., Hurley, K., Pesavento, P. A., Poland, A. & Pedersen, N. C. Virulent systemic feline calicivirus infection: Local cytokine modulation and contribution of viral mutants. Journal of Feline Medicine and Surgery 8, 55–61 (2006).

Glaser, F. et al. ConSurf: identification of functional regions in proteins by surface-mapping of phylogenetic information. Bioinformatics 19, 163–164 (2003).

Goddard, T. D. et al. UCSF ChimeraX: Meeting modern challenges in visualization and analysis. Protein Science 27, 14–25 (2018).

Hurley, K. F. et al. An outbreak of virulent systemic feline calicivirus disease. javma 224, 241–249 (2004).

Hurley, K. F. Virulent Systemic Feline Calicivirus: Recognition and Control. https://www.cabidigitallibrary.org/doi/pdf/10.5555/20053197418.

Ishiyama, R. et al. Production of infectious reporter murine norovirus by VP2 trans-complementation. J Virol 98, e01261–23 (2024).

Jarrett, O., Laird, H. M. & Hay, D. Determinants of the Host Range of Feline Leukaemia Viruses. Journal of General Virology 20, 169–175 (1973).

Lewis, C. B. et al. Conformational Flexibility in Capsids Encoded by the Caliciviridae. Viruses 16, 1835 (2024).

Lin, Y., Fengling, L., Lianzhu, W., Yuxiu, Z. & Yanhua, J. Function of VP2 protein in the stability of the secondary structure of virus-like particles of genogroup II norovirus at different pH levels: Function of VP2 protein in the stability of NoV VLPs. J Microbiol. 52, 970–975 (2014).

Lopman, B. A. et al. Epidemiology and cost of nosocomial gastroenteritis, Avon, England, 2002-2003. Emerg Infect Dis 10, 1827–1834 (2004).

Makino, A. et al. Junctional adhesion molecule 1 is a functional receptor for feline calicivirus. J Virol 80, 4482–4490 (2006).

Mastronarde, D. N. SerialEM: A Program for Automated Tilt Series Acquisition on Tecnai Microscopes Using Prediction of Specimen Position. Microsc Microanal 9, 1182–1183 (2003).

McDonagh, P., Sheehy, P. A., Fawcett, A. & Norris, J. M. Antiviral effect of mefloquine on feline calicivirus in vitro. Veter-inary Microbiology 176, 370–377 (2015).

Park, J., Lee, D., Hong, Y.-J., Hwang, C.-Y. & Hyun, J.-E. Outbreaks of nosocomial feline calicivirus-associated virulent systemic disease in Korea. J Vet Sci 25, e51 (2024).

Patel, M. M. et al. Systematic literature review of role of noroviruses in sporadic gastroenteritis. Emerg Infect Dis 14, 1224–1231 (2008).

Pedersen, N. C., Elliott, J. B., Glasgow, A., Poland, A. & Keel, K. An isolated epizootic of hemorrhagic-like fever in cats caused by a novel and highly virulent strain of feline calicivirus. Veterinary Microbiology 73, 281–300 (2000).

Procter, J. B. et al./person-group>. Alignment of Biological Sequences with Jalview. in Multiple Sequence Alignment (ed. Katoh, K.) vol. 2231 203–224 (Springer US, New York, NY, 2021).

Radford, A. D., Coyne, K. P., Dawson, S., Porter, C. J. & Gaskell, R. M. Feline calicivirus. Vet Res 38, 319–335 (2007).

Rohou, A. & Grigorieff, N. CTFFIND4: Fast and accurate defocus estimation from electron micrographs. Journal of Structural Biology 192, 216–221 (2015).

Sandmann, F. G. et al. Estimating the Hospital Burden of Norovirus-Associated Gastroenteritis in England and Its Opportunity Costs for Nonadmitted Patients. Clin Infect Dis 67, 693–700 (2018).

Scheres, S. H. W. RELION: Implementation of a Bayesian approach to cryo-EM structure determination. Journal of Structural Biology 180, 519–530 (2012).

Sosnovtsev, S. V., Belliot, G., Chang, K.-O., Onwudiwe, O. & Green, K. Y. Feline Calicivirus VP2 Is Essential for the Production of Infectious Virions. J Virol 79, 4012–4024 (2005).

Sosnovtsev, S. & Green, K. Y. RNA transcripts derived from a cloned full-length copy of the feline calicivirus genome do not require VpG for infectivity. Virology 210, 383–390 (1995).

Spiri, A. M. et al. Environmental Contamination and Hygienic Measures After Feline Calicivirus Field Strain Infections of Cats in a Research Facility. Viruses 11, 958 (2019).

Sun, W. et al. VP2 mediates the release of the feline calicivirus RNA genome by puncturing the endosome membrane of infected cells. J Virol 98, e00350–24 (2024).

Vongpunsawad, S., Venkataram Prasad, B. V. & Estes, M. K. Norwalk Virus Minor Capsid Protein VP2 Associates within the VP1 Shell Domain. J Virol 87, 4818–4825 (2013).

Wei, C., Farkas, T., Sestak, K. & Jiang, X. Recovery of Infectious Virus by Transfection of In Vitro-Generated RNA from Tulane Calicivirus cDNA. J Virol 82, 11429–11436 (2008).

Willi, B. et al. Molecular characterization and virus neutralization patterns of severe, non-epizootic forms of feline calicivirus infections resembling virulent systemic disease in cats in Switzerland and in Liechtenstein. Vet Microbiol 182, 202–212 (2016).

