## Supplemental information for "Inhibition of VP2-mediated entry: a potential antiviral strategy to treat or prevent calicivirus disease"

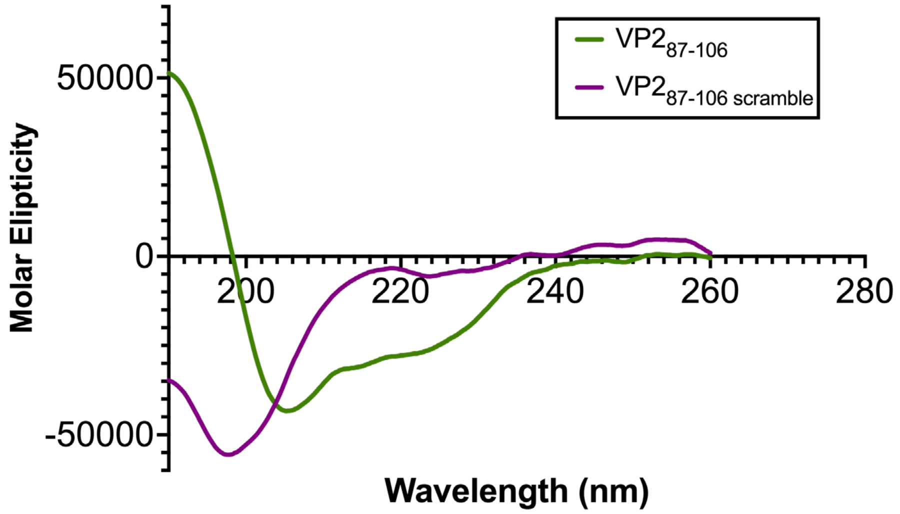


**Supplemental Figure 1.** Circular dichroism (CD) spectra of VP2$_{87\text{-}106}$ and VP2$_{87\text{-}106\ \text{scramble}}$ at a concentration of 0.5mg/ml in ultrapure water..

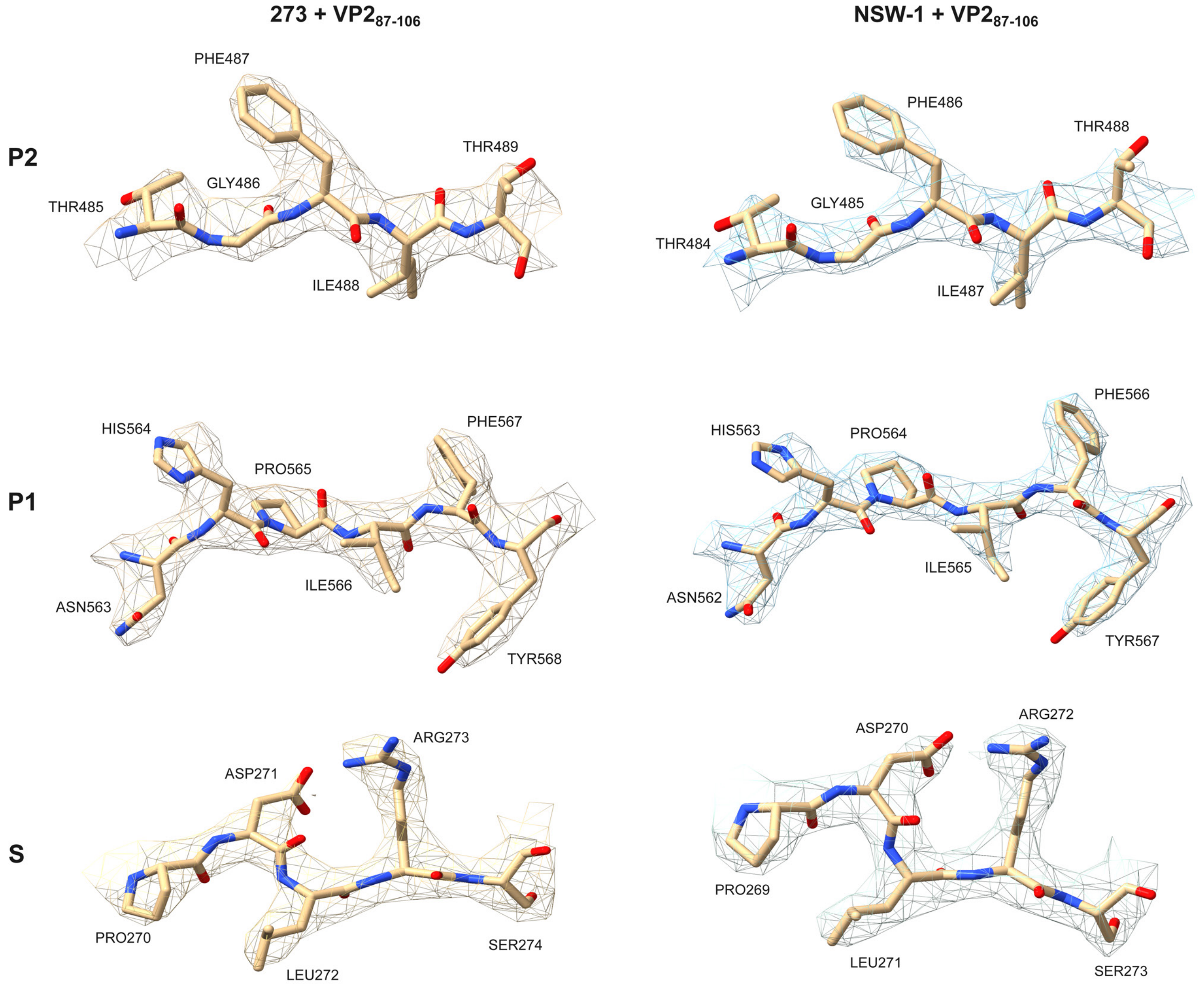


**Supplemental Figure 2.** Representative densities and models for sections of the P2, P1, and S domains in cryo-EM maps calculated for FCV 273 + VP2$_{87\text{-}106}$ (left) and NSW-E1 + VP2$_{87\text{-}106}$ (right), with residue positions noted.

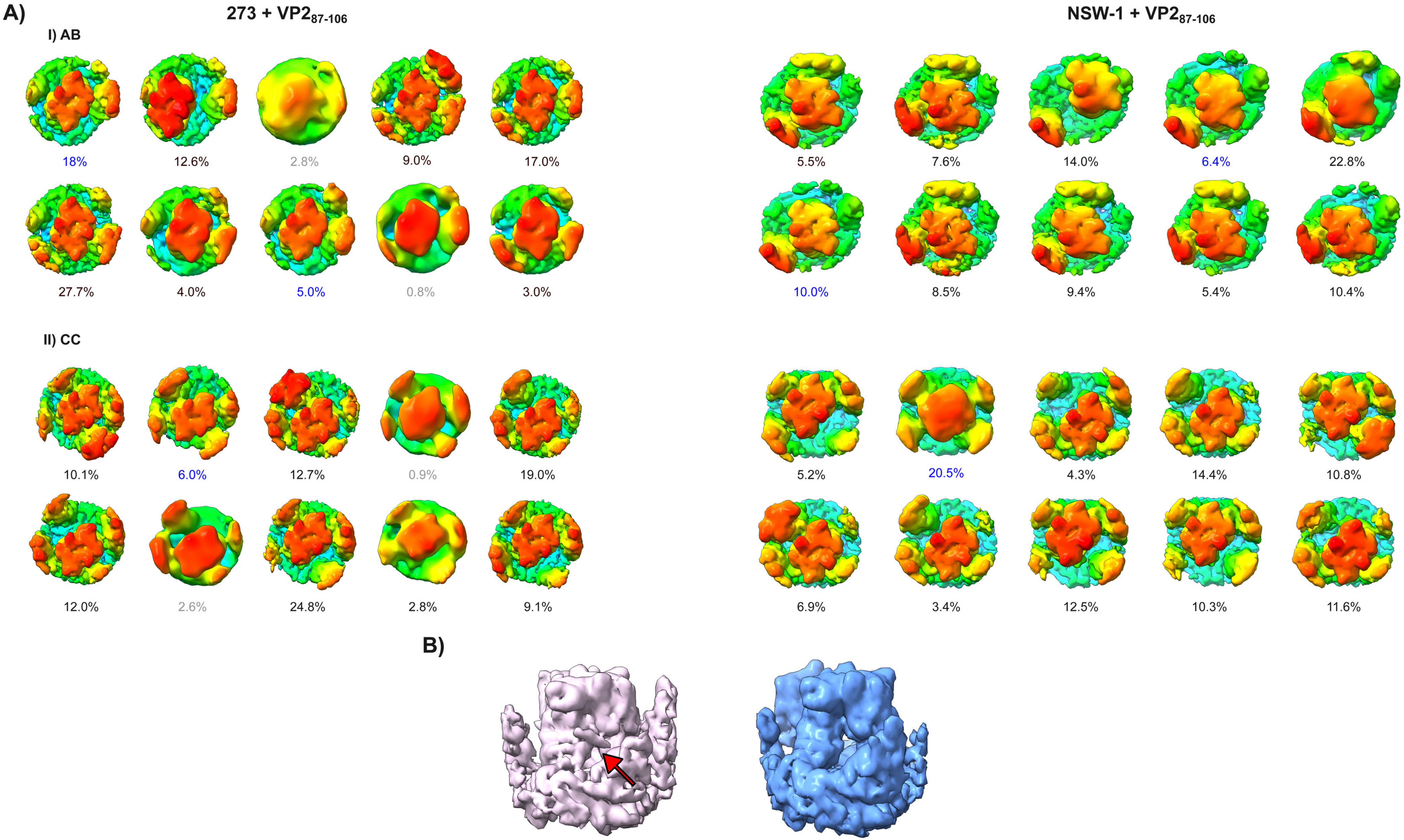


**Supplemental Figure 3.** A) All focused classes of I) AB and II) CC P-dimers 273 + VP2$_{87\text{-}106}$ (left) and NSW-E1 + VP2$_{87\text{-}106}$ (right). The proportion of the total number of particles for each class is indicated as a percentage. Text is coloured dark blue in cases where no peptide was bound. In cases where the density was insufficient to determine peptide binding, the text is coloured gray. B) Example of focused classes with peptide bound (left) and no peptide bound (right), with red arrow indicating the sausage-shaped density representing VP287-106.

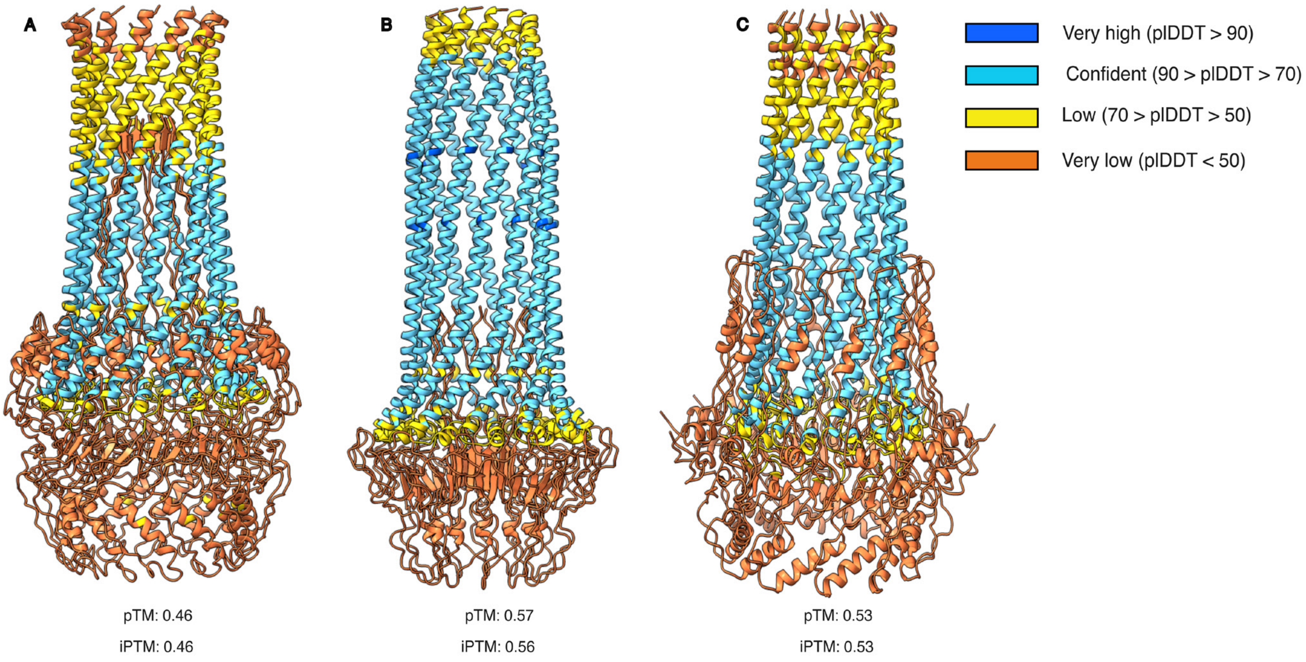


**Supplemental Figure 4.** Alphafold 3 predictions of VP2 dodecamers for human norovirus (A), murine norovirus (B) and Tulane virus (C), coloured according to per-residue prediction confidence scores (pLDDT). Prediction confidence score ranges and corresponding colours are indicated in the key.

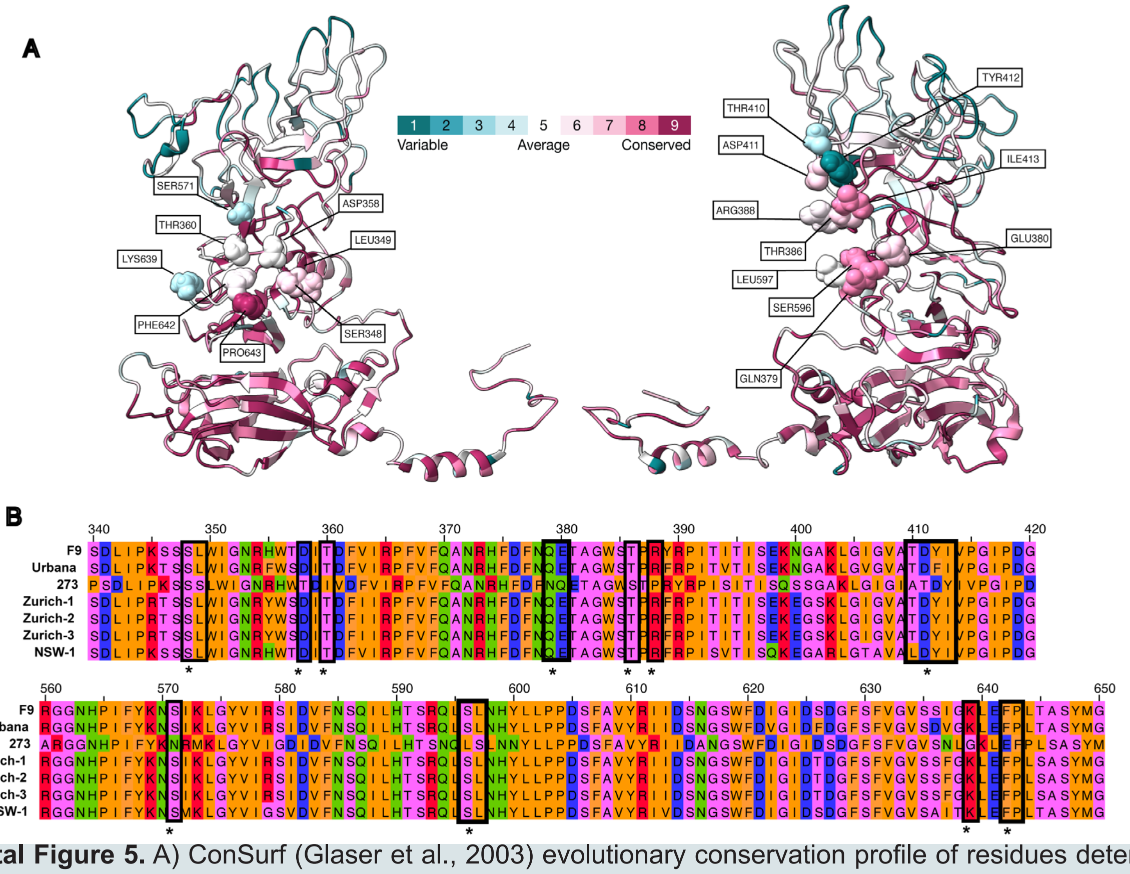


**Supplemental Figure 5.** A) ConSurf (Glaser et al., 2003) evolutionary conservation profile of residues determined by the ChimeraX 'Contacts' tool to interact with VP2 helix c (residues 87-106) in A/C P-dimers (left) and B/D P-dimers (right). Structures are coloured according to the ConSurf evolutionary conservation grades indicated in the legend. Contacting residues are illustrated as spheres with residue name and location indicated. B) Amino acid sequence alignment of residues 340-420 and 560-650 from the FCV isolates used in this study. Residues identified as contacting VP2 helix c are highlighted with squares and asterisks. Residues are coloured according to biochemical properties. Pink: non-polar (Gly, Ala, Ser, Thr); orange: hydrophobic (Cys, Val, Ile, Leu, Pro, Phe, Tyr, Met, Trp); green: polar (Asn, Gln, His); blue: negatively charged (Asp, Glu); red: positively charged (Lys, Arg). Sequences were aligned and visualised in Jalview 2.11.4.0 (Procter *et al.,* 2021).

**Supplemental Methods**

### General Information and Protocols

#### Reagents and Instrumentation

All reagents were purchased from commercial sources and used without further purification unless otherwise stated. Standard Fmoc protected amino acids were purchased from IRIS Biotceh, unless specifically stated differently below. Side chain protecting groups of $N^{\alpha}$-Fmoc amino acids were as follows; Fmoc-Asn(Trt)-OH (Trt = triphenylmethane), Fmoc-Asp(*t*Bu)-OH (*t*Bu = *tert*-butyl), Fmoc-Arg(Pbf)-OH (Pbf = 2,2,4,6,7-pentamethyldihydrobenzofuran-5- sulfonyl), Fmoc-Gln(Trt)-OH, Fmoc-Lys(Boc)-OH (Boc = *tert*-butyloxycarbonyl), Fmoc-Ser(*t*Bu)-OH and Fmoc-Tyr(*t*Bu)-OH.

*N*,*N*-Dimethylformamide (DMF) and diethyl ether were purchased from Rathburn. Trifluoracetic acid (TFA), *N*,*N*'-diisopropylethylamine (DIPEA), *N*,*N*'-diisopropylcarbodiimide (DIC), triisopropylsilane (TIPS), ethyl(hydroxy)cyanoacetate (Oxyma Pure), 1-[bis(dimethylamino)methylene]-1H-1,2,3-triazolo[4,5-b]pyridinium 3-oxid hexafluorophosphate (HATU), (*R*)-2-((((9H-Fluoren-9-yl)methoxy)carbonyl)amino)-2-methyldec-9-enoic acid, (*S*)-2-((((9H-fluoren-9-yl)methoxy)carbonyl)amino)-2-methylhept-6-enoic acid, and benzylidenebis(tricyclohexylphosphine)dichlororuthenium (Grubbs 1st Catalyst) were purchased from Fluorochem. Dichloromethane was purchased from VWR. Acetonitrile (MeCN) and formic acid were purchased from Fisher Scientific. Piperidine, morpholine and acetic anhydride were purchased from Sigma-Aldrich, Merck. Rink-Amide resin was purchased from Activotec.

Analytical reverse-phase high-performance liquid chromatography (RP-HPLC) was performed on a Shimadzu RP-HPLC system with Shimadzu LC-20AT pumps, a Shimadzu SIL20A autosampler and a Shimadzu SPD-20A UV-vis detector using a Phenomenex Aeris™ 5 µm Peptide XB-C18 column (100Å, 150 × 4.6 mm). Peptides were eluted with linear gradients at column-dependent flow rates (1 mL/min for the Aeris), where buffer A = 0.1% TFA in $H_2O$ and buffer B = 0.1% TFA in MeCN. Data is reported as column retention time ($t_R$) in minutes (min). Crude peptides were purified by preparative RP-HPLC using an Agilent Technologies 1260 Infinity II Preparative LC system (monitoring at 214 nm and 280 nm) with a Phenomenex Gemini 5 µm NX-C18 column (110Å, 250 × 21.2 mm). Peptides were eluted with linear gradients (as determined by analytical RP-HPLC) at column-dependent flow rates (10 mL/min for the Gemini), where buffer A = 0.1% TFA in $H_2O$ and buffer B = 0.1% TFA in MeCN. Data is reported as column retention time ($t_R$) in minutes (min).

Liquid chromatography-mass spectrometry (LC-MS) was performed on a Thermo Scientific Dionex Ultimate 3000 LC system coupled to a Thermo Scientific LCQ Fleet Ion trap mass spectrometer using positive mode electrospray ionisation ($ESI^+$). A linear gradient of buffer A (95:5 $H_2O$:MeCN + 0.1% TFA) to buffer B (95:5 MeCN:$H_2O$ + 0.1% TFA) was used over 10 min or 20 min with a Phenomenex Gemini 5 µm NX-C18 column (110Å, 250 × 21.2 mm) with a flow rate of 1 mL/min.

**General Protocol 1 for the Automated Fmoc-SPPS**

Peptides were synthesised using a Syro II peptide synthesiser on a 0.02 mmol scale. The peptides were synthesised using Rink Amide functionalised aminomethyl resin (loading 0.37 mmol/g or loading 0.53 mmol/g), employing a SPPS Fmoc/*t*Bu protecting group strategy. Resin was swelled at 75 °C for 15 min in DMF (0.5 mL). Peptides were elongated in cycles of amino acid coupling followed by Fmoc removal. Coupling reactions were performed using Fmoc-amino acid (5 equiv., 0.5 M in DMF), DIC (5 equiv., 0.5 M in DMF) and Oxyma Pure (5 equiv., 1 M in DMF). Reaction mixtures were stirred at room temperature for 1 h. This procedure was then repeated with fresh reagents at room temperature for a further 1 h. Following coupling, the resin was washed with DMF (6 × 0.6 mL). Fmoc removal was carried out using 40% piperidine in DMF with 5% formic acid (0.5 mL, *v/v/v*) at room temperature for 3 min followed by 20% piperidine in DMF with 5% formic acid (0.5 mL, *v/v/v*) at room temperature for 12 min. Following deprotection, the resin was washed with DMF (6 × 0.5 mL). Resin-bound peptides were washed with dichloromethane (5 × 1 mL) prior to peptide cleavage.

**General Protocol 2 for the Acetyl Capping**

Peptides requiring *N*-terminal acetylation were treated on-resin with acetic anhydride (5 equiv.), DIPEA (10 equiv.) and DMF (6 mL for 0.1 mmol of resin). The reaction mixture was agitated for 2 h at room temperature. Following capping completion, the peptidyl resin was washed with DMF (3 × 5 mL), dichloromethane (3 × 5 mL), and dried under vacuum.

**General Protocol 3 for Resin Cleavage and Global Deprotection**

Peptide cleavage was carried out using a MultiSynTech Peptide Cleavage Station. The resin bound peptides were treated with cleavage cocktail (1 mL, TFA:TIPS:$H_2O$; 95:2.5:2.5 *v/v/v*). The reaction mixture was agitated for 4 h at room temperature. Cleaved peptides were transferred to 15 mL falcons using a further 0.5 mL cleavage cocktail. Peptides were precipitated from a solution of cold diethyl ether (15 mL) and centrifuged (4500 rpm for 5 min). Peptides were dissolved using 0.1% TFA in MeCN:$H_2O$ (2:8, *v/v*) and lyophilised on a Christ Alpha 2–4 LSCBasic freeze drier.

**General Protocol 4 for Ring Closing Metathesis**

Peptides containing (*R*)-2-Fmoc-2-methyldec-9-enoic and (*S*)-2-Fmoc-2-methylhept-6-enoic acid were stapled by on-resin ring-closing metathesis (RCM). Following resin swelling, Grubbs 1st Catalyst (20 mol%) was added in dry dichloroethane (4 mL for 0.1 mmol resin). The reaction mixture was agitated at room temperature for 2 h. The resin was washed with dichloroethane (3 × 1 mL). This procedure was then repeated as described at room temperature for 2 h with replenished reagents. The resin was dried under vacuum.

### Experimental

#### Table of Peptides

**Table 1.** Name and associated alanine substitution, sequence, % yield, % purity, *m/z* and retention time of peptides. Abbreviations: $NH_2$ = *C*-terminal amide, OH = *C*-terminal acid, Ac = *N*-terminal acetyl capped. Detailed characterisation data for peptides can be found in **Supplementary Methods Figures 1-38**.

| Peptide | Sequence | Yield (%) | % Purity | Calculated *m/z* | Observed *m/z* | $t_R$ (mins) |
|---|---|---|---|---|---|---|
| **A1 (N1A)** | *H-AFYDDQLNAIRLSYRNLFKNKKKK-OH* | 5% | 98% | 2974.47 | 2975.12 ± 1.37 | 24.5 |
| **A2 (F2A)** | *H-NAYDDQLNAIRLSYRNLFKNKKKK-OH* | 14% | 97% | 2941.40 | 2941.93 ± 1.36 | 23.5 |
| **A3 (Y3A)** | *H-NFADDQLNAIRLSYRNLFKNKKKK-OH* | 9% | 98% | 2925.40 | 2925.74 ± 1.28 | 25.0 |
| **A4 (D4A)** | *H-NFYADQLNAIRLSYRNLFKNKKKK-OH* | 6% | 95% | 2973.48 | 2974.16 ± 0.94 | 25.6 |
| **A5 (D5A)** | *H-NFYDAQLNAIRLSYRNLFKNKKKK-OH* | 5% | 97% | 2973.48 | 2973.61 ± 0.95 | 25.3 |
| **A6 (Q6A)** | *H-NFYDDALNAIRLSYRNLFKNKKKK-OH* | 9% | 97% | 2960.44 | 2960.50 ± 0.68 | 28.3 |
| **A7 (L7A)** | *H-NFYDDQANAIRLSYRNLFKNKKKK-OH* | 18% | 98% | 2975.41 | 2976.56 ± 1.75 | 23.5 |
| **A8 (N8A)** | *H-NFYDDQLAAIRLSYRNLFKNKKKK-OH* | 16% | 96% | 2974.47 | 2976.53 ± 1.23 | 23.5 |
| **A9 (I10A)** | *H-NFYDDQLNAARLSYRNLFKNKKKK-OH* | 22% | 98% | 2975.41 | 2976.48 ± 1.03 | 23.4 |

| | | | | | | |
|---|---|---|---|---|---|---|
| **A10 (R11A)** | *H-NFYDDQLNAIALSYRNLFKNKKKK-OH* | 8% | 99% | 2932.38 | 2932.42 ± 1.03 | 25.4 |
| **A11 (L12A)** | *H-NFYDDQLNAIRASYRNLFKNKKKK-OH* | 18% | 99% | 2975.41 | 2976.09 ± 0.84 | 24.4 |
| **A12 (S13A)** | *H-NFYDDQLNAIRLAYRNLFKNKKKK-OH* | 6% | 96% | 3001.50 | 3001.62 ± 0.88 | 25.7 |
| **B1 (Y14A)** | *H-NFYDDQLNAIRLSARNLFKNKKKK-OH* | 15% | 98% | 2925.40 | 2925.82 ± 0.67 | 24.7 |
| **B2 (R15A)** | *H-NFYDDQLNAIRLSYANLFKNKKKK- OH* | 16% | 99% | 2932.38 | 932.78 ± 1.12 | 25.1 |
| **B3 (N16A)** | *H-NFYDDQLNAIRLSYRALFKNKKKK-OH* | 8% | 98% | 2974.47 | 2974.79 ± 1.26 | 25.3 |
| **B4 (L17A)** | *H-NFYDDQLNAIRLSYRNAFKNKKKK-OH* | 16% | 96% | 2975.41 | 2975.92 ± 0.29 | 23.6 |
| **B5 (F18A)** | *H-AFYDDQLNAIRLSYRNLAKNKKKK-OH* | 16% | 97% | 2941.40 | 2941.70 ± 1.32 | 23.4 |
| **B6 (K19A)** | *H-NFYDDQLNAIRLSYRNLFANKKKK-OH* | 18% | 98% | 2960.40 | 2960.95 ± 1.13 | 25.0 |
| **B7 (N20A)** | *H-NFYDDQLNAIRLSYRNLFKAKKKK-OH* | 20% | 96% | 2974.47 | 2974.16 ± 0.94 | 25.0 |

### Peptide Synthesis

#### Synthesis of A1

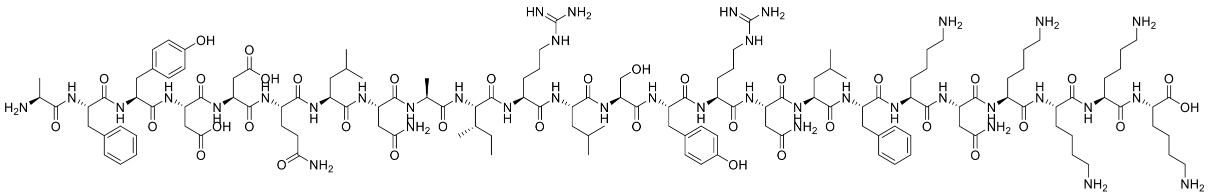

**A1** was synthesised by automated Fmoc-SPPS as described in **General Protocol 1** (0.02 mmol). The final peptide was liberated from the resin and its protecting groups simultaneously removed under the conditions described in **General Protocol 3**. Crude peptide was solubilised in 0.1% TFA in MeCN:$H_2O$ (2:8, *v/v*) and purified by preparative RP-HPLC employing a gradient of 20–60%B over 60 min (*ca.* 1.5%B/min) at a flow rate of 10 mL/min. Fractions were analysed by RP-HPLC and LCMS for compound identification and lyophilised to afford the compound **A1** as a white powder (3.2 mg, 98% purity, 5% overall yield).

**LCMS**: Mass calculated for [$C_{136}H_{217}N_{39}O_{36}$ + H] 2974.47; deconvoluted mass observed: 2975.12 ± 1.37. Charge states; 497.11 $[M+6H]^{6+}$, 596.22 $[M+5H]^{5+}$, 744.76 $[M+4H]^{4+}$, 992.56 $[M+3H]^{3+}$, 1478.56 $[M+2H]^{2+}$.

**RP-HPLC**: $t_R$ = 24.5 min. Phenomenex Aeris™ 5 µm Peptide XB-C18 column (100Å, 150 × 4.6 mm), linear gradient 5%–95%B over 50 min (*ca.* 1.8%B/min) at 1 mL/min.

#### Synthesis of A2

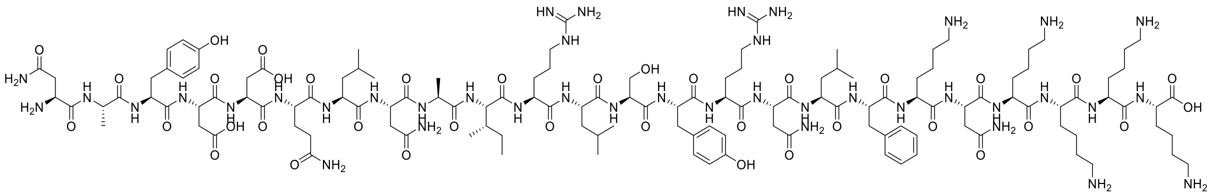

**A2** was synthesised by automated Fmoc-SPPS as described in **General Protocol 1** (0.02 mmol). The final peptide was liberated from the resin and its protecting groups simultaneously removed under the conditions described in **General Protocol 3**. Crude peptide was solubilised in 0.1% TFA in MeCN:$H_2O$ (2:8, *v/v*) and purified by preparative RP-HPLC employing a gradient of 20–60%B over 60 min (*ca.* 1.5%B/min) at a flow rate of 10 mL/min. Fractions were analysed by RP-HPLC and LCMS for compound identification and lyophilised to afford the compound **A2** as a white powder (8.1 mg, 97% purity, 14% overall yield).

**LCMS**: Mass calculated for [$C_{131}H_{214}N_{40}O_{37}$ + H] 2941.40; deconvoluted mass observed: 2941.93 ± 1.36. Charge states; 589.73 $[M+5H]^{5+}$, 736.53 $[M+4H]^{4+}$, 981.52 $[M+3H]^{3+}$, 1471.19 $[M+2H]^{2+}$.

**RP-HPLC**: $t_R$ = 23.5 min. Phenomenex Aeris™ 5 µm Peptide XB-C18 column (100Å, 150 × 4.6 mm), linear gradient 5%–95%B over 50 min (*ca.* 1.8%B/min) at 1 mL/min.

**Synthesis of A3**

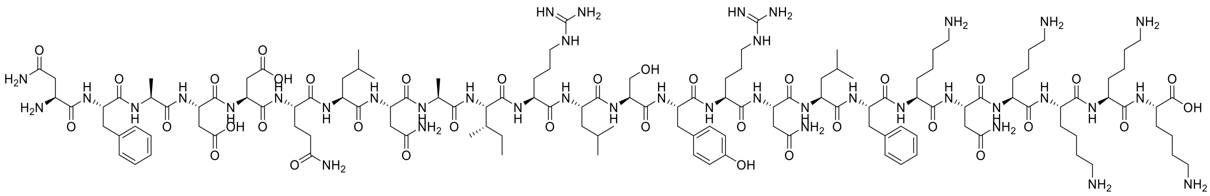

**A3** was synthesised by automated Fmoc-SPPS as described in **General Protocol 1** (0.02 mmol). The final peptide was liberated from the resin and its protecting groups simultaneously removed under the conditions described in **General Protocol 3**. Crude peptide was solubilised in 0.1% TFA in MeCN:$H_2O$ (2:8, *v/v*) and purified by preparative RP-HPLC employing a gradient of 20–60%B over 60 min (*ca.* 1.5%B/min) at a flow rate of 10 mL/min. Fractions were analysed by RP-HPLC and LCMS for compound identification and lyophilised to afford the compound **A3** as a white powder (5.3 mg, 98% purity, 9% overall yield).

**LCMS**: Mass calculated for [$C_{131}H_{214}N_{40}O_{36}$ + H] 2925.40; deconvoluted mass observed: 2925.74 ± 1.28. Charge states; 586.44 $[M+5H]^{5+}$, 732.54 $[M+4H]^{4+}$, 976.15 $[M+3H]^{3+}$, 1463.06 $[M+2H]^{2+}$.

**RP-HPLC**: $t_R$ = 25.0 min. Phenomenex Aeris™ 5 µm Peptide XB-C18 column (100Å, 150 × 4.6 mm), linear gradient 5%–95%B over 50 min (*ca.* 1.8%B/min) at 1 mL/min.

### Synthesis of A4

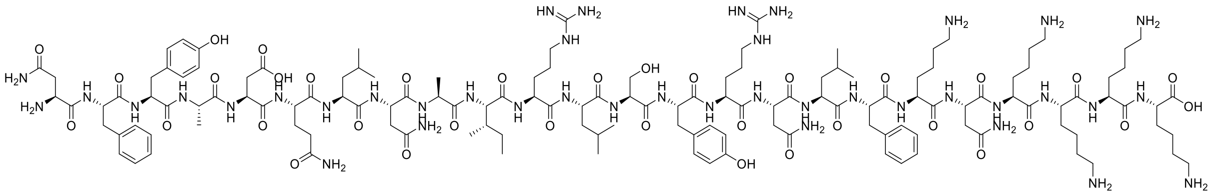

**A4** was synthesised by automated Fmoc-SPPS as described in **General Protocol 1** (0.02 mmol). The final peptide was liberated from the resin and its protecting groups simultaneously removed under the conditions described in **General Protocol 3**. Crude peptide was solubilised in 0.1% TFA in MeCN:$H_2O$ (2:8, *v/v*) and purified by preparative RP-HPLC employing a gradient of 20–60%B over 60 min (*ca.* 1.5%B/min) at a flow rate of 10 mL/min. Fractions were analysed by RP-HPLC and LCMS for compound identification and lyophilised to afford the compound **A4** as a white powder (3.5 mg, 95% purity, 6% overall yield).

**LCMS**: Mass calculated for [$C_{136}H_{218}N_{40}O_{35}$ + H] 2973.48; deconvoluted mass observed: 2974.16 ± 0.94. Charge states; 596.11 $[M+5H]^{5+}$, 744.43 $[M+4H]^{4+}$, 992.16 $[M+3H]^{3+}$, 1487.94 $[M+2H]^{2+}$.

**RP-HPLC**: $t_R$ = 25.6 min. Phenomenex Aeris$^{TM}$ 5 µm Peptide XB-C18 column (100Å, 150 × 4.6 mm), linear gradient 5%–95%B over 50 min (*ca.* 1.8%B/min) at 1 mL/min.

### Synthesis of A5

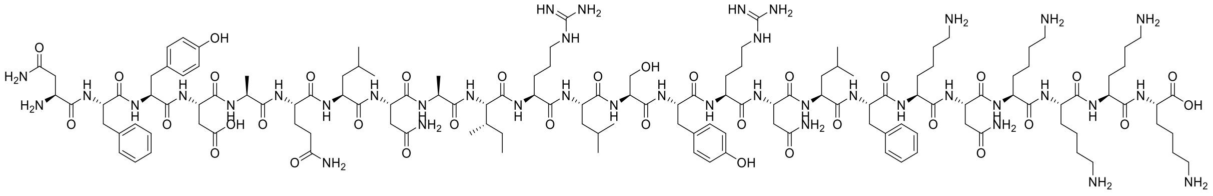

**A5** was synthesised by automated Fmoc-SPPS as described in **General Protocol 1** (0.02 mmol). The final peptide was liberated from the resin and its protecting groups simultaneously removed under the conditions described in **General Protocol 3**. Crude peptide was solubilised in 0.1% TFA in MeCN:$H_2O$ (2:8, *v/v*) and purified by preparative RP-HPLC employing a gradient of 20–60%B over 60 min (*ca.* 1.5%B/min) at a flow rate of 10 mL/min. Fractions were analysed by RP-HPLC and LCMS for compound identification and lyophilised to afford the compound **A5** as a white powder (3.0 mg, 97% purity, 5% overall yield).

**LCMS**: Mass calculated for [$C_{136}H_{218}N_{40}O_{35}$ + H] 2973.48; deconvoluted mass observed: 2973.61 ± 0.95. Charge states; 595.97 $[M+5H]^{5+}$, 744.42 $[M+4H]^{4+}$, 992.12 $[M+3H]^{3+}$, 1487.28 $[M+2H]^{2+}$.

**RP-HPLC**: $t_R$ = 25.3 min. Phenomenex Aeris$^{TM}$ 5 µm Peptide XB-C18 column (100Å, 150 × 4.6 mm), linear gradient 5%–95%B over 50 min (*ca.* 1.8%B/min) at 1 mL/min.

#### Synthesis of A6

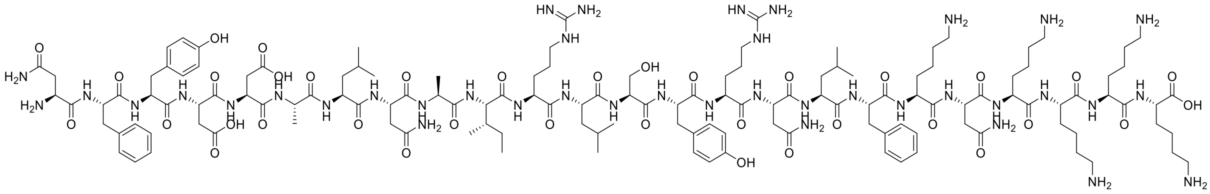

**A6** was synthesised by automated Fmoc-SPPS as described in **General Protocol 1** (0.02 mmol). The final peptide was liberated from the resin and its protecting groups simultaneously removed under the conditions described in **General Protocol 3**. Crude peptide was solubilised in 0.1% TFA in MeCN:$H_2O$ (2:8, *v/v*) and purified by preparative RP-HPLC employing a gradient of 20–60%B over 60 min (*ca.* 1.5%B/min) at a flow rate of 10 mL/min. Fractions were analysed by RP-HPLC and LCMS for compound identification and lyophilised to afford the compound **A6** as a white powder (5.4 mg, 97% purity, 9% overall yield).

**LCMS**: Mass calculated for [$C_{135}H_{215}N_{39}O_{36}$ + H] 2960.44; deconvoluted mass observed: 2960.50 ± 0.68. Charge states; 593.28 $[M+5H]^{5+}$, 741.14 $[M+4H]^{4+}$, 987.74 $[M+3H]^{3+}$, 1480.90 $[M+2H]^{2+}$.

**RP-HPLC**: $t_R$ = 28.3 min. Phenomenex Aeris$^{TM}$ 5 µm Peptide XB-C18 column (100Å, 150 × 4.6 mm), linear gradient 5%–95%B over 50 min (*ca.* 1.8%B/min) at 1 mL/min.

#### Synthesis of A7

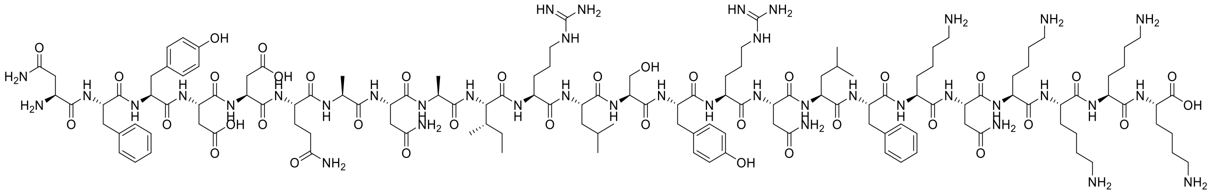

**A7** was synthesised by automated Fmoc-SPPS as described in **General Protocol 1** (0.02 mmol). The final peptide was liberated from the resin and its protecting groups simultaneously removed under the conditions described in **General Protocol 3**. Crude peptide was solubilised in 0.1% TFA in MeCN:$H_2O$ (2:8, *v/v*) and purified by preparative RP-HPLC employing a gradient of 20–60%B over 60 min (*ca.* 1.5%B/min) at a flow rate of 10 mL/min. Fractions were analysed by RP-HPLC and LCMS for compound identification and lyophilised to afford the compound **A7** as a white powder (10.8 mg, 98% purity, 18% overall yield).

**LCMS**: Mass calculated for [$C_{134}H_{212}N_{40}O_{37}$ + H] 2975.41; deconvoluted mass observed: 2976.56 ± 1.75. Charge states; 596.80 $[M+5H]^{5+}$, 745.14 $[M+4H]^{4+}$, 992.88 $[M+3H]^{3+}$, 1488.51 $[M+2H]^{2+}$.

**RP-HPLC**: $t_R$ = 23.5 min. Phenomenex Aeris™ 5 µm Peptide XB-C18 column (100Å, 150 × 4.6 mm), linear gradient 5%–95%B over 50 min (*ca.* 1.8%B/min) at 1 mL/min.

**Synthesis of A8**

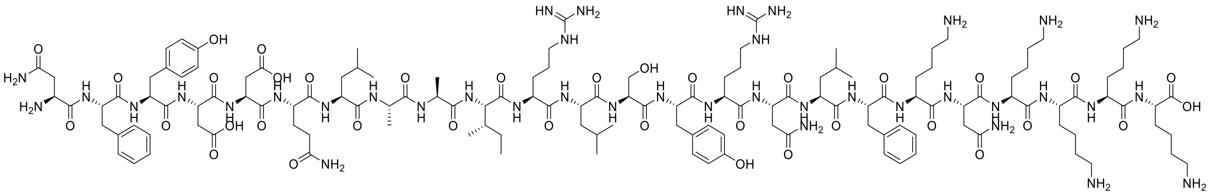

**A8** was synthesised by automated Fmoc-SPPS as described in **General Protocol 1** (0.02 mmol). The final peptide was liberated from the resin and its protecting groups simultaneously removed under the conditions described in **General Protocol 3**. Crude peptide was solubilised in 0.1% TFA in MeCN:$H_2O$ (2:8, *v/v*) and purified by preparative RP-HPLC employing a gradient of 20–60%B over 60 min (*ca.* 1.5%B/min) at a flow rate of 10 mL/min. Fractions were analysed by RP-HPLC and LCMS for compound identification and lyophilised to afford the compound **A8** as a white powder (9.3 mg, 96% purity, 16% overall yield).

**LCMS**: Mass calculated for [$C_{136}H_{217}N_{39}O_{36}$ + H] 2974.47; deconvoluted mass observed: 2976.53 ± 1.23. Charge states; 596.67 $[M+5H]^{5+}$, 744.9 $[M+4H]^{4+}$, 992.90 $[M+3H]^{3+}$, 1489.05 $[M+2H]^{2+}$.

**RP-HPLC**: $t_R$ = 23.5 min. Phenomenex Aeris$^{TM}$ 5 µm Peptide XB-C18 column (100Å, 150 × 4.6 mm), linear gradient 5%–95%B over 50 min (*ca.* 1.8%B/min) at 1 mL/min.

**Synthesis of A9**

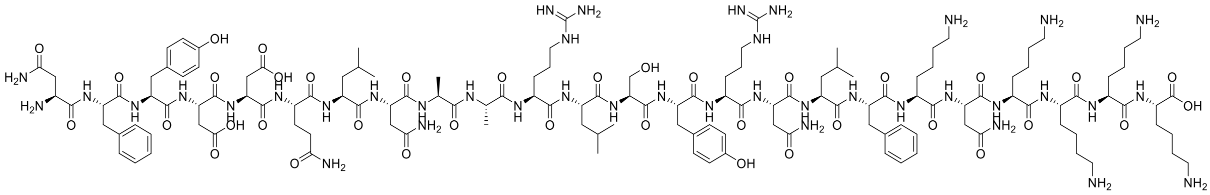

**A9** was synthesised by automated Fmoc-SPPS as described in **General Protocol 1** (0.02 mmol). The final peptide was liberated from the resin and its protecting groups simultaneously removed under the conditions described in **General Protocol 3**. Crude peptide was solubilised in 0.1% TFA in MeCN:$H_2O$ (2:8, *v/v*) and purified by preparative RP-HPLC employing a gradient of 20–60%B over 60 min (*ca.* 1.5%B/min) at a flow rate of 10 mL/min. Fractions were analysed by RP-HPLC and LCMS for compound identification and lyophilised to afford the compound **A9** as a white powder (12.6 mg, 98% purity, 22% overall yield).

**LCMS**: Mass calculated for [$C_{134}H_{212}N_{40}O_{37}$ + H] 2975.41; deconvoluted mass observed: 2976.48 ± 1.03. Charge states; 596.53 $[M+5H]^{5+}$, 745.02 $[M+4H]^{4+}$, 992.90 $[M+3H]^{3+}$.

**RP-HPLC**: $t_R$ = 23.4 min. Phenomenex Aeris$^{TM}$ 5 µm Peptide XB-C18 column (100Å, 150 × 4.6 mm), linear gradient 5%–95%B over 50 min (*ca.* 1.8%B/min) at 1 mL/min.

**Synthesis of A10**

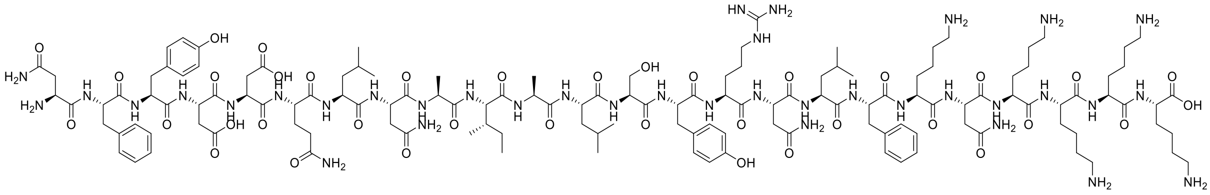

**A10** was synthesised by automated Fmoc-SPPS as described in **General Protocol 1** (0.02 mmol). The final peptide was liberated from the resin and its protecting groups simultaneously removed under the conditions described in **General Protocol 3**. Crude peptide was solubilised in 0.1% TFA in MeCN:$H_2O$ (2:8, *v/v*) and purified by preparative RP-HPLC employing a gradient of 20–60%B over 60 min (*ca.* 1.5%B/min)

at a flow rate of 10 mL/min. Fractions were analysed by RP-HPLC and LCMS for compound identification and lyophilised to afford the compound **A10** as a white powder (4.5 mg, 99% purity, 8% overall yield).

**LCMS**: Mass calculated for [$C_{134}H_{211}N_{37}O_{37}$ + H] 2932.38; deconvoluted mass observed: 2932.42 ± 1.03. Charge states; 587.73 $[M+5H]^{5+}$, 734.17 $[M+4H]^{4+}$, 978.39 $[M+3H]^{3+}$, 1466.59 $[M+2H]^{2+}$.

**RP-HPLC**: $t_R$ = 25.4 min. Phenomenex Aeris™ 5 µm Peptide XB-C18 column (100Å, 150 × 4.6 mm), linear gradient 5%–95%B over 50 min (*ca.* 1.8%B/min) at 1 mL/min.

**Synthesis of A11**

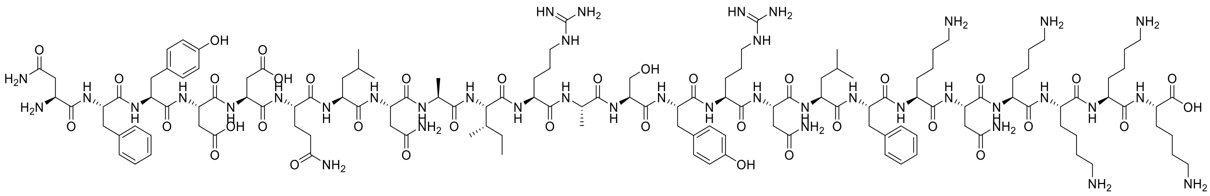

**A11** was synthesised by automated Fmoc-SPPS as described in **General Protocol 1** (0.02 mmol). The final peptide was liberated from the resin and its protecting groups simultaneously removed under the conditions described in **General Protocol 3.** Crude peptide was solubilised in 0.1% TFA in MeCN:$H_2O$ (2:8, *v/v*) and purified by preparative RP-HPLC employing a gradient of 20–60%B over 60 min (*ca.* 1.5%B/min) at a flow rate of 10 mL/min. Fractions were analysed by RP-HPLC and LCMS for compound identification and lyophilised to afford the compound **A11** as a white powder (10.6 mg, 99% purity, 18% overall yield).

**LCMS**: Mass calculated for [$C_{134}H_{212}N_{40}O_{37}$ + H] 2975.41; deconvoluted mass observed: 2976.09 ± 0.84. Charge states; 596.41 $[M+5H]^{5+}$, 744.93 $[M+4H]^{4+}$, 992.83 $[M+3H]^{3+}$.

**RP-HPLC**: $t_R$ = 24.4 min. Phenomenex Aeris™ 5 µm Peptide XB-C18 column (100Å, 150 × 4.6 mm), linear gradient 5%–95%B over 50 min (*ca.* 1.8%B/min) at 1 mL/min.

#### Synthesis of A12

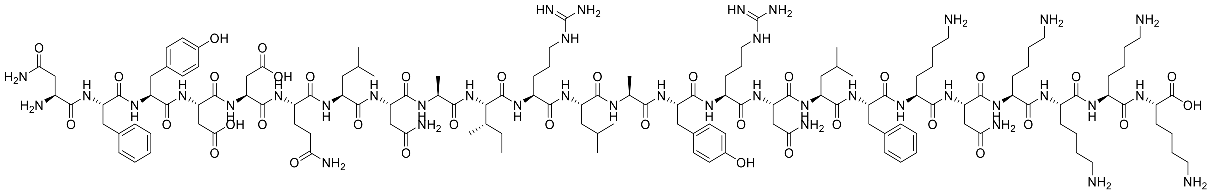

**A12** was synthesised by automated Fmoc-SPPS as described in **General Protocol 1** (0.02 mmol). The final peptide was liberated from the resin and its protecting groups simultaneously removed under the conditions described in **General Protocol 3**. Crude peptide was solubilised in 0.1% TFA in MeCN:$H_2O$ (2:8, *v/v*) and purified by preparative RP-HPLC employing a gradient of 20–60%B over 60 min (*ca.* 1.5%B/min) at a flow rate of 10 mL/min. Fractions were analysed by RP-HPLC and LCMS for compound identification and lyophilised to afford the compound **A12** as a white powder (3.5 mg, 96% purity, 6% overall yield).

**LCMS**: Mass calculated for [$C_{137}H_{218}N_{40}O_{36}$ + H] 3001.50; deconvoluted mass observed: 3001.62 ± 0.88. Charge states; 601.56 $[M+5H]^{5+}$, 751.43 $[M+4H]^{4+}$, 1001.41 $[M+3H]^{3+}$, 1501.27 $[M+2H]^{2+}$.

**RP-HPLC**: $t_R$ = 25.7 min. Phenomenex Aeris™ 5 µm Peptide XB-C18 column (100Å, 150 × 4.6 mm), linear gradient 5%–95%B over 50 min (*ca.* 1.8%B/min) at 1 mL/min.

#### Synthesis of B1

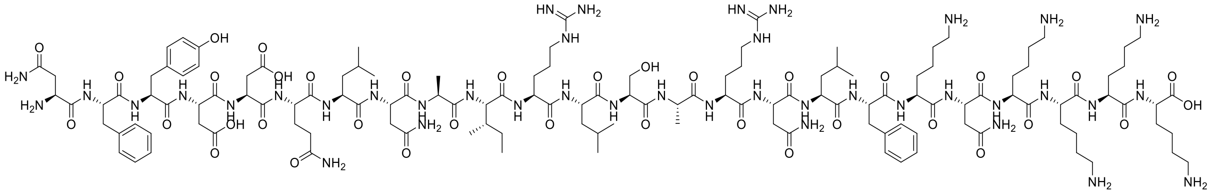

**B1** was synthesised by automated Fmoc-SPPS as described in **General Protocol 1** (0.02 mmol). The final peptide was liberated from the resin and its protecting groups simultaneously removed under the conditions described in **General Protocol 3**. Crude peptide was solubilised in 0.1% TFA in MeCN:$H_2O$ (2:8, *v/v*) and purified by preparative RP-HPLC employing a gradient of 20–60%B over 60 min (*ca.* 1.5%B/min) at a flow rate of 10 mL/min. Fractions were analysed by RP-HPLC and LCMS for compound identification and lyophilised to afford the compound **B1** as a white powder (8.8 mg, 98% purity, 15% overall yield).

**LCMS**: Mass calculated for [$C_{131}H_{214}N_{40}O_{36}$ + H] 2925.40; deconvoluted mass observed: 2925.82 ± 0.67. Charge states; 586.33 $[M+5H]^{5+}$, 732.48 $[M+4H]^{4+}$, 976.23 $[M+3H]^{3+}$, 1463.51 $[M+2H]^{2+}$.

**RP-HPLC**: $t_R$ = 24.7 min. Phenomenex Aeris™ 5 µm Peptide XB-C18 column (100Å, 150 × 4.6 mm), linear gradient 5%–95%B over 50 min (*ca.* 1.8%B/min) at 1 mL/min.

**Synthesis of B2**

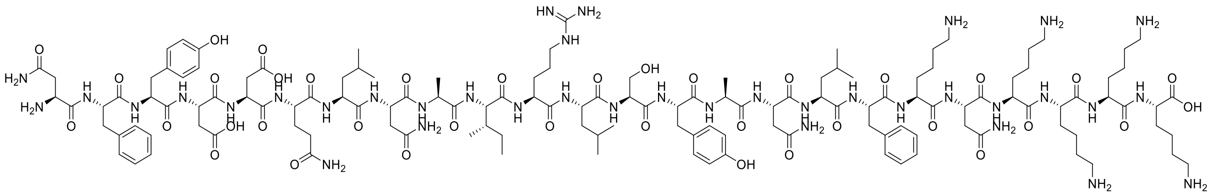

**B2** was synthesised by automated Fmoc-SPPS as described in **General Protocol 1** (0.02 mmol). The final peptide was liberated from the resin and its protecting groups simultaneously removed under the conditions described in **General Protocol 3**. Crude peptide was solubilised in 0.1% TFA in MeCN:$H_2O$ (2:8, *v/v*) and purified by preparative RP-HPLC employing a gradient of 20–60%B over 60 min (*ca.* 1.5%B/min) at a flow rate of 10 mL/min. Fractions were analysed by RP-HPLC and LCMS for compound identification and lyophilised to afford the compound **B2** as a white powder (9.1 mg, 99% purity, 16% overall yield).

**LCMS**: Mass calculated for [$C_{134}H_{211}N_{37}O_{37}$ + H] 2932.38; deconvoluted mass observed: 2932.78 ± 1.12. Charge states; 587.83 $[M+5H]^{5+}$, 734.25 $[M+4H]^{4+}$, 978.51 $[M+3H]^{3+}$, 1466.72 $[M+2H]^{2+}$.

**RP-HPLC**: $t_R$ = 25.1 min. Phenomenex Aeris™ 5 µm Peptide XB-C18 column (100Å, 150 × 4.6 mm), linear gradient 5%–95%B over 50 min (*ca.* 1.8%B/min) at 1 mL/min.

**Synthesis of B3**

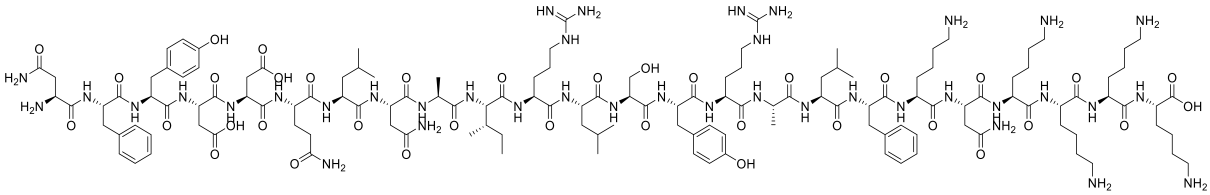

**B3** was synthesised by automated Fmoc-SPPS as described in **General Protocol 1** (0.02 mmol). The final peptide was liberated from the resin and its protecting groups simultaneously removed under the conditions described in **General Protocol 3**. Crude peptide was solubilised in 0.1% TFA in MeCN:$H_2O$ (2:8, *v/v*) and purified by preparative RP-HPLC employing a gradient of 20–60%B over 60 min (*ca.* 1.5%B/min) at a flow rate of 10 mL/min. Fractions were analysed by RP-HPLC and LCMS for compound identification and lyophilised to afford the compound **B3** as a white powder (5.0 mg, 98% purity, 8% overall yield).

**LCMS**: Mass calculated for [$C_{136}H_{217}N_{39}O_{36}$ + H] 2974.47; deconvoluted mass observed: 2974.79 ± 1.26. Charge states; 596.25 $[M+5H]^{5+}$, 744.75 $[M+4H]^{4+}$, 992.57 $[M+3H]^{3+}$, 1487.59 $[M+2H]^{2+}$.

**RP-HPLC**: $t_R$ = 25.3 min. Phenomenex Aeris™ 5 µm Peptide XB-C18 column (100Å, 150 × 4.6 mm), linear gradient 5%–95%B over 50 min (*ca.* 1.8%B/min) at 1 mL/min.

**Synthesis of B4**

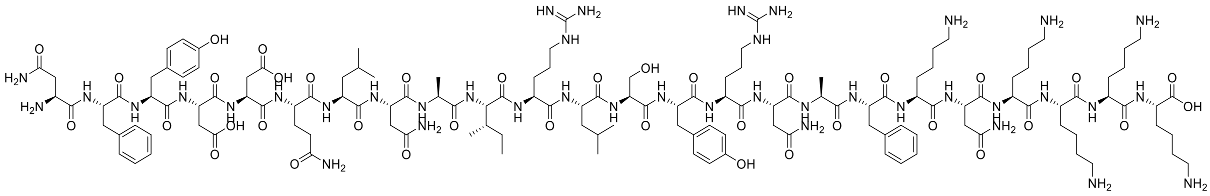

**B4** was synthesised by automated Fmoc-SPPS as described in **General Protocol 1** (0.02 mmol). The final peptide was liberated from the resin and its protecting groups simultaneously removed under the conditions described in **General Protocol 3**. Crude peptide was solubilised in 0.1% TFA in MeCN:$H_2O$ (2:8, *v/v*) and purified by preparative RP-HPLC employing a gradient of 20–60%B over 60 min (*ca.* 1.5%B/min) at a flow rate of 10 mL/min. Fractions were analysed by RP-HPLC and LCMS for compound identification and lyophilised to afford the compound **B4** as a white powder (9.5 mg, 96% purity, 16% overall yield).

**LCMS**: Mass calculated for [$C_{134}H_{212}N_{40}O_{37}$ + H] 2975.41; deconvoluted mass observed: 2975.92 ± 0.29. Charge states; 496.02 $[M+6H]^{6+}$, 596.24 $[M+5H]^{5+}$, 744.90 $[M+4H]^{4+}$, 992.92 $[M+3H]^{3+}$.

**RP-HPLC**: $t_R$ = 23.6 min. Phenomenex Aeris™ 5 µm Peptide XB-C18 column (100Å, 150 × 4.6 mm), linear gradient 5%–95%B over 50 min (*ca.* 1.8%B/min) at 1 mL/min.

#### Synthesis of B5

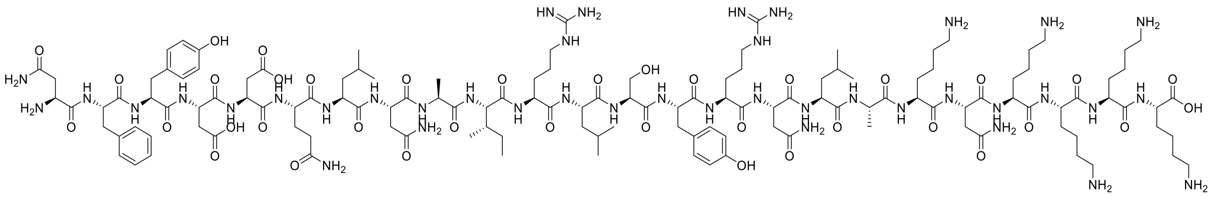

**B5** was synthesised by automated Fmoc-SPPS as described in **General Protocol 1** (0.02 mmol). The final peptide was liberated from the resin and its protecting groups simultaneously removed under the conditions described in **General Protocol 3**. Crude peptide was solubilised in 0.1% TFA in MeCN:$H_2O$ (2:8, *v/v*) and purified by preparative RP-HPLC employing a gradient of 20–60%B over 60 min (*ca.* 1.5%B/min) at a flow rate of 10 mL/min. Fractions were analysed by RP-HPLC and LCMS for compound identification and lyophilised to afford the compound **B5** as a white powder (9.6 mg, 97% purity, 16% overall yield).

**LCMS**: Mass calculated for [$C_{131}H_{214}N_{40}O_{37}$ + H] 2941.40; deconvoluted mass observed: 2941.70 ± 1.32. Charge states; 589.69 $[M+5H]^{5+}$, 736.43 $[M+4H]^{4+}$, 981.45 $[M+3H]^{3+}$, 1471.13 $[M+2H]^{2+}$.

**RP-HPLC**: $t_R$ = 23.4 min. Phenomenex Aeris™ 5 µm Peptide XB-C18 column (100Å, 150 × 4.6 mm), linear gradient 5%–95%B over 50 min (*ca.* 1.8%B/min) at 1 mL/min.

#### Synthesis of B6

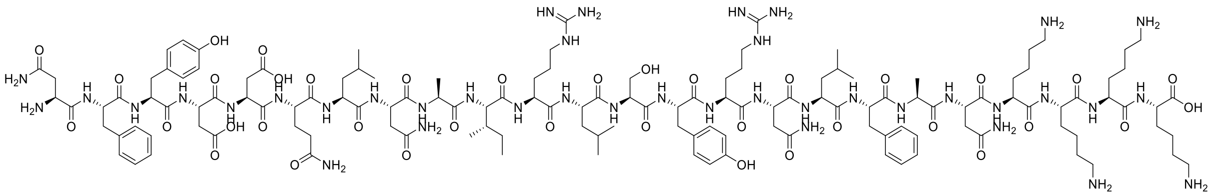

**B6** was synthesised by automated Fmoc-SPPS as described in **General Protocol 1** (0.02 mmol). The final peptide was liberated from the resin and its protecting groups simultaneously removed under the conditions described in **General Protocol 3**. Crude peptide was solubilised in 0.1% TFA in MeCN:$H_2O$ (2:8, *v/v*) and purified by preparative RP-HPLC employing a gradient of 20–60%B over 60 min (*ca.* 1.5%B/min)

at a flow rate of 10 mL/min. Fractions were analysed by RP-HPLC and LCMS for compound identification and lyophilised to afford the compound **B6** as a white powder (10.5 mg, 98% purity, 18% overall yield).

**LCMS**: Mass calculated for [$C_{134}H_{211}N_{39}O_{37}$ + H] 2960.40; deconvoluted mass observed: 2960.95 ± 1.13. Charge states; 593.46 $[M+5H]^{5+}$, 741.28 $[M+4H]^{4+}$, 987.87 $[M+3H]^{3+}$, 1480.64 $[M+2H]^{2+}$.

**RP-HPLC**: $t_R$ = 25.0 min. Phenomenex Aeris™ 5 µm Peptide XB-C18 column (100Å, 150 × 4.6 mm), linear gradient 5%–95%B over 50 min (*ca.* 1.8%B/min) at 1 mL/min.

**Synthesis of B7**

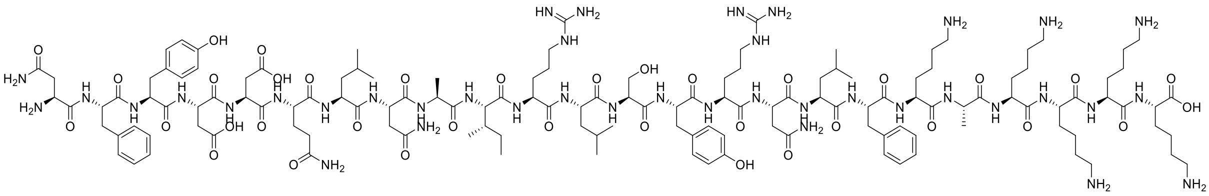

**B7** was synthesised by automated Fmoc-SPPS as described in **General Protocol 1** (0.02 mmol). The final peptide was liberated from the resin and its protecting groups simultaneously removed under the conditions described in **General Protocol 3**. Crude peptide was solubilised in 0.1% TFA in MeCN:$H_2O$ (2:8, *v/v*) and purified by preparative RP-HPLC employing a gradient of 20–60%B over 60 min (*ca.* 1.5%B/min) at a flow rate of 10 mL/min. Fractions were analysed by RP-HPLC and LCMS for compound identification and lyophilised to afford the compound **B7** as a white powder (11.8 mg, 96% purity, 20% overall yield).

**LCMS**: Mass calculated for [$C_{136}H_{217}N_{39}O_{36}$ + H] 2974.47; deconvoluted mass observed: 2974.16 ± 0.94. Charge states; 598.11 $[M+5H]^{5+}$, 744.43 $[M+4H]^{4+}$, 992.16 $[M+3H]^{3+}$, 1487.94 $[M+2H]^{2+}$.

**RP-HPLC**: $t_R$ = 25.0 min. Phenomenex Aeris™ 5 µm Peptide XB-C18 column (100Å, 150 × 4.6 mm), linear gradient 5%–95%B over 50 min (*ca.* 1.8%B/min) at 1 mL/min.

### Characterisation

#### Reverse Phase – High Performance Liquid Chromatography (RP-HPLC)

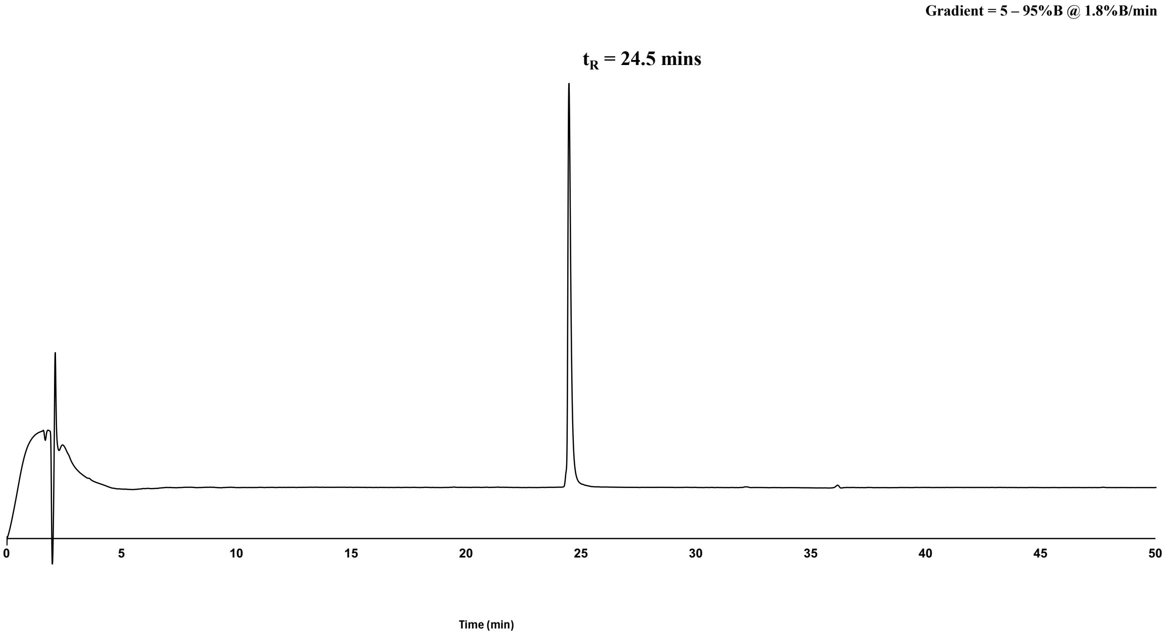


**Supplementary Methods Figure 1.** Analytical RP-HPLC chromatogram (214 nm) of purified peptide, **A1** (*ca.* 98% as analysed by peak area). Phenomenex Aeris™ 5 μm Peptide XB-C18 column (100Å, 150 × 4.6 mm), linear gradient 5%–95%B over 50 min (*ca.* 1.8%B/min) at 1 mL/min. $t_R$ = 24.5 mins.

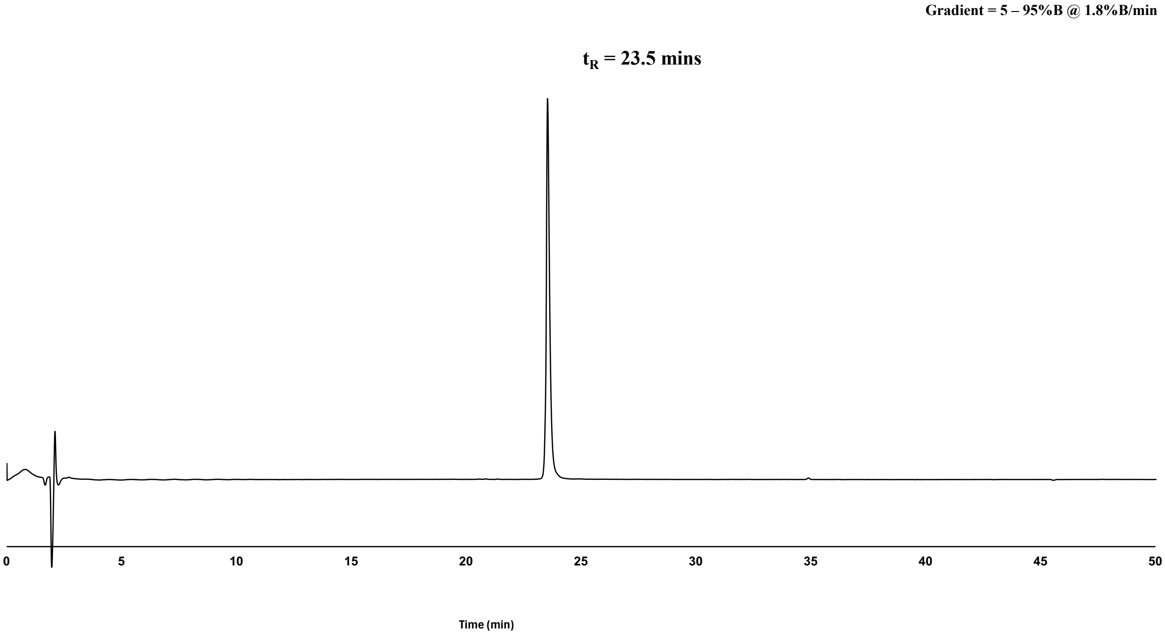

**Supplementary Methods Figure 2.** Analytical RP-HPLC chromatogram (214 nm) of purified peptide, **A2** (*ca.* 97% as analysed by peak area). Phenomenex Aeris™ 5 µm Peptide XB-C18 column (100Å, 150 × 4.6 mm), linear gradient 5%–95%B over 50 min (*ca.* 1.8%B/min) at 1 mL/min. $t_R$ = 23.5 mins.

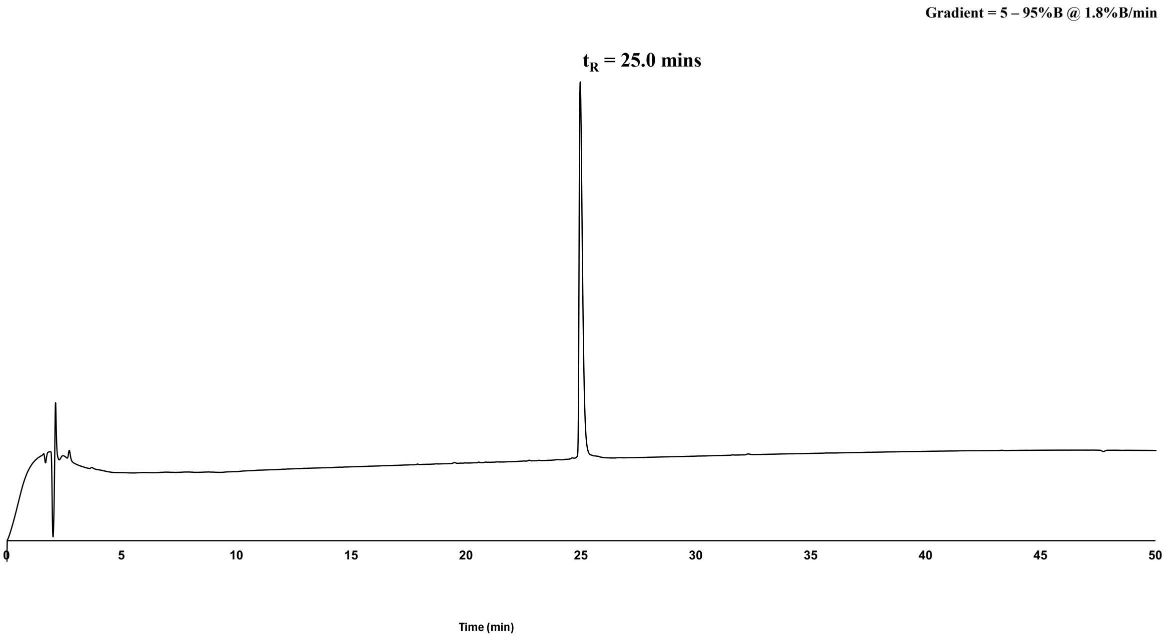


**Supplementary Methods Figure 3.** Analytical RP-HPLC chromatogram (214 nm) of purified peptide, **A3** (*ca.* 98% as analysed by peak area). Phenomenex Aeris™ 5 µm Peptide XB-C18 column (100Å, 150 × 4.6 mm), linear gradient 5%–95%B over 50 min (*ca.* 1.8%B/min) at 1 mL/min. $t_R$ = 25.0 mins.

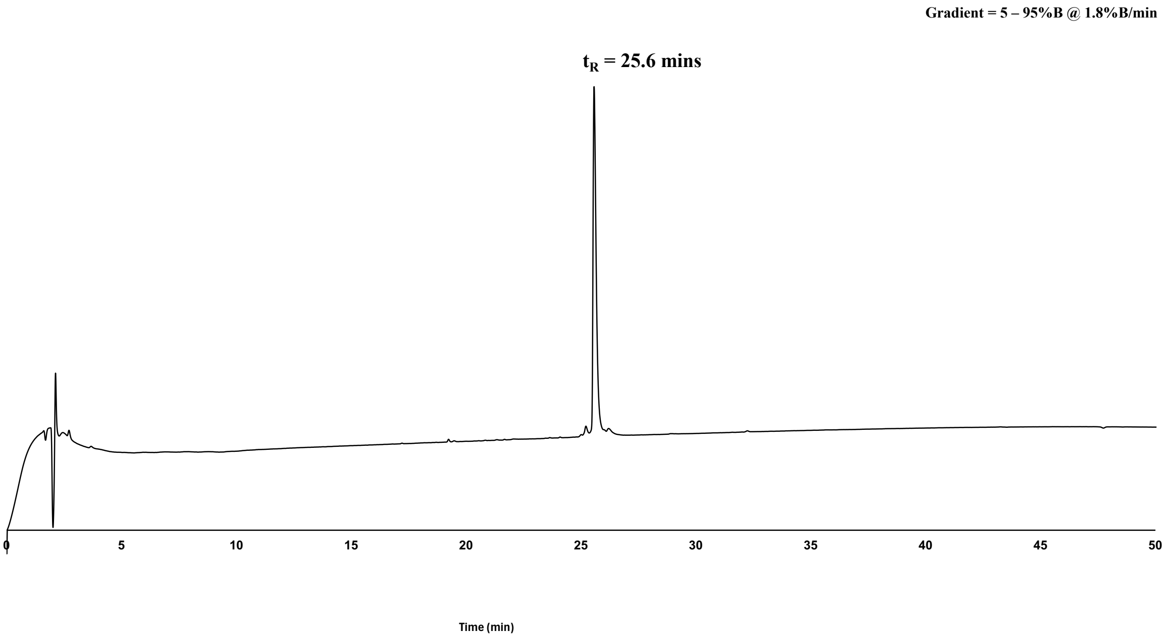

**Supplementary Methods Figure 4**.Analytical RP-HPLC chromatogram (214 nm) of purified peptide, **A4** (*ca.* 95% as analysed by peak area). Phenomenex Aeris[TM] 5 µm Peptide XB-C18 column (100Å, 150 × 4.6 mm), linear gradient 5%–95%B over 50 min (*ca.* 1.8%B/min) at 1 mL/min. $t_R$ = 25.6 mins.

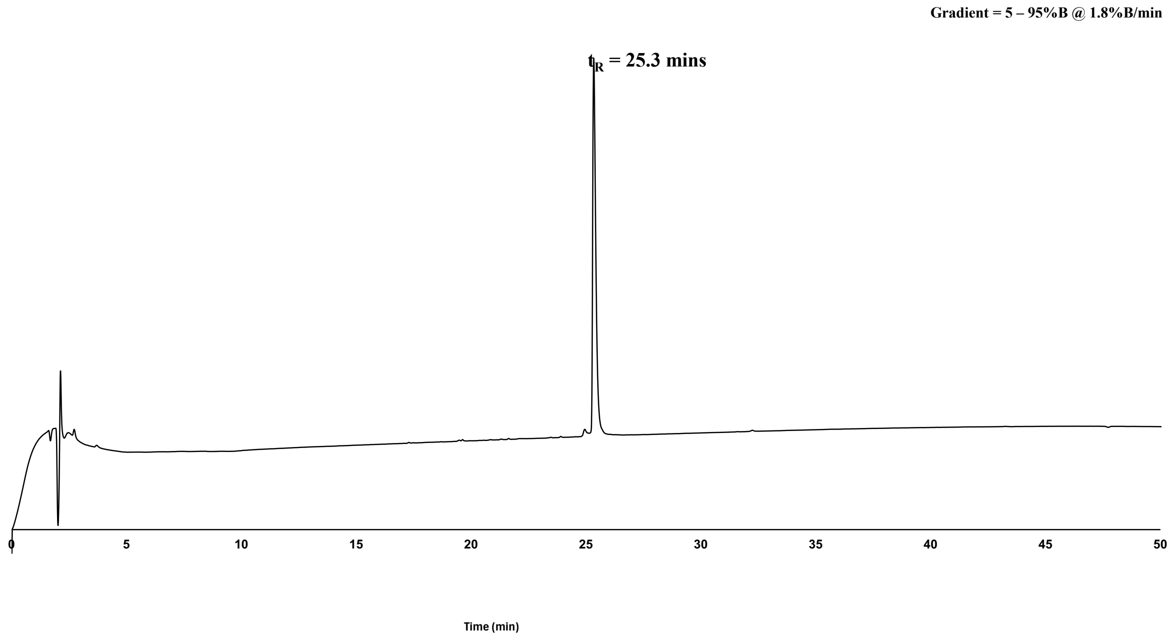


**Supplementary Methods Figure 5.** Analytical RP-HPLC chromatogram (214 nm) of purified peptide, **A5** (*ca.* 97% as analysed by peak area). Phenomenex Aeris[TM] 5 µm Peptide XB-C18 column (100Å, 150 × 4.6 mm), linear gradient 5%–95%B over 50 min (*ca.* 1.8%B/min) at 1 mL/min. $t_R$ = 25.3 mins.

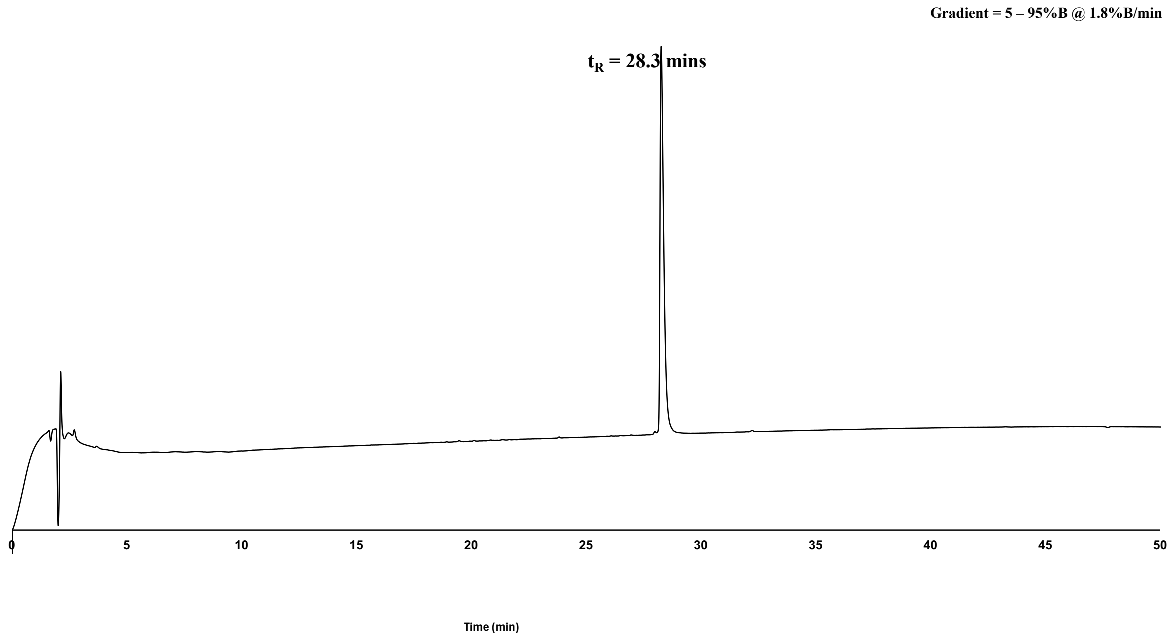

**Supplementary Methods Figure 6.** Analytical RP-HPLC chromatogram (214 nm) of purified peptide, **A6** (*ca.* 97% as analysed by peak area). Phenomenex Aeris[TM] 5 µm Peptide XB-C18 column (100Å, 150 × 4.6 mm), linear gradient 5%–95%B over 50 min (*ca.* 1.8%B/min) at 1 mL/min. $t_R$ = 28.3 mins.

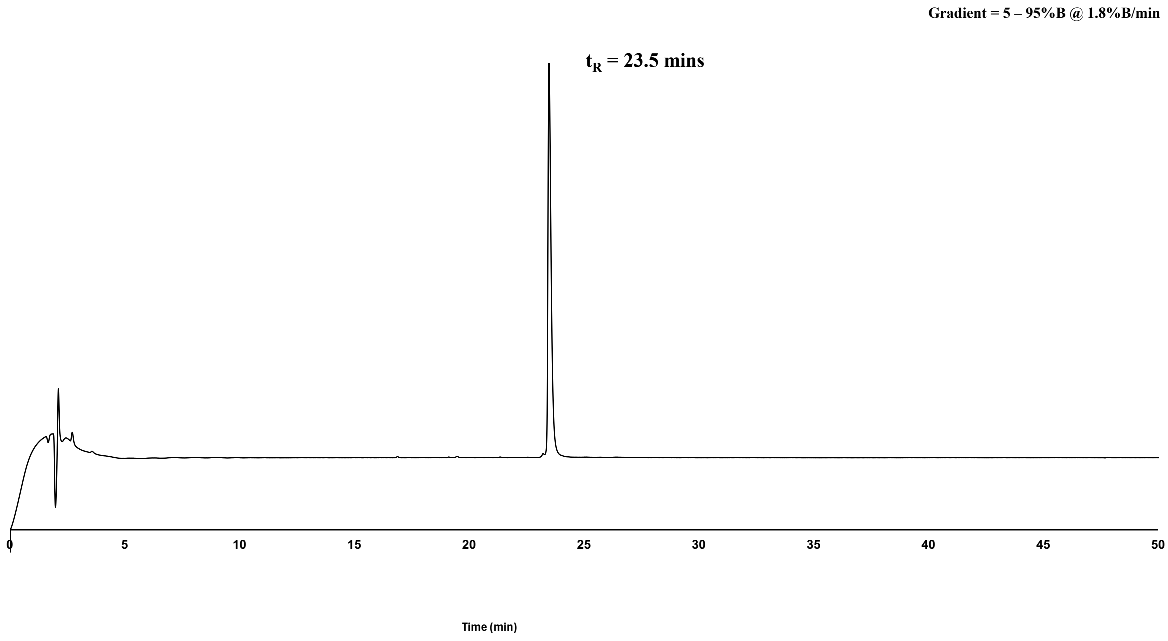


**Supplementary Methods Figure 7.** Analytical RP-HPLC chromatogram (214 nm) of purified peptide, **A7** (*ca.* 98% as analysed by peak area). Phenomenex Aeris[TM] 5 µm Peptide XB-C18 column (100Å, 150 × 4.6 mm), linear gradient 5%–95%B over 50 min (*ca.* 1.8%B/min) at 1 mL/min. $t_R$ = 23.5 mins.

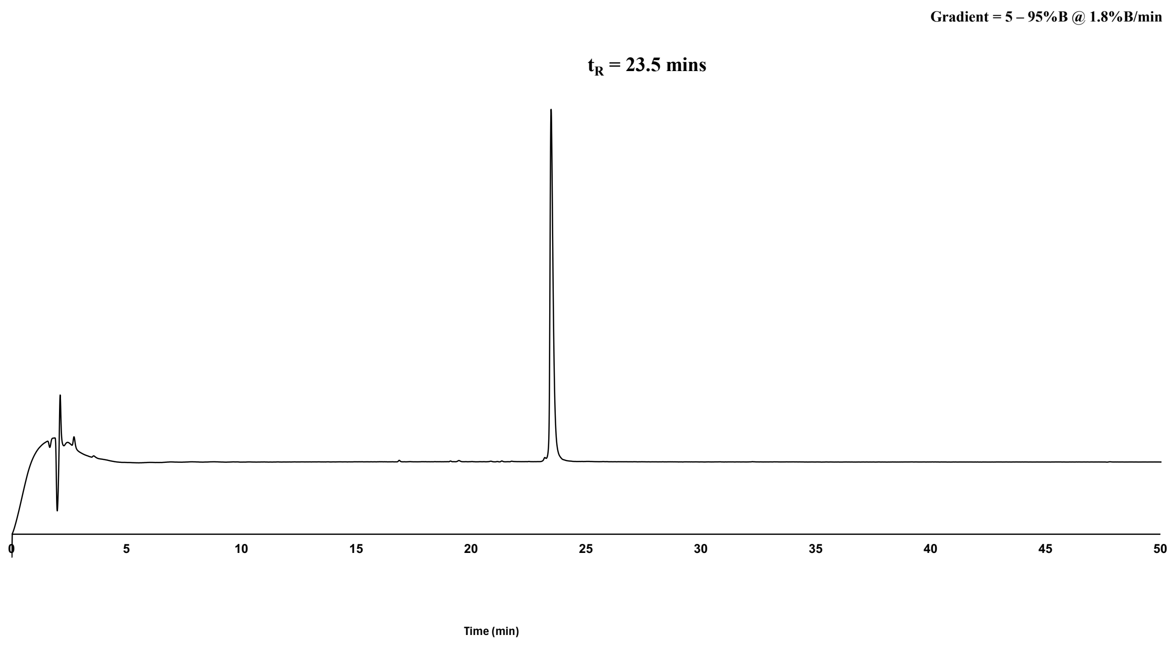


**Supplementary Methods Figure 8.** Analytical RP-HPLC chromatogram (214 nm) of purified peptide, **A8** (*ca.* 96% as analysed by peak area). Phenomenex Aeris™ 5 μm Peptide XB-C18 column (100Å, 150 × 4.6 mm), linear gradient 5%–95%B over 50 min (*ca.* 1.8%B/min) at 1 mL/min. $t_R$ = 23.5 mins.

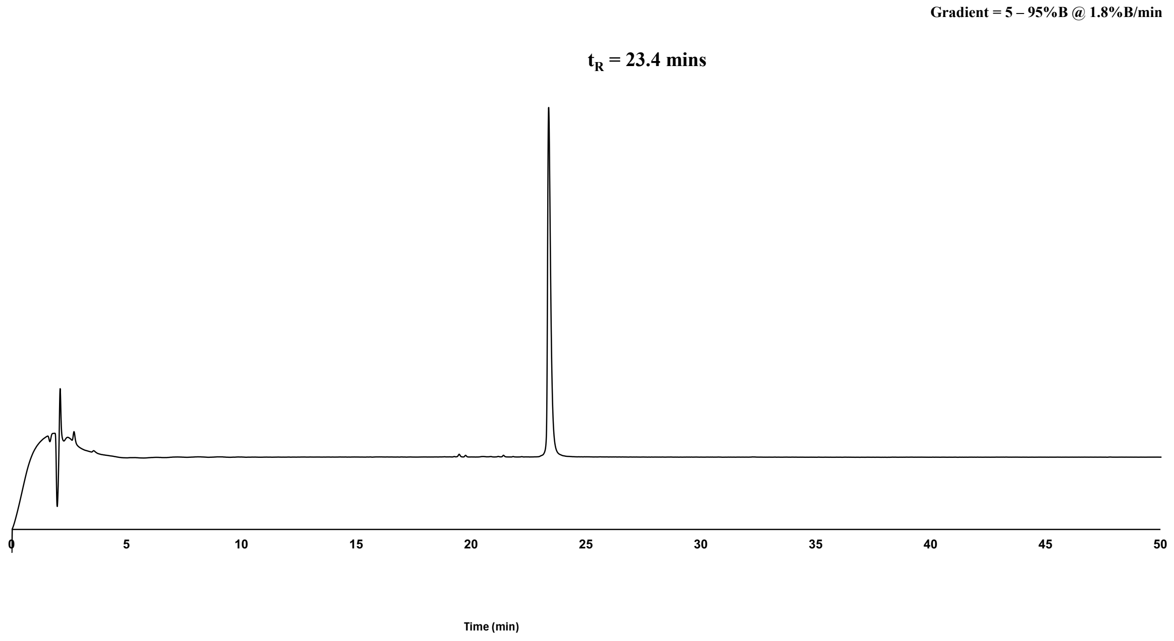


**Supplementary Methods Figure 9.** Analytical RP-HPLC chromatogram (214 nm) of purified peptide, **A9** (*ca.* 98% as analysed by peak area). Phenomenex Aeris™ 5 μm Peptide XB-C18 column (100Å, 150 × 4.6 mm), linear gradient 5%–95%B over 50 min (*ca.* 1.8%B/min) at 1 mL/min. $t_R$ = 23.4 mins.

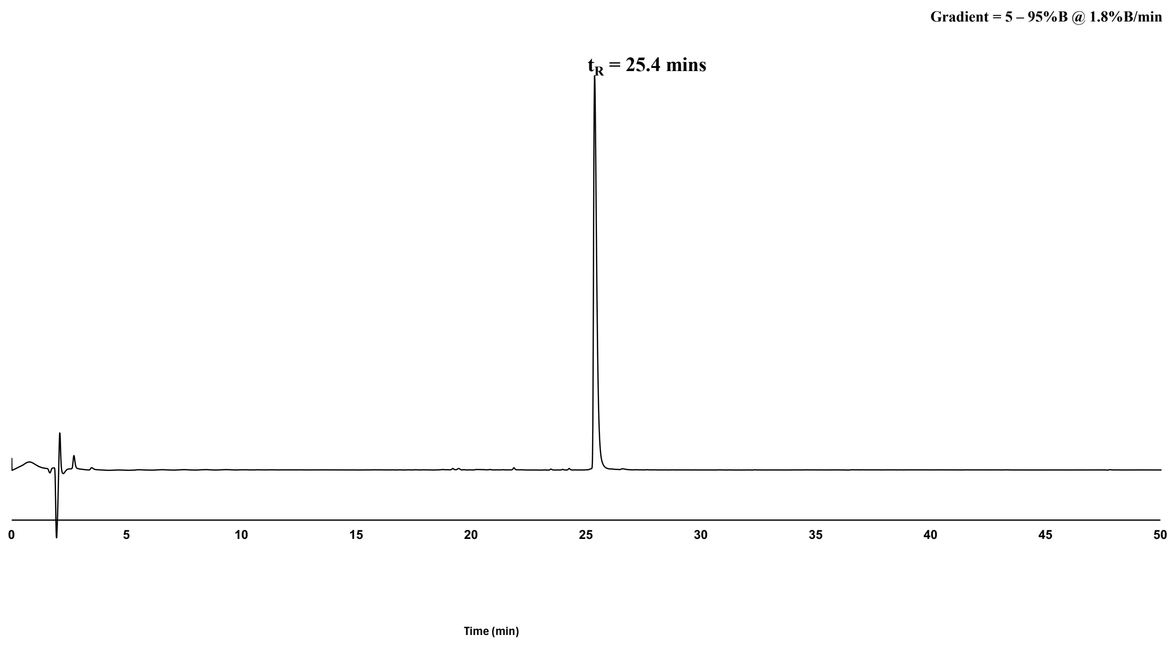


**Supplementary Methods Figure 10.** Analytical RP-HPLC chromatogram (214 nm) of purified peptide, **A10** (*ca.* 99% as analysed by peak area). Phenomenex Aeris$^{TM}$ 5 µm Peptide XB-C18 column (100Å, 150 × 4.6 mm), linear gradient 5%–95%B over 50 min (*ca.* 1.8%B/min) at 1 mL/min. $t_R$ = 25.4 mins.

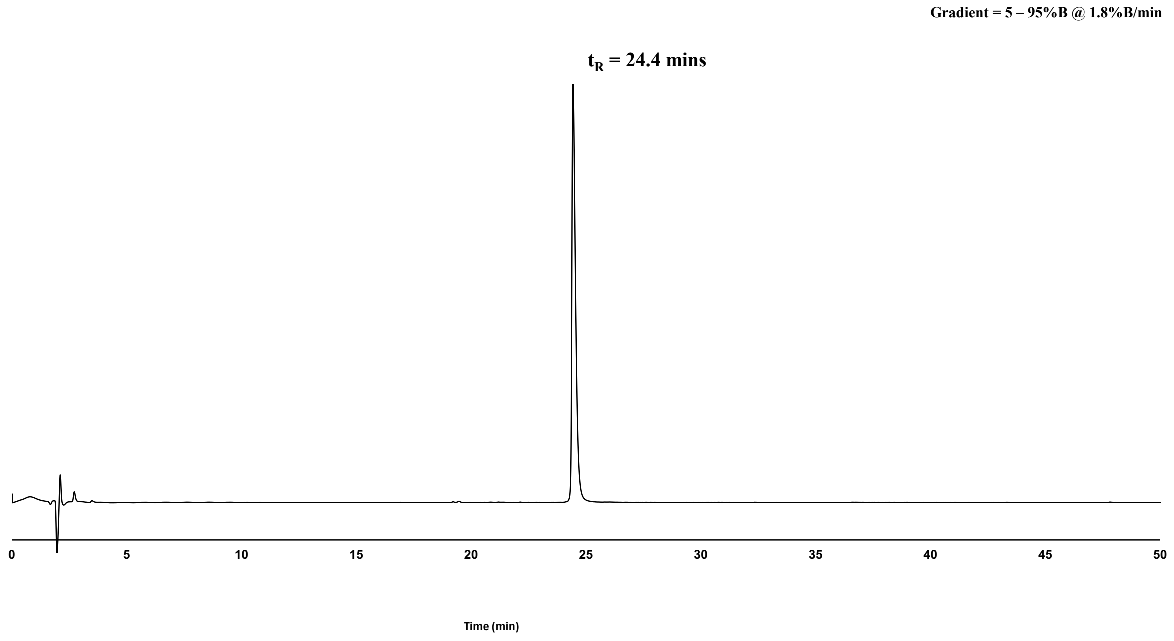


**Supplementary Methods Figure 11.** Analytical RP-HPLC chromatogram (214 nm) of purified peptide, **A11** (*ca.* 99% as analysed by peak area). Phenomenex Aeris$^{TM}$ 5 µm Peptide XB-C18 column (100Å, 150 × 4.6 mm), linear gradient 5%–95%B over 50 min (*ca.* 1.8%B/min) at 1 mL/min. $t_R$ = 24.4 mins.

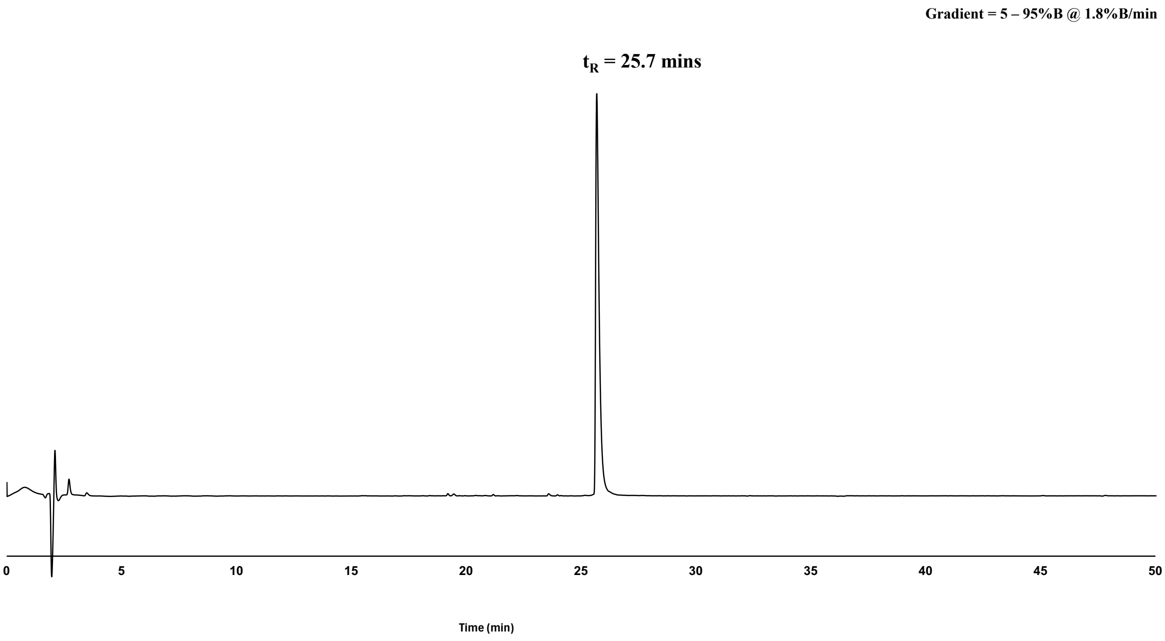


**Supplementary Methods Figure 12.** Analytical RP-HPLC chromatogram (214 nm) of purified peptide, **A12** (*ca.* 96% as analysed by peak area). Phenomenex Aeris™ 5 µm Peptide XB-C18 column (100Å, 150 × 4.6 mm), linear gradient 5%–95%B over 50 min (*ca.* 1.8%B/min) at 1 mL/min. $t_R$ = 25.7 mins.

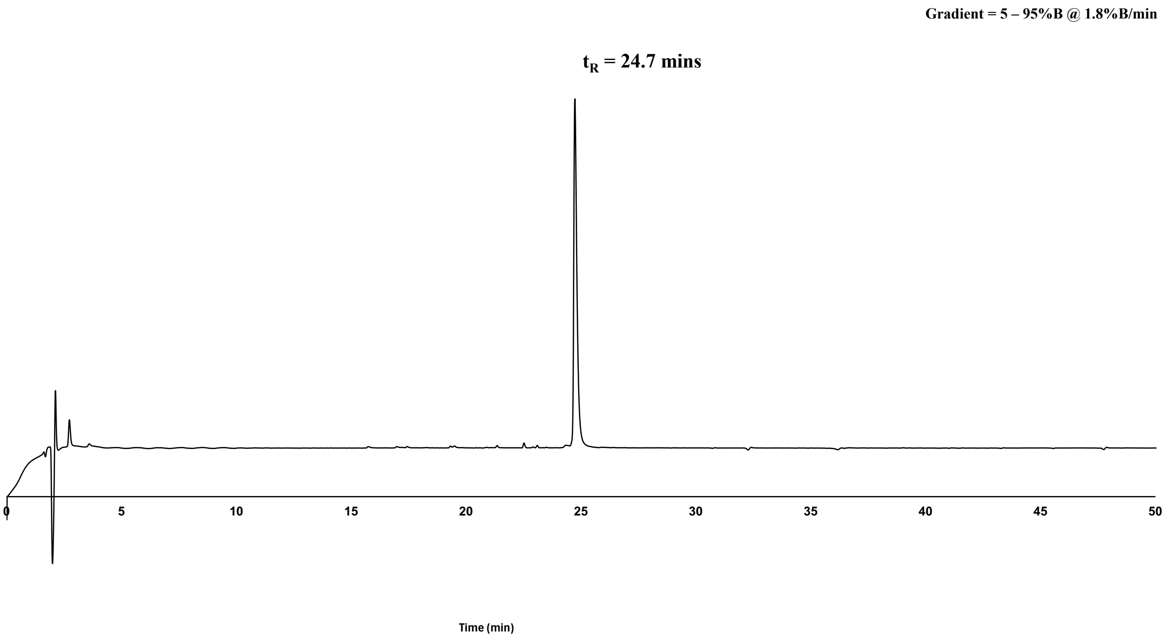


**Supplementary Methods Figure 13.** Analytical RP-HPLC chromatogram (214 nm) of purified peptide, **B1** (*ca.* 98% as analysed by peak area). Phenomenex Aeris™ 5 µm

Peptide XB-C18 column (100Å, 150 × 4.6 mm), linear gradient 5%–95%B over 50 min (*ca.* 1.8%B/min) at 1 mL/min. $t_R$ = 24.7 mins.

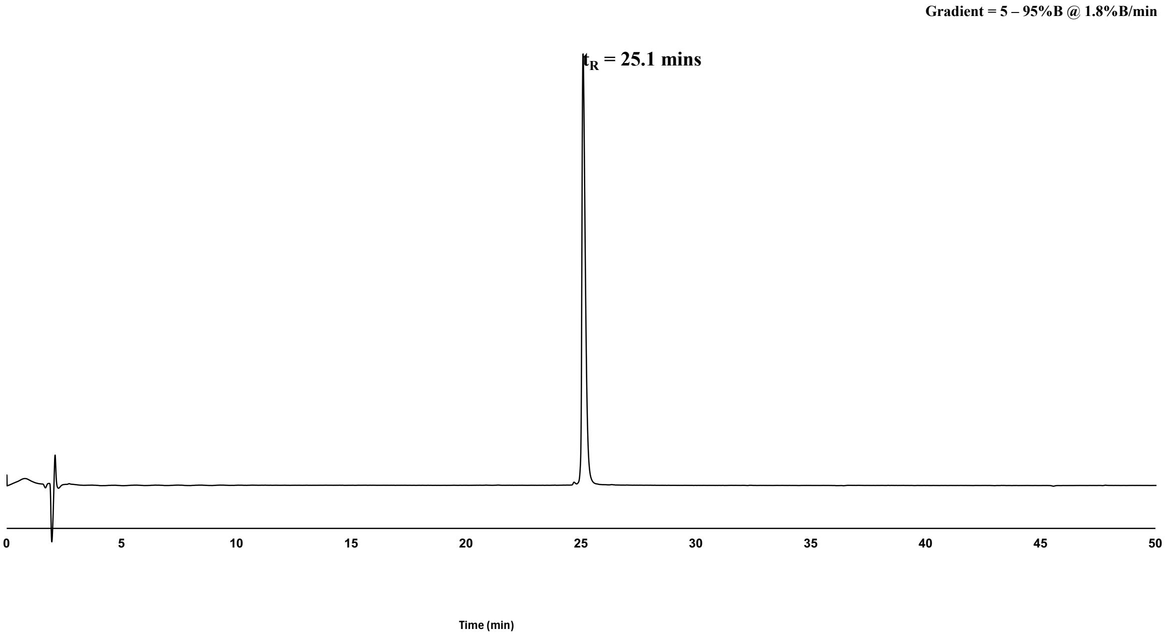


**Supplementary Methods Figure 14.** Analytical RP-HPLC chromatogram (214 nm) of purified peptide, **B2** (*ca.* 99% as analysed by peak area). Phenomenex Aeris™ 5 μm Peptide XB-C18 column (100Å, 150 × 4.6 mm), linear gradient 5%–95%B over 50 min (*ca.* 1.8%B/min) at 1 mL/min. $t_R$ = 25.1 mins.

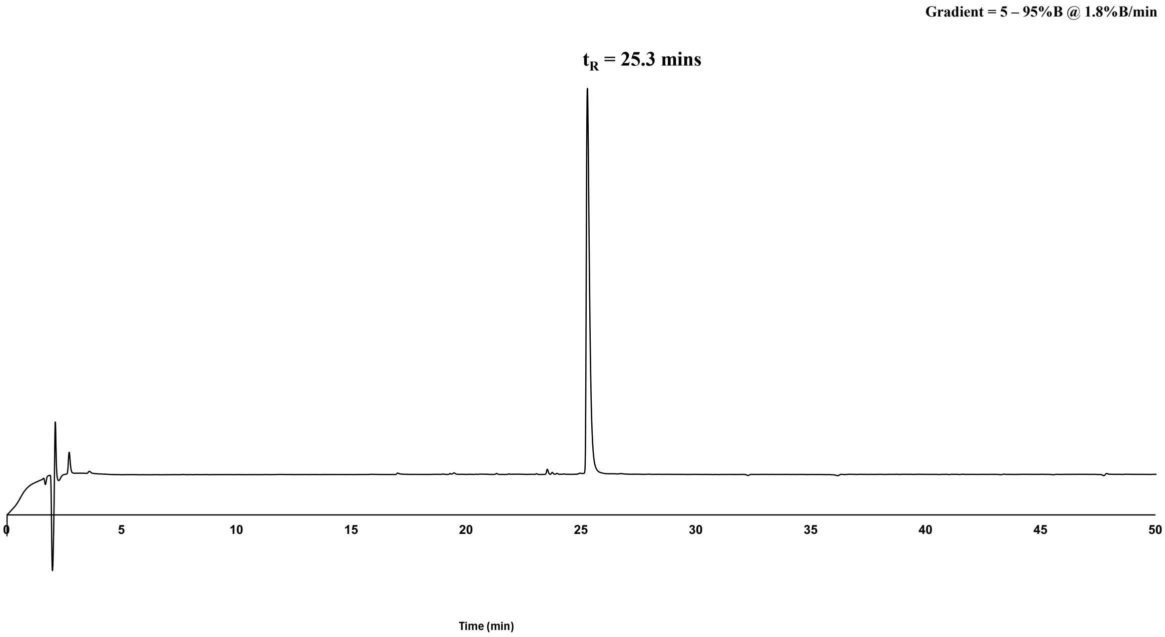

**Supplementary Methods Figure 15.** Analytical RP-HPLC chromatogram (214 nm) of purified peptide, **B3** (*ca.* 98% as analysed by peak area). Phenomenex Aeris™ 5 μm Peptide XB-C18 column (100Å, 150 × 4.6 mm), linear gradient 5%–95%B over 50 min (*ca.* 1.8%B/min) at 1 mL/min. $t_R$ = 25.3 mins.

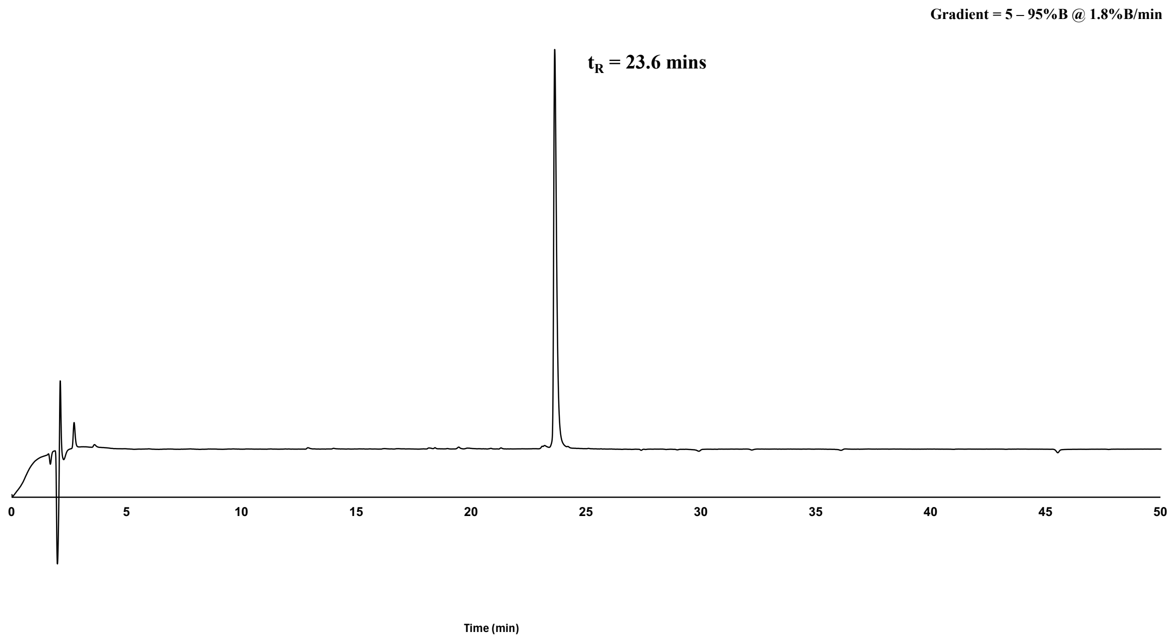


**Supplementary Methods Figure 16.** Analytical RP-HPLC chromatogram (214 nm) of purified peptide, **B4** (*ca.* 96% as analysed by peak area). Phenomenex Aeris™ 5 μm Peptide XB-C18 column (100Å, 150 × 4.6 mm), linear gradient 5%–95%B over 50 min (*ca.* 1.8%B/min) at 1 mL/min. $t_R$ = 23.6 mins.

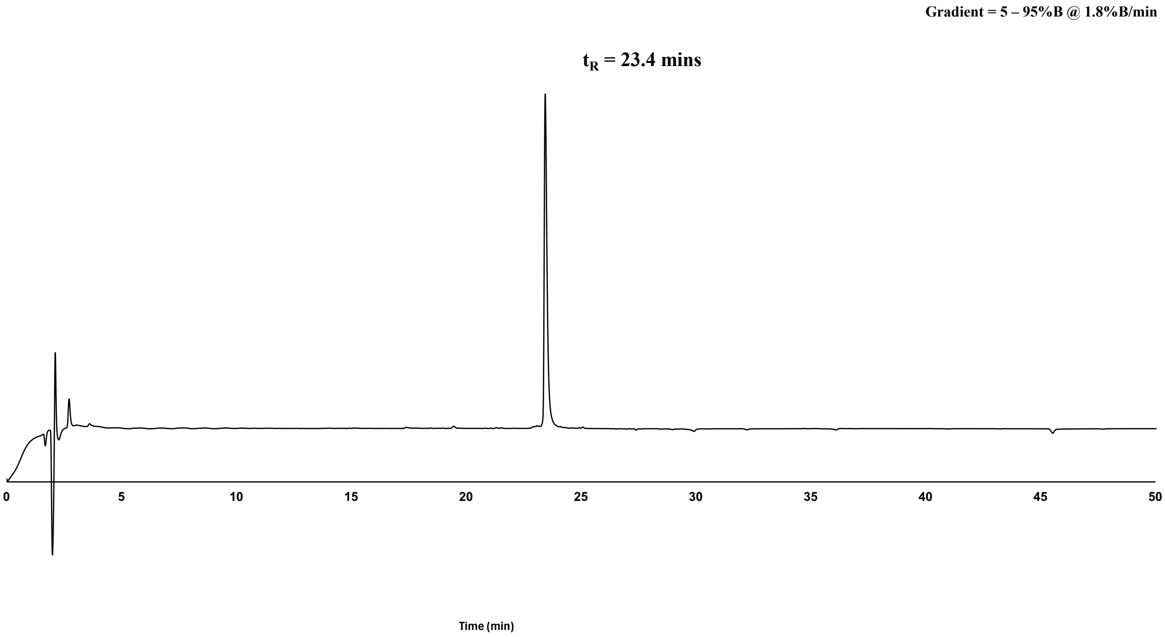


**Supplementary Methods Figure 17.** Analytical RP-HPLC chromatogram (214 nm) of purified peptide, **B5** (*ca.* 97% as analysed by peak area). Phenomenex Aeris™ 5 µm Peptide XB-C18 column (100Å, 150 × 4.6 mm), linear gradient 5%–95%B over 50 min (*ca.* 1.8%B/min) at 1 mL/min. $t_R$ = 23.4 mins.

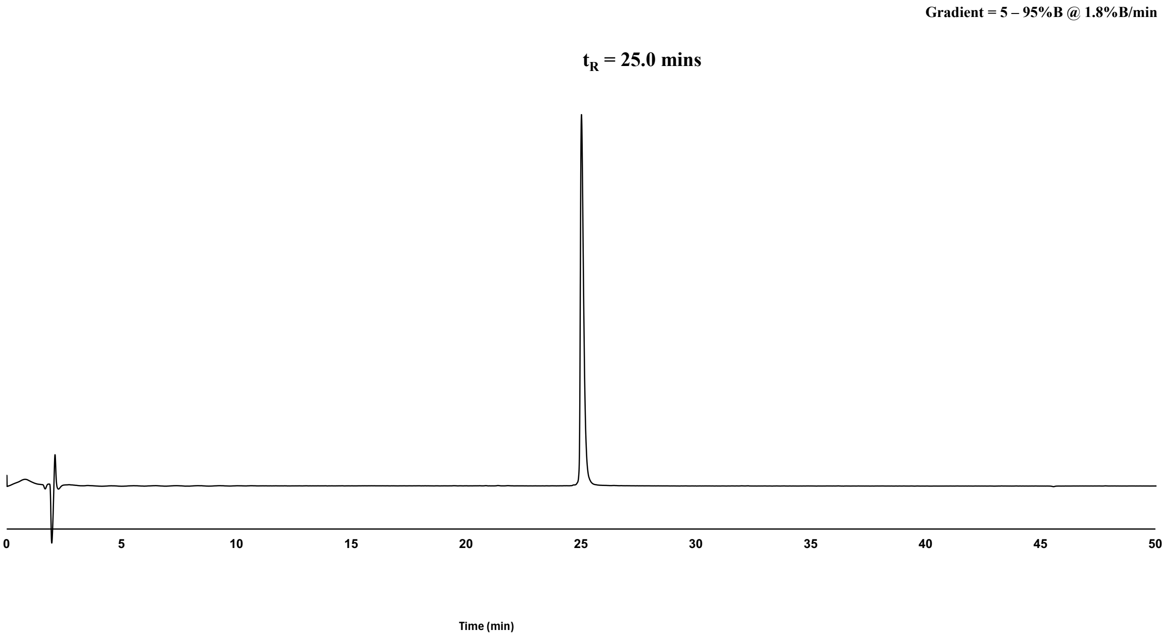


**Supplementary Methods Figure 18.** Analytical RP-HPLC chromatogram (214 nm) of purified peptide, **B6** (*ca.* 98% as analysed by peak area). Phenomenex Aeris™ 5 µm Peptide XB-C18 column (100Å, 150 × 4.6 mm), linear gradient 5%–95%B over 50 min (*ca.* 1.8%B/min) at 1 mL/min. $t_R$ = 25.0 mins.

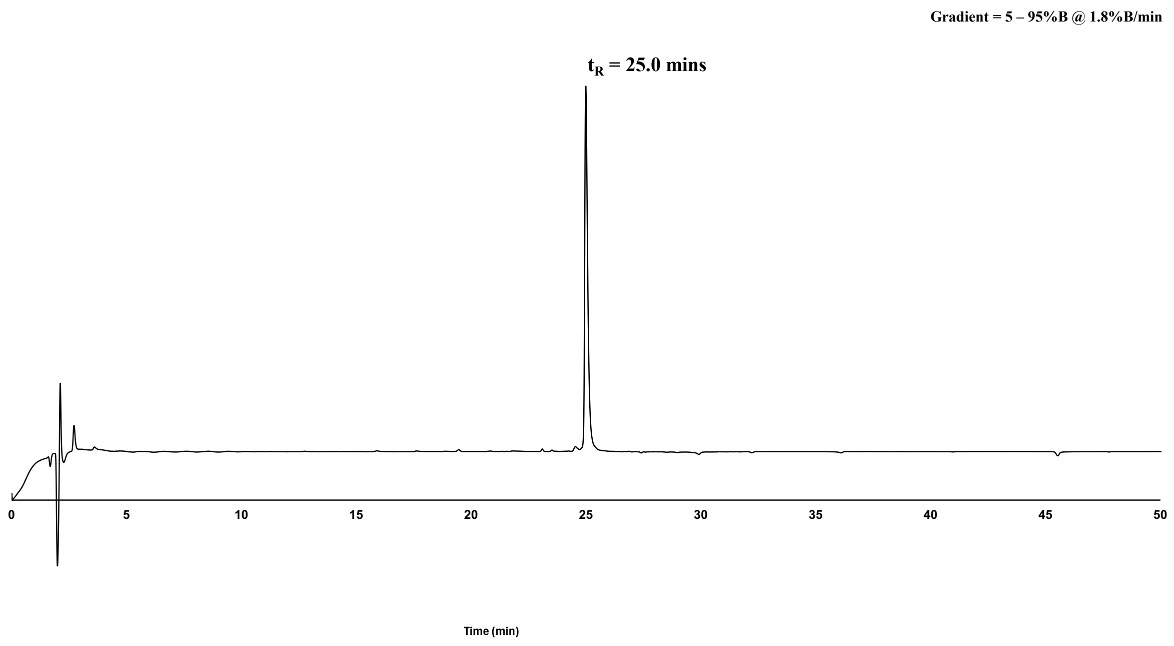


**Supplementary Methods Figure 19.** Analytical RP-HPLC chromatogram (214 nm) of purified peptide, **B7** (*ca.* 96% as analysed by peak area). Phenomenex Aeris™ 5 µm Peptide XB-C18 column (100Å, 150 × 4.6 mm), linear gradient 5%–95%B over 50 min (*ca.* 1.8%B/min) at 1 mL/min. $t_R$ = 25.0 mins.

**Liquid Chromatography Mass Spectrometry (LCMS)**

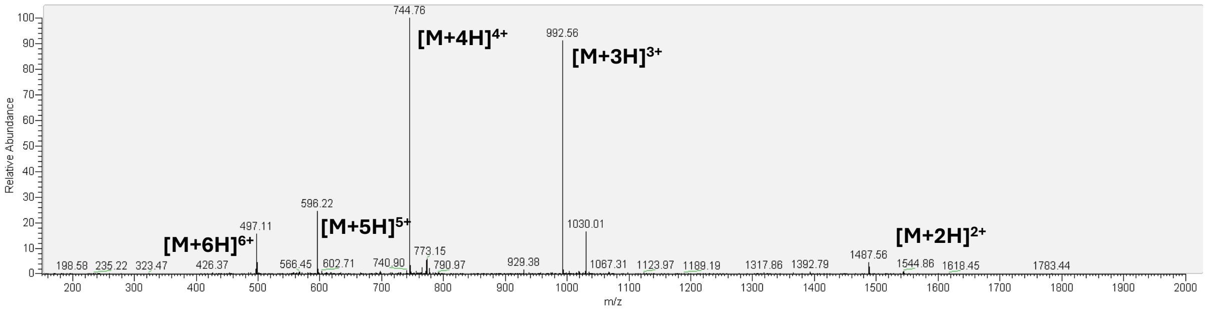


**Supplementary Methods Figure 20.** LCMS of purified peptide, **A1**, mass calculated for [$C_{136}H_{217}N_{39}O_{36}$ + H] 2974.47; deconvoluted mass observed: 2975.12 ± 1.37. Charge states; 497.11 [M+6H]$^{6+}$, 596.22 [M+5H]$^{5+}$, 744.76 [M+4H]$^{4+}$, 992.56 [M+3H]$^{3+}$, 1478.56 [M+2H]$^{2+}$.

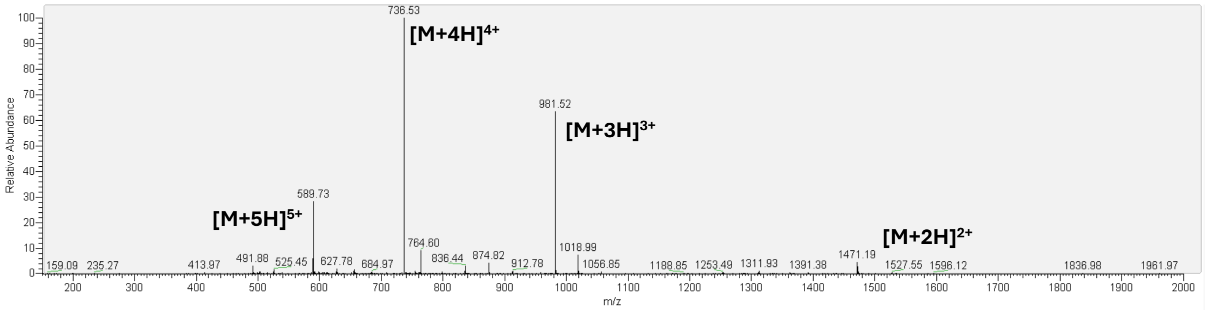


**Supplementary Methods Figure 21.** LCMS of purified peptide, **A2**, mass calculated for [$C_{131}H_{214}N_{40}O_{37}$ + H] 2941.40; deconvoluted mass observed: 2941.93 ± 1.36. Charge states; 589.73 [M+5H]$^{5+}$, 736.53 [M+4H]$^{4+}$, 981.52 [M+3H]$^{3+}$, 1471.19 [M+2H]$^{2+}$.

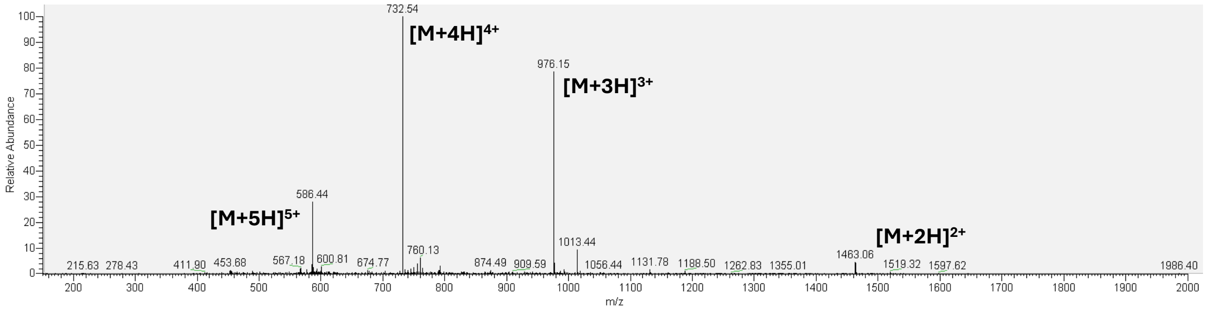


**Supplementary Methods Figure 22.** LCMS of purified peptide, **A3**, mass calculated for [$C_{131}H_{214}N_{40}O_{36}$ + H] 2925.40; deconvoluted mass observed: 2925.74 ± 1.28. Charge states; 586.44 [M+5H]$^{5+}$, 732.54 [M+4H]$^{4+}$, 976.15 [M+3H]$^{3+}$, 1463.06 [M+2H]$^{2+}$.

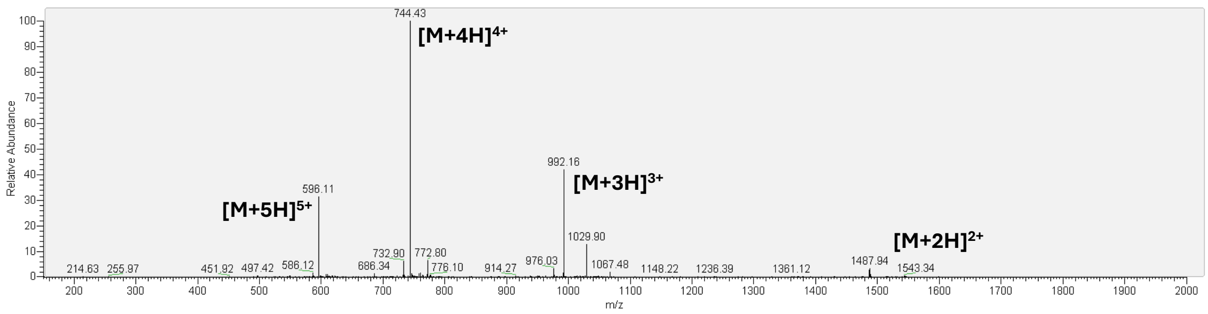


**Supplementary Methods Figure** 23. LCMS of purified peptide, A4, mass calculated for

[$C_{136}H_{218}N_{40}O_{35}$ + H] 2973.48; deconvoluted mass observed: 2974.16 ± 0.94. Charge states; 596.11 $[M+5H]^{5+}$, 744.43 $[M+4H]^{4+}$, 992.16 $[M+3H]^{3+}$, 1487.94 $[M+2H]^{2+}$.

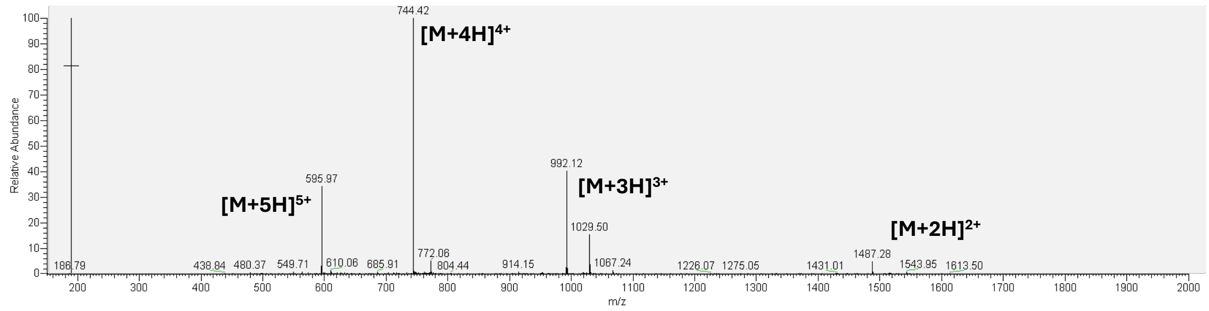


**Supplementary Methods Figure** 24. LCMS of purified peptide, A5, mass calculated for [$C_{136}H_{218}N_{40}O_{35}$ + H] 2973.48; deconvoluted mass observed: 2973.61 ± 0.95. Charge states; 595.97 $[M+5H]^{5+}$, 744.42 $[M+4H]^{4+}$, 992.12 $[M+3H]^{3+}$, 1487.28 $[M+2H]^{2+}$.

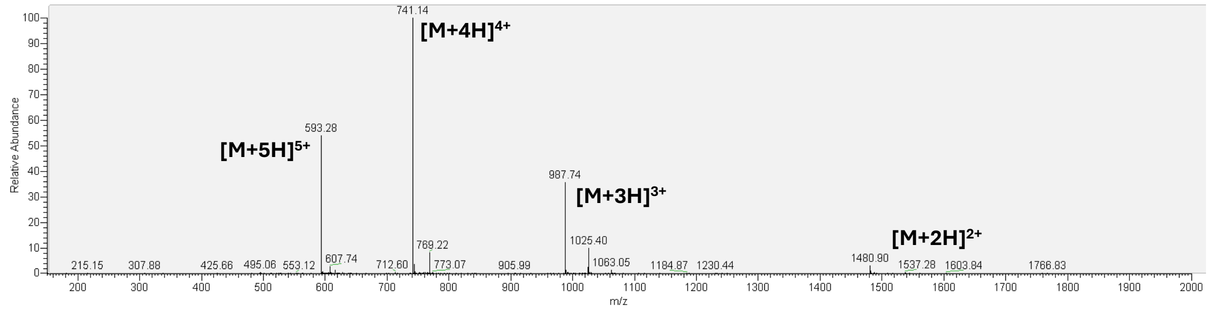


**Supplementary Methods Figure** 25. LCMS of purified peptide, A6, mass calculated for [$C_{135}H_{215}N_{39}O_{36}$ + H] 2960.44; deconvoluted mass observed: 2960.50 ± 0.68. Charge states; 593.28 $[M+5H]^{5+}$, 741.14 $[M+4H]^{4+}$, 987.74 $[M+3H]^{3+}$, 1480.90 $[M+2H]^{2+}$.

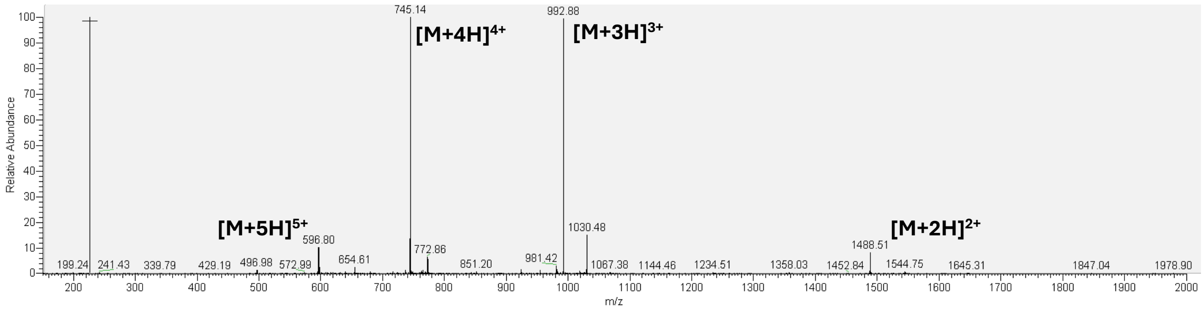


**Supplementary Methods Figure 26.** LCMS of purified peptide, **A7,** mass calculated for [$C_{134}H_{212}N_{40}O_{37}$ + H] 2975.41; deconvoluted mass observed: 2976.56 ± 1.75. Charge states; 596.80 $[M+5H]^{5+}$, 745.14 $[M+4H]^{4+}$, 992.88 $[M+3H]^{3+}$, 1488.51 $[M+2H]^{2+}$.

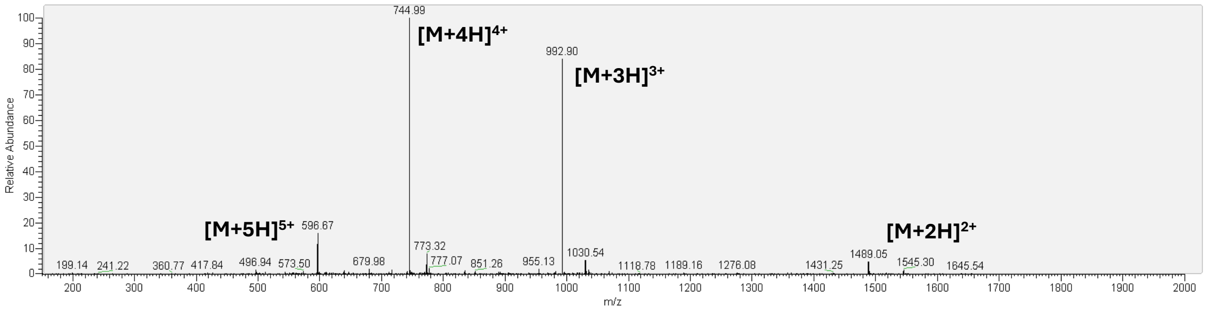


**Supplementary Methods Figure 27.** LCMS of purified peptide, **A8,** mass calculated for [$C_{136}H_{217}N_{39}O_{36}$ + H] 2974.47; deconvoluted mass observed: 2976.53 ± 1.23. Charge states; 596.67 $[M+5H]^{5+}$, 744.9 $[M+4H]^{4+}$, 992.90 $[M+3H]^{3+}$, 1489.05 $[M+2H]^{2+}$.

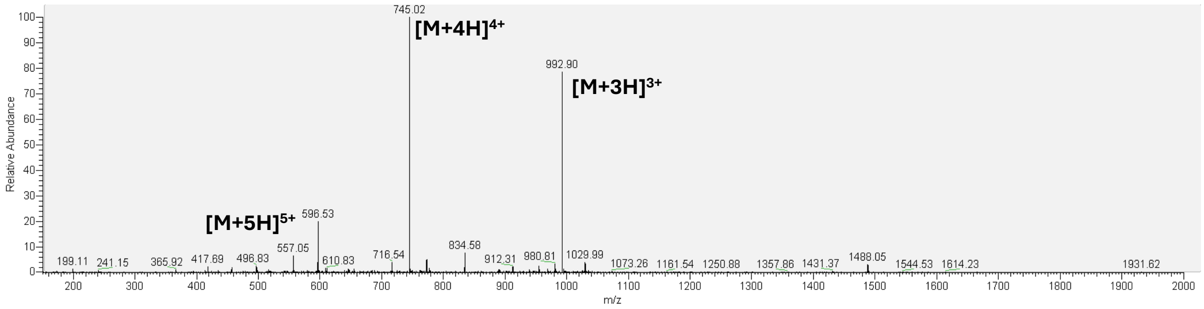


**Supplementary Methods Figure 28.** LCMS of purified peptide, **A9,** mass calculated for [$C_{134}H_{212}N_{40}O_{37}$ + H] 2975.41; deconvoluted mass observed: 2976.48 ± 1.03. Charge states; 596.53 $[M+5H]^{5+}$, 745.02 $[M+4H]^{4+}$, 992.90 $[M+3H]^{3+}$.

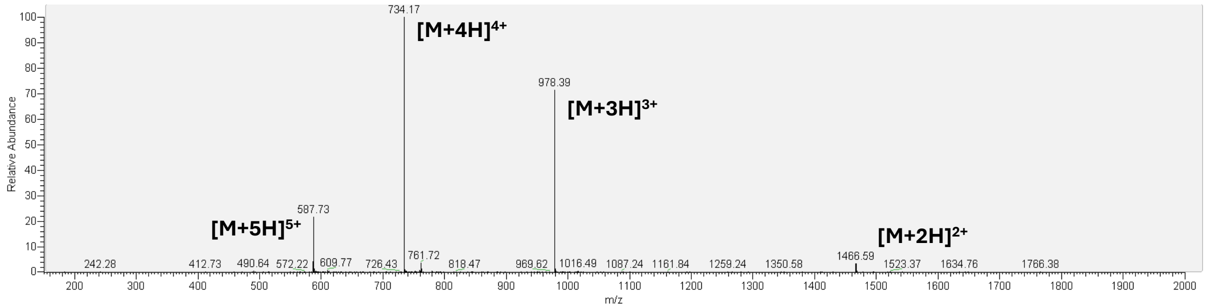


**Supplementary Methods Figure 29.** LCMS of purified peptide, **A10**, mass calculated for [$C_{134}H_{211}N_{37}O_{37}$ + H] 2932.38; deconvoluted mass observed: 2932.42 ± 1.03. Charge states; 587.73 $[M+5H]^{5+}$, 734.17 $[M+4H]^{4+}$, 978.39 $[M+3H]^{3+}$, 1466.59 $[M+2H]^{2+}$.

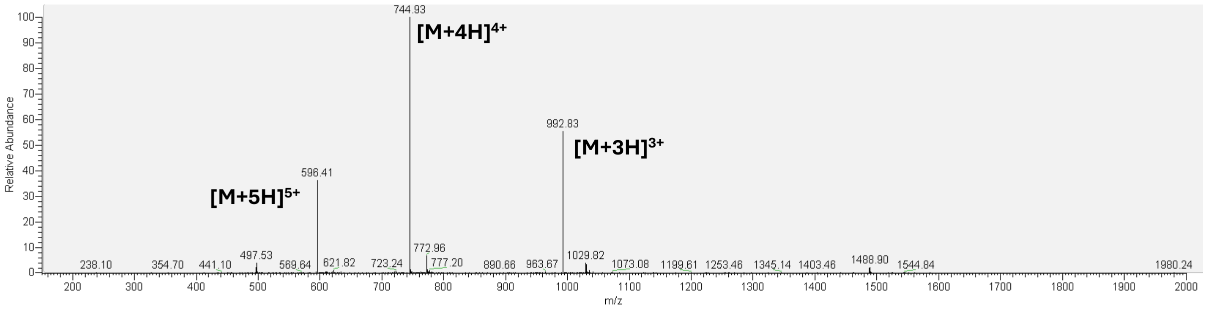


**Supplementary Methods Figure 30.** LCMS of purified peptide, **A11**, mass calculated for [$C_{134}H_{212}N_{40}O_{37}$ + H] 2975.41; deconvoluted mass observed: 2976.09 ± 0.84. Charge states; 596.41 $[M+5H]^{5+}$, 744.93 $[M+4H]^{4+}$, 992.83 $[M+3H]^{3+}$.

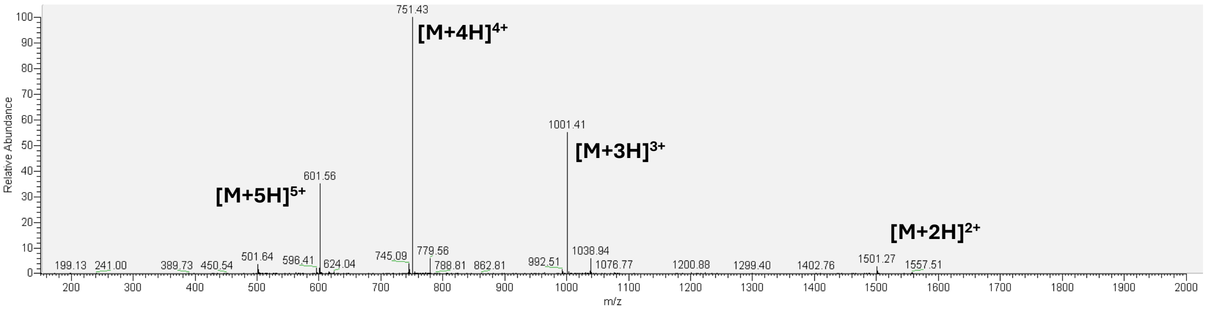


**Supplementary Methods Figure 31.** LCMS of purified peptide, **A12**, mass calculated for

[$C_{137}H_{218}N_{40}O_{36}$ + H] 3001.50; deconvoluted mass observed: 3001.62 ± 0.88. Charge states; 601.56 $[M+5H]^{5+}$, 751.43 $[M+4H]^{4+}$, 1001.41 $[M+3H]^{3+}$, 1501.27 $[M+2H]^{2+}$.

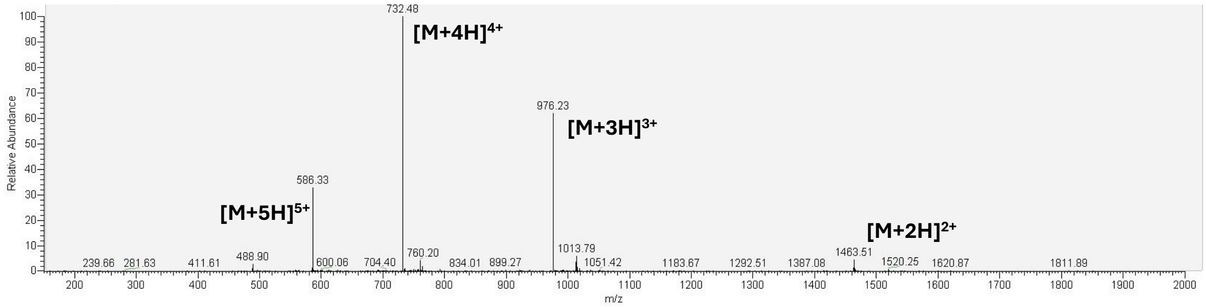


**Supplementary Methods Figure 32.** LCMS of purified peptide, **B1**, mass calculated for [$C_{131}H_{214}N_{40}O_{36}$ + H] 2925.40; deconvoluted mass observed: 2925.82 ± 0.67. Charge states; 586.33 $[M+5H]^{5+}$, 732.48 $[M+4H]^{4+}$, 976.23 $[M+3H]^{3+}$, 1463.51 $[M+2H]^{2+}$.

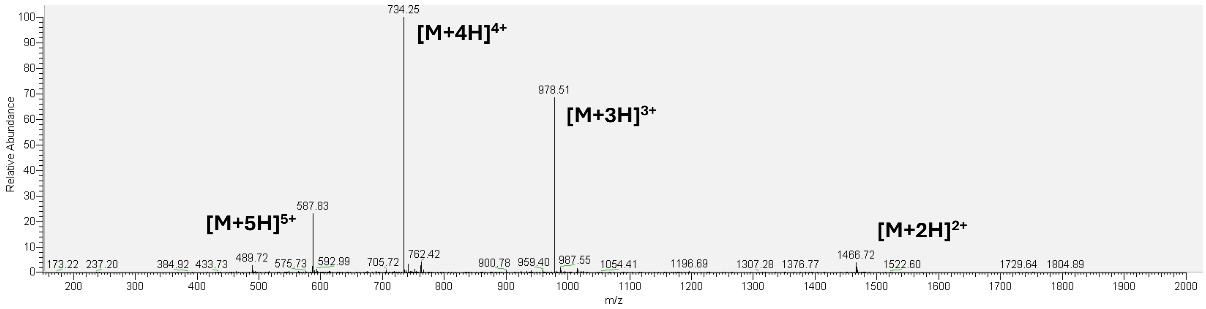


**Supplementary Methods Figure 33.** LCMS of purified peptide, **B2**, mass calculated for [$C_{134}H_{211}N_{37}O_{37}$ + H] 2932.38; deconvoluted mass observed: 2932.78 ± 1.12. Charge states; 587.83 $[M+5H]^{5+}$, 734.25 $[M+4H]^{4+}$, 978.51 $[M+3H]^{3+}$, 1466.72 $[M+2H]^{2+}$.

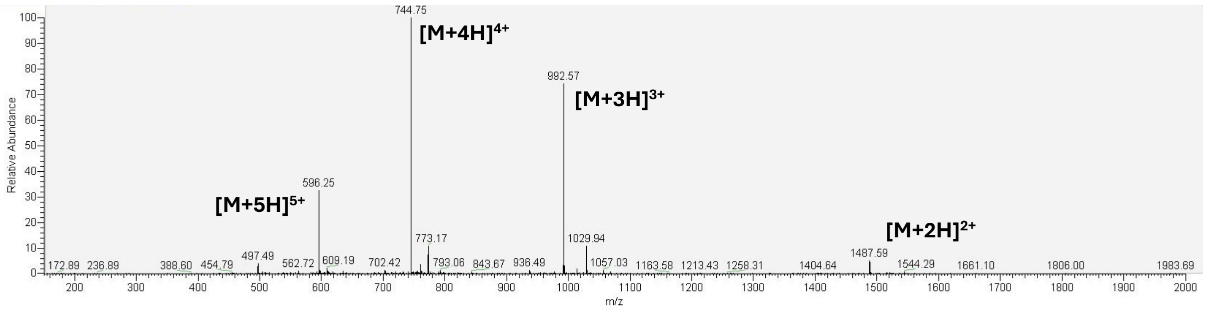


**Supplementary Methods Figure 34.** LCMS of purified peptide, **B3**, mass calculated for [$C_{136}H_{217}N_{39}O_{36}$ + H] 2974.47; deconvoluted mass observed: 2974.79 ± 1.26. Charge states; 596.25 $[M+5H]^{5+}$, 744.75 $[M+4H]^{4+}$, 992.57 $[M+3H]^{3+}$, 1487.59 $[M+2H]^{2+}$.

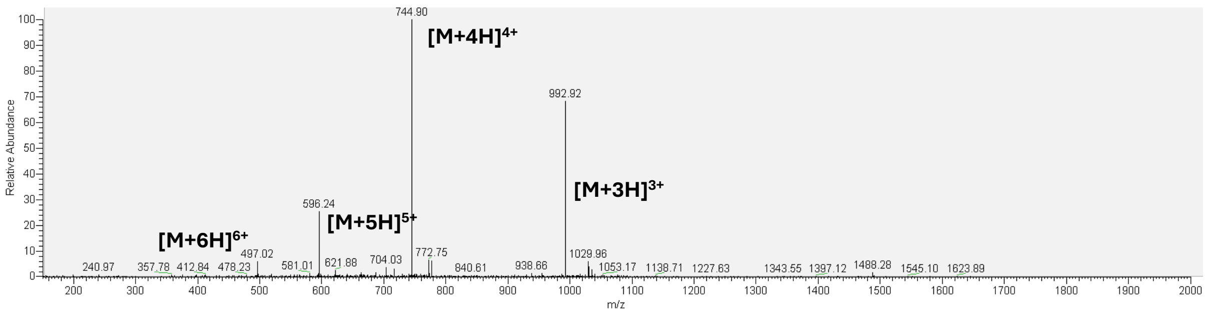


**Supplementary Methods Figure** **35.** LCMS of purified peptide, **B4,** mass calculated for [$C_{134}H_{212}N_{40}O_{37}$ + H] 2975.41; deconvoluted mass observed: 2975.92 ± 0.29. Charge states; 496.02 [M+6H]$^{6+}$, 596.24 [M+5H]$^{5+}$, 744.90 [M+4H]$^{4+}$, 992.92 [M+3H]$^{3+}$.

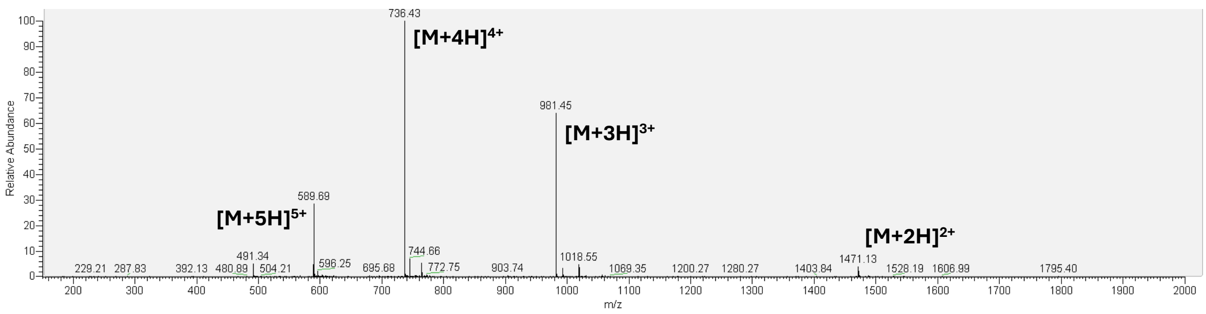


**Supplementary Methods Figure** **36.** LCMS of purified peptide, **B5,** mass calculated for [$C_{131}H_{214}N_{40}O_{37}$ + H] 2941.40; deconvoluted mass observed: 2941.70 ± 1.32. Charge states; 589.69 [M+5H]$^{5+}$, 736.43 [M+4H]$^{4+}$, 981.45 [M+3H]$^{3+}$, 1471.13 [M+2H]$^{2+}$.

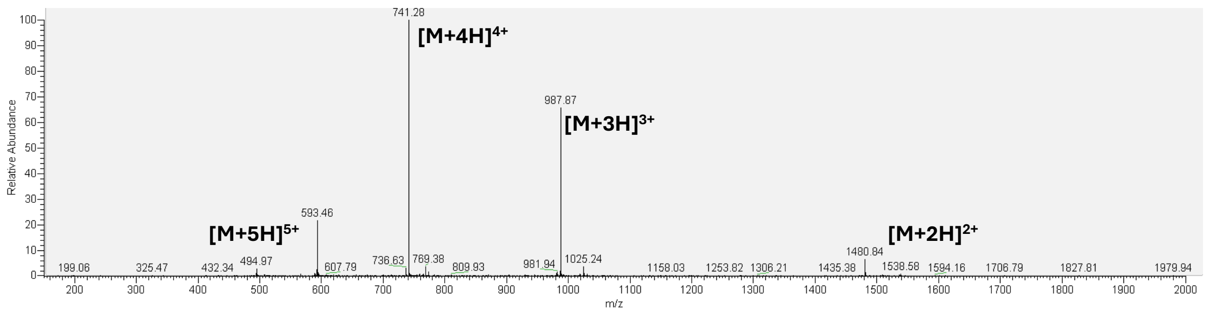


**Supplementary Methods Figure** **37.** LCMS of purified peptide, **B6,** mass calculated for [$C_{134}H_{211}N_{39}O_{37}$ + H] 2960.40; deconvoluted mass observed: 2960.95 ± 1.13. Charge states; 593.46 [M+5H]$^{5+}$, 741.28 [M+4H]$^{4+}$, 987.87 [M+3H]$^{3+}$, 1480.64 [M+2H]$^{2+}$.

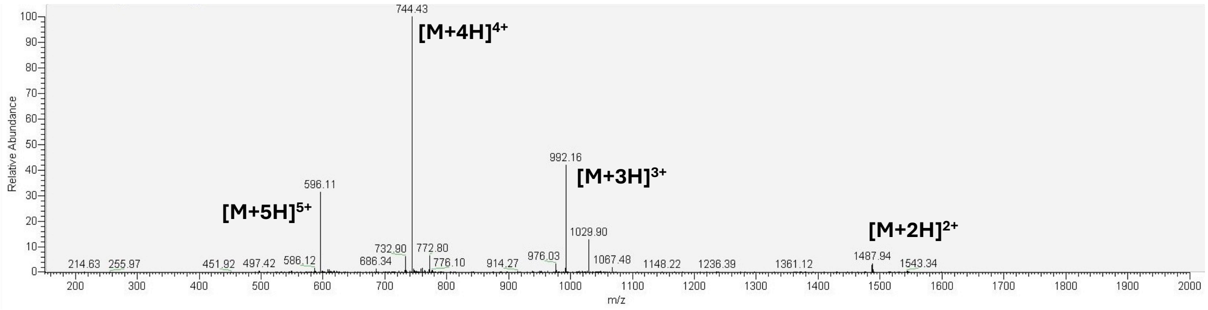


**Supplementary Methods Figure 38.** LCMS of purified peptide, **B7**, mass calculated for [$C_{136}H_{217}N_{39}O_{36}$ + H] 2974.47; deconvoluted mass observed: 2974.16 ± 0.94. Charge states; 598.11 $[M+5H]^{5+}$, 744.43 $[M+4H]^{4+}$, 992.16 $[M+3H]^{3+}$, 1487.94 $[M+2H]^{2+}$.
